# Identification of a specialized lipid barrier for *Drosophila* metamorphosis

**DOI:** 10.1101/2024.10.11.617838

**Authors:** Lena Lampe, Clare L. Newell, Bing-Jun Wang, Rami Makki, Cyrille Alexandre, Ian S. Gilmore, Li Zhao, Alex P. Gould

## Abstract

In many terrestrial insects, the onset of metamorphosis marks a transition from humid to dry environments. Yet how metamorphosing insect pupae protect themselves against the threat of dehydration remains unclear. Here, we identify the chemical composition and biosynthetic origins of a lipid desiccation barrier specific to the pupal and sexually-immature adult stages of *Drosophila melanogaster*. This barrier comprises unisex hyper-long hydrocarbons, 29-37 carbons in length, which are synthesized by larval oenocytes and stored in the larval fat body before being deployed on the pupal and young adult cuticles. We show that the fatty acid elongase *EloHL* is required for the biosynthesis of hyper-long hydrocarbons that are essential for the barrier to water loss during metamorphosis. Across the *Drosophila* genus, many species express unisex profiles of hyper-long hydrocarbons and, as young adults, transition to sex-specific shorter hydrocarbons with known pheromonal functions. The desert species *D. mojavensis*, however, retains hyper-long hydrocarbons during adulthood likely as an adaptation to an arid environment. Our study reveals how the cuticular lipid barrier is tuned to meet changing environmental pressures during insect development and evolution.

## INTRODUCTION

Dehydration is a major threat to all terrestrial life. The evolution of multiple water conservation mechanisms has nevertheless allowed animals to colonize much of the earth’s land surface, including hot and arid zones. [1, 2] One critical adaptation for restricting water loss is the cutaneous lipid barrier - a hydrophobic coating of the integument that provides a direct interface between the animal and its environment. This widespread if not universal feature of terrestrial fauna is particularly important for small animals, like insects, as their large surface area to volume ratio makes them highly vulnerable to desiccation. [3-5] Insects have therefore evolved very effective cuticular lipid barriers to water loss, contributing to their remarkable success as the most diverse and species-rich class of animals. [6-11] The superficial site and chemistry of insect cuticular lipids make them well suited to perform an additional function as semi-volatile pheromones, conveying social cues about location and physiological status. [12-14] The potential dual role of some cuticular lipid species - as both desiccation barrier and sex pheromone - raises intriguing evolutionary questions about the links between ecological adaptation and speciation. [15-18]

Holometabolous insects undergo complete metamorphosis. This remarkable and complex developmental process involves a dramatic morphological and physiological transformation of the larva into the adult (imago) and is often accompanied by a shift in ecological niche. In many holometabolous insects, including the fruit fly *Drosophila melanogaster*, larval stages typically burrow in a semi-aquatic environment of rotting fruit but subsequent pupal and adult stages take place in a terrestrial environment that necessitates a more efficient barrier to desiccation. [19, 20] In adult insects, it is well established that their cuticle is coated with a lipid layer consisting primarily of hydrocarbons. [21-23] These hydrocarbons play key roles both in desiccation resistance and in social communication, especially mating behaviour. [16, 24-26] *Drosophila* cuticular hydrocarbons fall into four main categories (*n*-alkanes, methyl-branched alkanes, monoenes and dienes) and the precise blend of these determines the overall biophysical properties of the cuticular hydrocarbon layer, such as melting temperature, transpiration rate and volatility. [27-29] Adult oenocytes are the site of synthesis of cuticular hydrocarbons in pterygote insects.[30-34] In *D. melanogaster* and many other insects, these specialized lipid-secreting cells are confined to abdominal segments, where they are located beneath the epidermis and cross-talk with other tissues to coordinate lipid metabolism. [35-38] Adult oenocytes synthesize hydrocarbons from fatty acyl-CoA precursors via reduction to fatty aldehydes and loss of one carbon by cytochrome-P450 dependent oxidative decarbonylation [39, 40]. In *Drosophila*, knockout or oenocyte-specific knockdown of the P450 decarbonylase, Cyp4g1, results in loss of cuticular hydrocarbons, susceptibility to desiccation, and reduced viability upon adult emergence (eclosion). [40, 41] Multiple fatty acid synthases, elongases, and desaturases expressed in adult oenocytes regulate the types of fatty acyl-CoA substrates for Cyp4g1, in turn determining the length and desaturation of molecules in the complex blend of cuticular hydrocarbons. [42-47]

Across the *Drosophila* genus, adult cuticular hydrocarbon profiles are highly divergent and tend to be sex-specific.[48] Natural selection is thought to favour the production of long-chain hydrocarbons with low volatility as these are thought to contribute most to an effective adult desiccation barrier. [29, 49-51] Consistent with this, carbon chain length correlates across species of *Drosophila*, and also ants, with the temperature and humidity of their habitats. [52-54] In particular, there is a strong correlation between the evolution of longer chain methyl-branched hydrocarbons and higher desiccation resistance across the *Drosophila* genus.[51] Furthermore, in the desert species *D. mojavensis,* evolutionary changes in an oenocyte fatty acyl–CoA elongase (*mElo*) confer chain-lengthening of methyl-branched hydrocarbons and increased desiccation resistance.[43] The pheromonal role of adult hydrocarbons also places them under strong sexual selection, with both males and females exhibiting a mate preference for particular blends. [55-58] In *D. melanogaster*, several hydrocarbon molecules have been identified as sex and species-specific pheromones. [24, 25, 59] For example, female dienes of 27 or 29 carbons in length (C27:2 and C29:2) are synthesized by a sex-specific oenocyte fatty acid elongase (*EloF*) and desaturase (*desatF*) required for mating success. [45-47] In summary, the available evidence suggests that the evolution of adult cuticular hydrocarbons is shaped by natural selection, promoting non-volatile n-alkanes and methyl-branched alkenes for desiccation resistance, and sexual selection favouring semi-volatile monoenes and dienes for pheromonal communication.

In contrast to the barrier lipids of the adult fly, the cellular origin and molecular composition of their counterparts in the pupa remain mysterious. This raises fundamental questions about how this terrestrial phase of development is protected from desiccation. Here we use a combination of genetics, mass spectrometry and a surface-specific analytical technique to identify the molecular composition of the pupal desiccation barrier as well as its cellular origin and biosynthetic pathway.

## RESULTS

### Sexually immature flies express unisex hyper-long cuticular hydrocarbons

To characterise hydrocarbon (HC) profiles during *D. melanogaster* development, we optimised a high temperature gas-chromatography mass-spectrometry (HT-GC-MS) workflow capable of resolving chain lengths of up to ∼40 carbons (C40, see Methods). Hexane extracts of the body surface analysed using this method did not contain detectable amounts of HCs at the wandering third-instar larval (wL3) or pupal (48h after pupariation) stages. In contrast, at 0-6 hr after adult emergence (eclosion), young flies express abundant HCs on their cuticle (**Figure 1A and Table S1**). The male and female cuticular hydrocarbon profiles of 0-6 hr flies are remarkably similar, contrasting with marked sex-specific differences in older flies at 1, 3 and 7 days after eclosion (**Figures 1A + S1, Table S1**). Principal component analysis (PCA) illustrates that the transition from unisex to male/female specific cuticular profiles is already evident at 1 day and very marked by 3 days (**Figure 1B**). Importantly, the range of cuticular hydrocarbon chain lengths switches from C27-C37 in young flies to C21-C31 in older flies, with an intermediate pattern at 1-3 days of age (**Figures 1C and S1**). Henceforth, we define C21 to C28 hydrocarbons as long (L) and C29 to C37 hydrocarbons as hyper-long (HL). The major categories of cuticular HL hydrocarbons are alkenes, dienes and methyl-branched (mb) alkanes, with only low levels of linear alkanes. Interestingly, although dienes are female-restricted in mature adults, we observed HL dienes in both sexes at 0-6 hr after eclosion (**Figure 1A**). During the first week after adult eclosion, both sexes transition to a lower cuticular ratio of HL to L hydrocarbons but females maintain a higher value, primarily because they continue to express C29-C31 hydrocarbons (**Figures 1A and 1C**). Together, the HT-GC-MS analysis shows that the external body surfaces of larvae and pupae lack detectable hydrocarbons, whereas those of adult flies transition from unisex HL to sex-specific L profiles within a day of eclosion.

**Figure 1.**
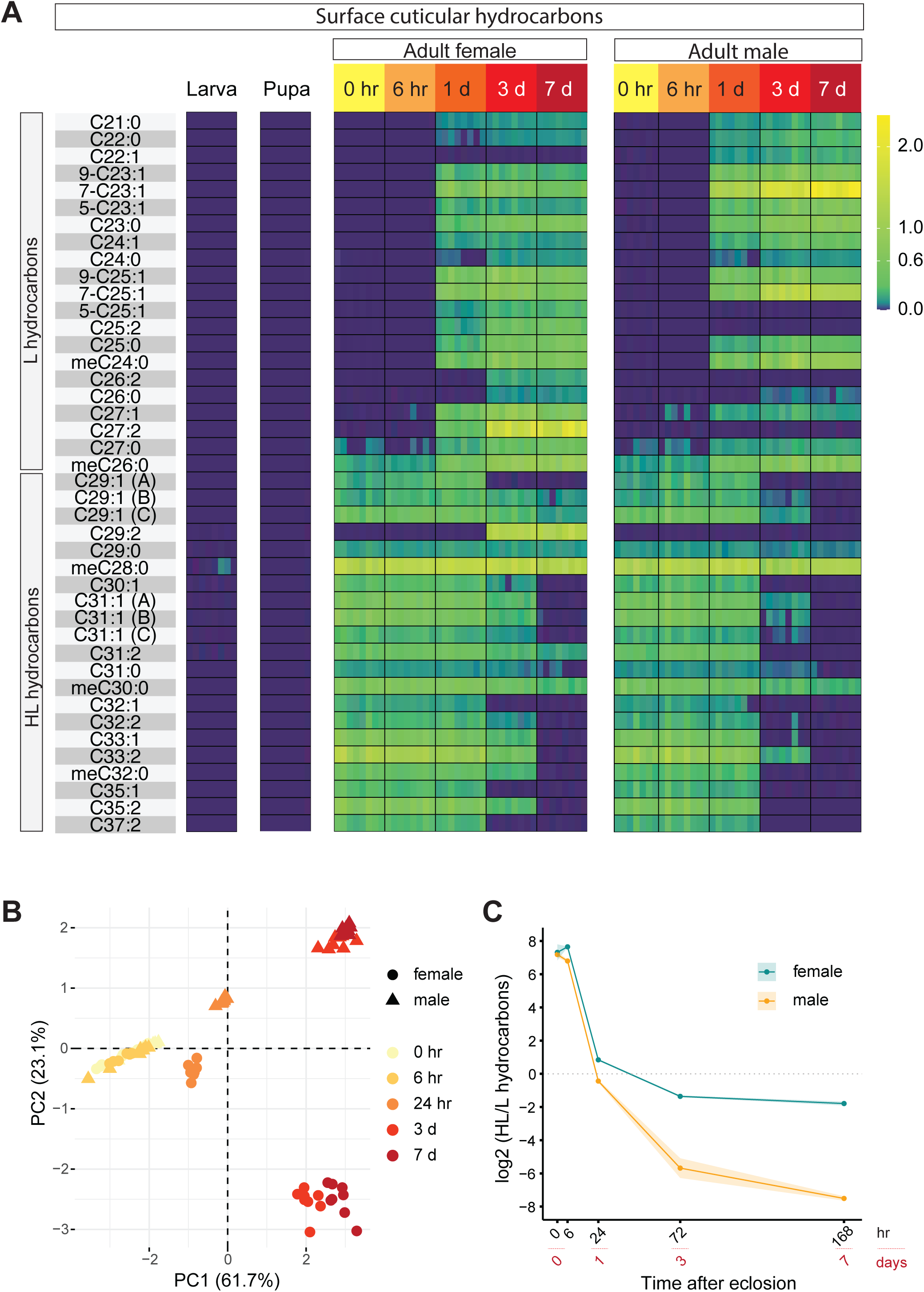
Surface cuticular hydrocarbons transition from unisex hyper-long (HL) to sex-specific profiles within a day of adult eclosion. (A) Heatmap of cuticular hydrocarbons of the larva (at wandering L3 stage), pupa (at 48h after puparium formation), and male and female adults (at 0 hr, 6 hr, 24 hr, 3 days and 7 days after eclosion). Hydrocarbons with a carbon length between C21-27 are defined as hydrocarbons (HCs) and C29-31 as hyper-long HCs (HL-HCs). A transition from unisex hyper-long (HL-HCs) to long sex-specific (L-HCs) hydrocarbon profiles occurs between 6-24 hr after adult eclosion. Replicates for each time point are plotted separately next to each other. Data is shown in **Table S1.** (B) Principal component analysis of cuticular HCs from adult males (triangles) and females (circles) at 0 to 7 days of age. (C) C29-C37 (HL-HCs) to C21-C27 (HCs) cuticular hydrocarbon ratios of male and female adult flies at 0 to 7 days after eclosion. The HL-HC/L-HC ratio decreases less for females than males as this sex maintains a subset of HL-HCs after sexual maturation. meC28 was excluded from the ratio analysis due to its robust high abundance across all time points. In panels B-D, the heatmap, PCA and HL-HC/L-HC ratio were determined from four replicates from each of two independent experiments (n=4, N=2) and shading either side of the mean represents SD.

### Hyper-long hydrocarbons are synthesized in larval oenocytes and stored in the fat body

HL hydrocarbons are already present on the cuticle surface at the time of adult eclosion, prompting us to investigate their developmental origin. HT-GC-MS analysis of hydrocarbons extracted from homogenized whole animals after external hydrocarbons were removed, revealed the presence of substantial amounts of "internal" HL hydrocarbons, and some L hydrocarbons (meC26 and C27:1), in wandering L3 larvae, even though "external" hydrocarbons are not detectable on the cuticle at this developmental stage (**Figure 2A**). These internal hydrocarbons have a similar, although not identical, composition to those of newly eclosed adults, suggesting that they may originate from a common larval source (**Figure 2A and Table S2**). To identify the larval source of hydrocarbons, we analysed individual dissected tissues. This revealed the presence of substantial amounts of all HL hydrocarbon species in the larval fat body, whereas in the larval carcass (cuticle, epidermis, muscle and oenocytes) and gut, only methyl-branched (mb) hydrocarbons were detected (**Figure 2B and Table S2**). Nevertheless, the levels of two major mb hydrocarbons (mbC28:0 and mbC30:0) were higher in the fat body than in the carcass or gut. These tissue-specific analyses indicate that HL hydrocarbons are stored during development, primarily in the fat body of the larva.

**Figure 2.**
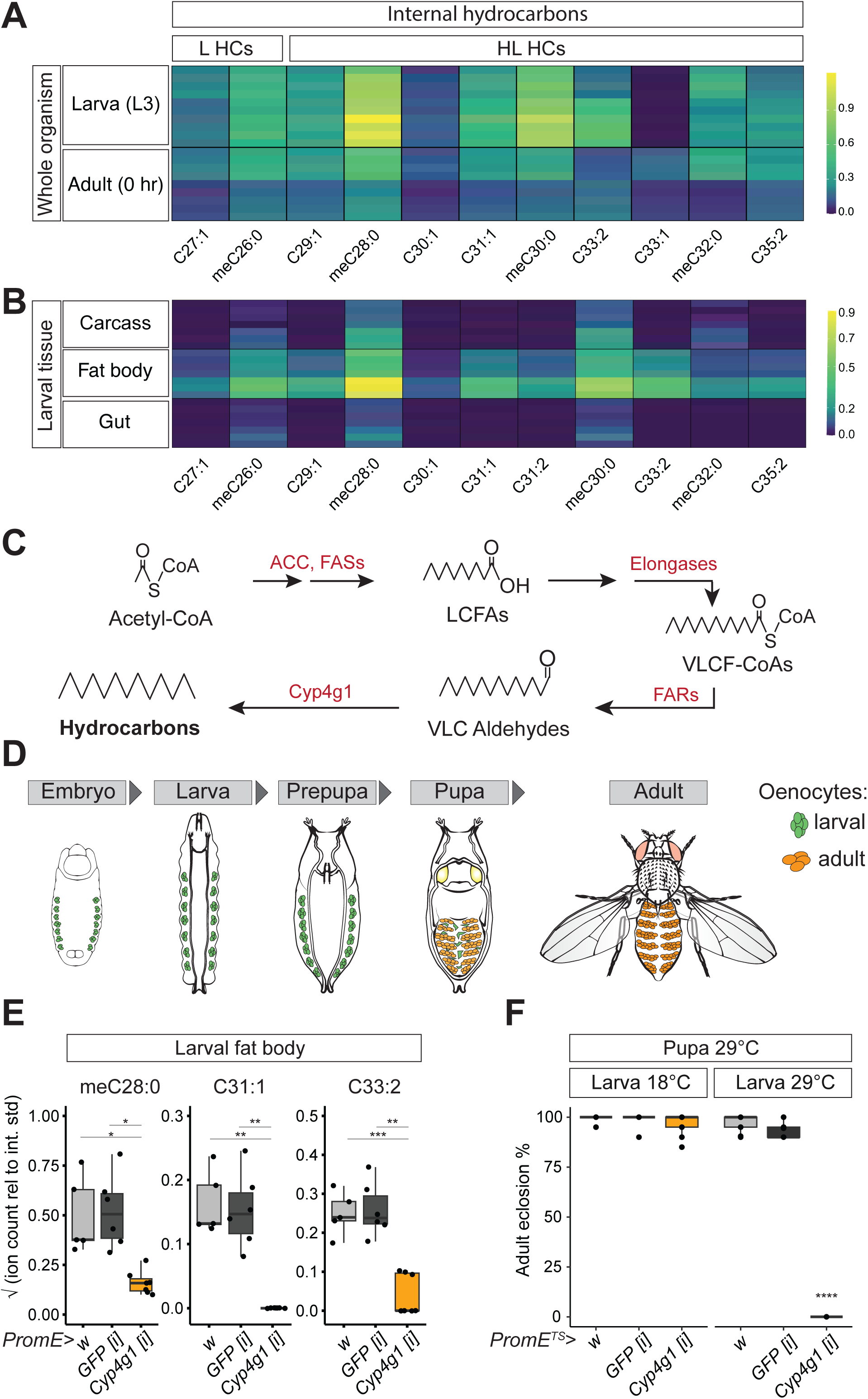
Hyper-long hydrocarbons are synthesized in larval oenocytes and stored in larval fat body. (A) Heatmap of internal hydrocarbons extracted from the wandering third instar larvae (L3 larva) or newly eclosed adult (0 hr Adult). Both the L3 larvae and the newly eclosed adult contain similar internal HL-HCs. Shown are two independent experiments with 4 repeats (n=4, N=2). Additional independent experiments are shown in **Table S2**. (B) Heatmap of internal hydrocarbons extracted from individual larval tissues of wandering L3 larvae (carcass, fat body and gut). Carcass samples include integument, oenocytes, epidermis and muscles and gut include the attached malpighian tubules and crop. (n=3 replicates, N=2 independent experiments). Additional independent experiments are shown in **Table S3**. (C) Hydrocarbon biosynthetic pathway in *Drosophila*. Acetyl-CoA carboxylase (ACC) and fatty acid synthases (FASs) generate long-chain fatty acids (LCFAs) from acetyl-CoA. Fatty acid elongases then convert LCFAs to very long-chain fatty acyl-CoA esters (VLCF-CoAs). Fatty acid reductases (FARs) reduce VLCF-CoAs to very long-chain fatty aldehydes (VLCF aldehydes), that are converted to hydrocarbons via Cyp4g1, a P450 oxidative decarbonylase. (D) Schematic of *Drosophila* development, highlighting the distinct larval (green) and adult (orange) populations of oenocytes. The embryo, larva, prepupa, pupa and adult are shown. Larval oenocytes undergo fragmentation at the mid-pupal stage and do not persist in adults of either sex. Sexual maturation takes place within 24 hr of adult eclosion and is associated with cuticle hardening and an increase in mating frequency. (E) Cyp4g1 is required in larval oenocytes for larval fat body HL-HCs. The y axis shows square root relative quantities of a HL methyl-branched alkane (me-C28:0), a HL monoene (C33:1) and a HL diene (C33:2). The experimental and two control genotypes are *PromE-GAL4* crossed to *UAS-Cyp4g1 RNAi* (*PromE>Cyp4g1[i]*), *UAS-GFP RNAi* (*PromE>GFP[i]*) or to *w[1118] iso31* (*PromE>w*) respectively. Statistical significance is indicated by *(p < 0.05), **(p< 0.05), ***(p<0.0005) using the statistical tests listed in **Table S4**. (n=2 replicates, N=3 independent experiments). (F) *Cyp4g1* is required in larval oenocytes for adult eclosion. PromE^TS^ (*PromE-GAL4, UAS-GAL80^TS^, UAS-CD8::GFP*), active at 29°C but not at 18°C, was crossed to *UAS-Cyp4g1 RNAi* (*PromE^TS^>Cyp4g1[i]*), *UAS-GFP RNAi* (*PromE^TS^>GFP[i]*) or to *w[1118] iso31* (*PromE^TS^>w*). Only expression of *Cyp4g1* RNAi in larval stages affects adult eclosion. *4 refers to p-values lower than 0.00005 (n=3 replicates and N=3 independent experiments) using the statistical tests listed in **Table S4**

We next determined the larval cell type that synthesizes HL hydrocarbons. In adult *Drosophila* and other dipterans, it is known that sex-specific L hydrocarbons are synthesized in specialised lipid-metabolizing cells called adult oenocytes [33, 34, 40]. These hydrocarbons are derived either from dietary or *de novo* synthesized fatty acids (FAs) (**Figure 2C**). In *de novo* synthesis, acetyl-CoA carboxylase (ACC) and fatty acid synthases (FASs) generate long-chain fatty acids (LCFAs) from acetyl-CoA. Fatty acid elongases then convert *de novo* synthesized or dietary LCFAs into very long-chain fatty acyl-CoA esters (VLCF-CoAs) of varying lengths. Fatty acid reductases (FARs) reduce VLCF-CoAs to very long-chain fatty aldehydes (VLCF aldehydes), which are then converted to hydrocarbons in an irreversible reaction via Cyp4g1, a P450 oxidative decarbonylase. [18, 40] Importantly, adult oenocytes do not differentiate until late in pupal development and so they cannot be the site of synthesis for larval HL hydrocarbons. However, in *D. melanogaster* and other dipterans it is known that there are two separate generations of oenocytes, larval and adult, which differ from one another with respect to cell size, cell number and developmental origin (**Figure 2D**). [37, 60-65] Larval oenocytes form in the embryo and are known to play developmental roles in lipid metabolism, molting and tracheal waterproofing. [41, 66] In contrast to their adult counterparts, larval oenocytes have not been implicated in the synthesis of cuticular hydrocarbons. Intriguingly, larval oenocytes are reported to express Cyp4g1 from embryonic stages onwards [41]. We therefore used an oenocyte-specific driver (*PromE-GAL4*, Ref [34]) to express RNAi for *Cyp4g1* specifically in oenocytes throughout development. This oenocyte *Cyp4g1* knock down resulted in larvae with a dramatic decrease in fat body storage of HL hydrocarbons of the methyl-branched, monoene and diene classes (each hydrocarbon species is represented by meC28:0, C31:1 and C33:2 respectively, **Figure 2E**). To achieve spatiotemporal control, we then combined *PromE-GAL4* with the temperature-sensitive Gal80 system [67], selectively knocking *Cyp4g1* down in oenocytes during the pupal period, either with or without additional knockdown during larval development. This revealed an oenocyte *Cyp4g1* requirement for adult eclosion and viability that maps specifically to the larval phase of development (**Figure 2F**). Together, the *Cyp4g1* and HT-GC-MS analyses indicate that larval not adult oenocytes synthesize HL hydrocarbons, which are then stored in the larval fat body.

### EloHL is required in oenocytes for the biosynthesis of hyper-long hydrocarbons

The carbon-chain length of insect cuticular hydrocarbons is thought to be determined by the length of their VLCF-CoA precursors, which in turn depends upon fatty acid elongases (**Figure 2C**). To identify the elongase(s) responsible for synthesizing HL rather than L hydrocarbons, we purified actively translated mRNAs from larval versus adult oenocytes using a RiboTag method. [68] qRT-PCR assays of RiboTag purified mRNAs detected the expression of 15 of the 20 predicted *D. melanogaster* elongases in the larval and/or adult oenocytes of males and/or females. For these 15 elongases, PCA revealed that male and female elongase expression profiles cluster together in larval oenocytes (at the wandering L3 stage) but markedly separate in adult oenocytes (at 7 days after eclosion) (**Figure 3A**). This correlates with the unisex versus sex specific profiles of larval HL and adult L hydrocarbons respectively. The principal component coefficients (loadings) of individual elongases show that two elongases, *CG6660* and *CG8534,* contribute the most to the PCA separation between larval and adult oenocytes (**Figure 3B**). These two elongases are highly expressed in the larval oenocytes of both sexes but *CG8534* is also highly expressed in the adult oenocytes of females but not males (**Figure 3C**).

**Figure 3.**
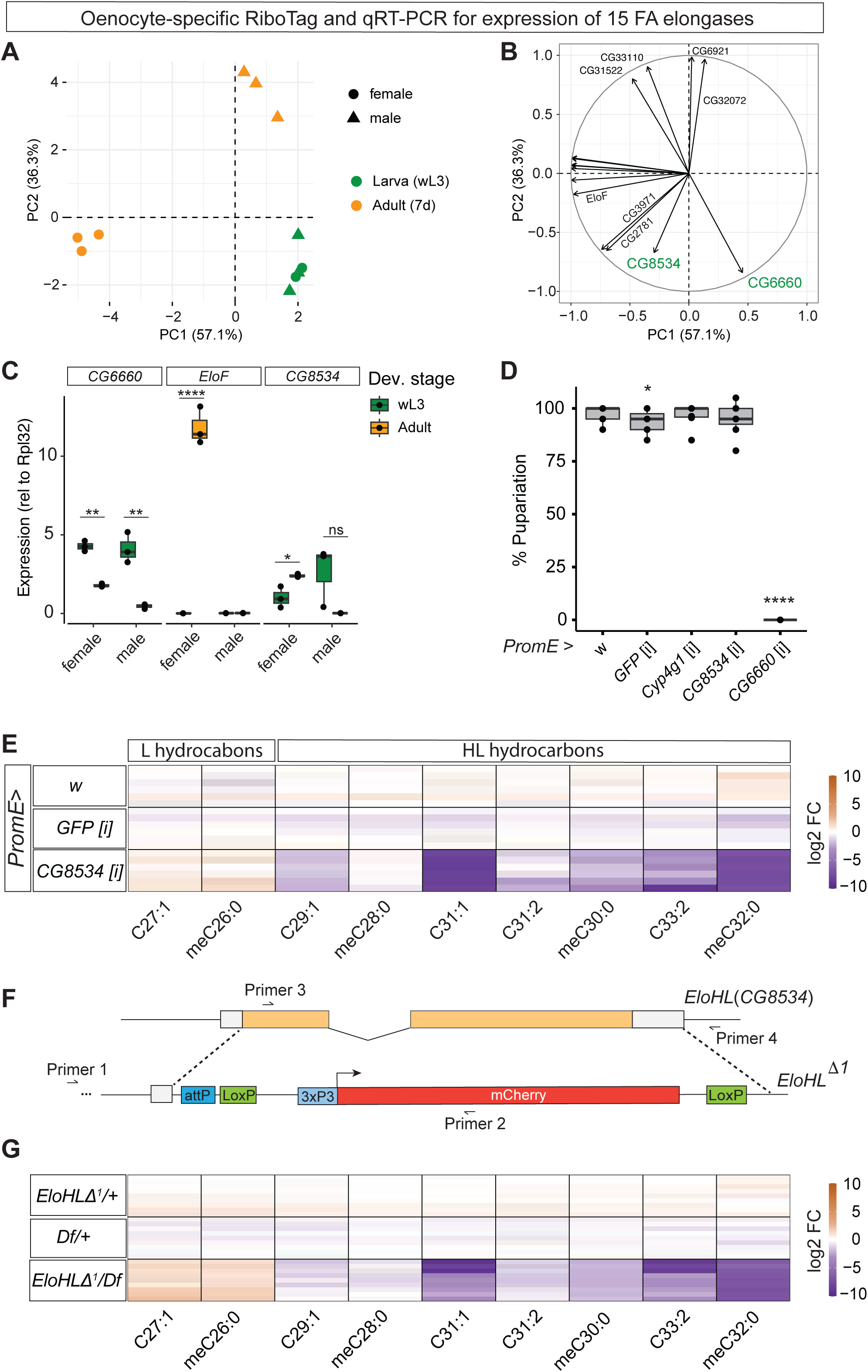
Larval oenocytes require EloHL for hyper-long hydrocarbon synthesis. (A-B) Oenocyte elongase expression transitions from unisex larval to sex-specific profiles. Principal component analysis (A) and loadings plot (B) of the expression levels of 15 elongases detected in oenocyte RNA purified by the RiboTag method from male or female wandering larvae (wL3) or 7-day old adult flies (adult 7d). Elongases and primers are listed in **Table S6**. N=3 independent experiments. (C) Relative expression of *EloF*, *CG6660* and *CG8534* (*EloHL*) in male and female oenocytes of larvae or 7-day adults (N=3 independent experiments). Statistical significance is indicated by *(p < 0.05), **(p< 0.05), ****(p<0.00005) and ns (p=0.08) using the tests listed in **Table S4.** (D) Larval viability of larva with oenocyte-specific knockdown of *Cyp4g1*, *CG8534* (*EloHL*) or *CG6660* versus controls. Larval viability was measured by counting the number of pupae arising from a defined number of L1 larvae. The graph depicts two independent replicates with at least three repeats each. Statistical significance is indicated by *(p < 0.05), ****(p<0.00005) using the tests listed in **Table S4.** (E) CG8534 (*EloHL)* is required for HL-HCs. Heatmap shows internal hydrocarbons of wandering L3 larvae with oenocyte-specific knockdown of CG8534. The fold changes of the indicated hydrocarbons are indicated relative to the mean of both controls (*PromE>GFP [i]* and *PromE>w*). Note that HL-HCs are decreased but L-HCs (C27:1 and me-C26:0) are increased in *PromE>CG8534 [i]*. (F) Generation of a mutant allele of *EloHL (CG8534)* using CRISPR-Cas9 mediated homologous recombination. In the *EloHL^Δ1^* allele, the *CG8534* open-reading frame (orange) and 3’ untranslated region (grey) are replaced with a LoxP flanked 3xP3-mCherry selection cassette and an attP site. Primers 1 to 4 were used to verify the structure of the resultant allele (**Figure S2**). (G) *EloHL* (*CG8534*) mutants recapitulate oenocyte-specific knockdown of *EloHL*. Heatmap shows internal HCs of wandering L3 larvae that are mutant transheterozygotes (*EloHL*^Δ1^/Df(3R)Excel6155) or control heterozygotes (*EloHL*^Δ1^/*+* or Df(3R) Excel6155/*+*). The fold changes of the indicated hydrocarbons are indicated relative to the mean of both heterozygous controls.

As a first step towards identifying whether *CG6660* or *CG8534* is required for the larval oenocyte synthesis of HL hydrocarbons, their developmental phenotypes were characterized. We reasoned that the RNAi knockdown phenotype of an oenocyte elongase dedicated to HL hydrocarbon biosynthesis would recapitulate, or be less severe than, that of *Cyp4g1*. Oenocyte-specific knockdown of Cyp4g1 gives normal larval development and pupariation but defective adult eclosion (**Figures 2F** and **3D**). Consistent with a previous study [66], larval oenocyte-specific knockdown of *CG6660* blocked all pupariation, ruling it out as an elongase dedicated to HL hydrocarbon synthesis (**Figure 3D**). In contrast, larval oenocyte-specific knockdown of *CG8534* did not affect pupariation (**Figure 3D**). We therefore used HT-GC-MS to quantify the internal hydrocarbons of these knockdown animals at the larval stage, observing a dramatic decrease in HL hydrocarbons, compared to *GFP* knockdown and *GAL4* driver only controls (**Figure 3E**). This decrease is accompanied by a corresponding increase in L hydrocarbons such as C27:1 and meC26:0 (**Figure 3E**). This suggests *CG8534* is specific for HL hydrocarbon biosynthesis but to rule out that the observed chain-length selectivity is due to partial knockdown, or even to RNAi off-target effects, we generated a precise null allele using CRISPR-Cas9 induced homologous recombination [69]. Both exons and the intron of *CG8534* were replaced with a 3xP3-mCherry cassette, thus removing the entire coding sequence (**Figure 3F**). The knockout mutant allele (*CG8534*^Δ1^) was verified by PCR, using primer pairs that detected the presence of 3xP3-mCherry and the absence of *CG8534* coding sequences in *CG8534*^Δ1^*/Df(3R)Excel6155* transheterozygotes (**Figure 3F and Figure S2**). Importantly, HT-GC-MS analysis of these *CG8534*^Δ1^ transheterozygotes, compared to control genotypes, revealed a strong decrease in HL hydrocarbons and a concomitant increase in L hydrocarbons, similar to the RNAi knockdown (**Figure 3G**). Together, the RNAi knockdown and null mutant analyses demonstrate that *CG8534* is required in larval oenocytes for HL but not L hydrocarbon biosynthesis. Furthermore, they suggest that, in the absence of *EloHL*, LCFA precursors are diverted towards other elongases that tend to produce shorter VLCF-CoA precursors for L hydrocarbons. Based on the selectivity of *CG8534* for HL hydrocarbons, we hereafter name it as *EloHL*.

### The switch from HL to L hydrocarbons is conserved across *Drosophila* species

To determine the extent to which HL hydrocarbons are conserved between *Drosophila* species, we first examined a chromosomal region of microsynteny that in *D. melanogaster* contains *EloHL* and four other predicted fatty acid elongases (*EloF*, *CG16904*, *CG9459* and *CG9458*). We identified the elongases in this microsyntenic region from 30 species across the *Drosophila* and *Sophophora* subgenera. This analysis suggests that the elongase microsynteny is restricted to the *Sophophora* subgenus (**Figure 4A**). Within the microsyntenic region of the 23 *Sophophora* species analysed, there are three examples of elongase duplications, but many species lack a recognisable ortholog of *EloHL* and/or the other elongases, and three species contain only *CG16904* (**Figure 4A**). To determine whether the presence of an *EloHL* gene correlates with HL hydrocarbons, six *Drosophila* species were selected for further comparative functional analysis. Two species have an *EloHL* gene (*D. melanogaster* and *D. biarmipes*) and four lack a clear microsyntenic ortholog of this elongase (*D. montana*, *D. ananassae*, *D. virilis* and *D. mojavensis*). HT-GC-MS analysis of cuticular hydrocarbons showed that there are clear unisex trends for shortening of the hydrocarbon chain length during the first week after adult eclosion in five of the six selected species. This is, however, the case not only in *D. melanogaster* and *D. biarmipes* but also in *D. ananassae, D. montana* and *D. virilis,* three species lacking microsyntenic *EloHL* (**Figures 4B**, **S3A** and **S3B**). Hydrocarbon shortening trends span *Drosophila* and *Sophophora,* but they do not correspond to a strict switch between HL to L molecules as we defined them in *D. melanogaster*. For example, in all species except *D. biarmipes,* one week old adults retain some C29 and other molecules of the HL hydrocarbon class. An extreme example of this is the desertic species *D. mojavensis,* with both males and females retain the complete developmental profile of HL hydrocarbons from eclosion to one week of age (**Figures 4B**, **S3A** and **S**). Together, the species comparisons show that HL hydrocarbons in newly eclosed flies are much more widespread across the *Drosophila* genus than the presence of a recognisable microsyntenic ortholog of *EloHL*.

**Figure 4:**
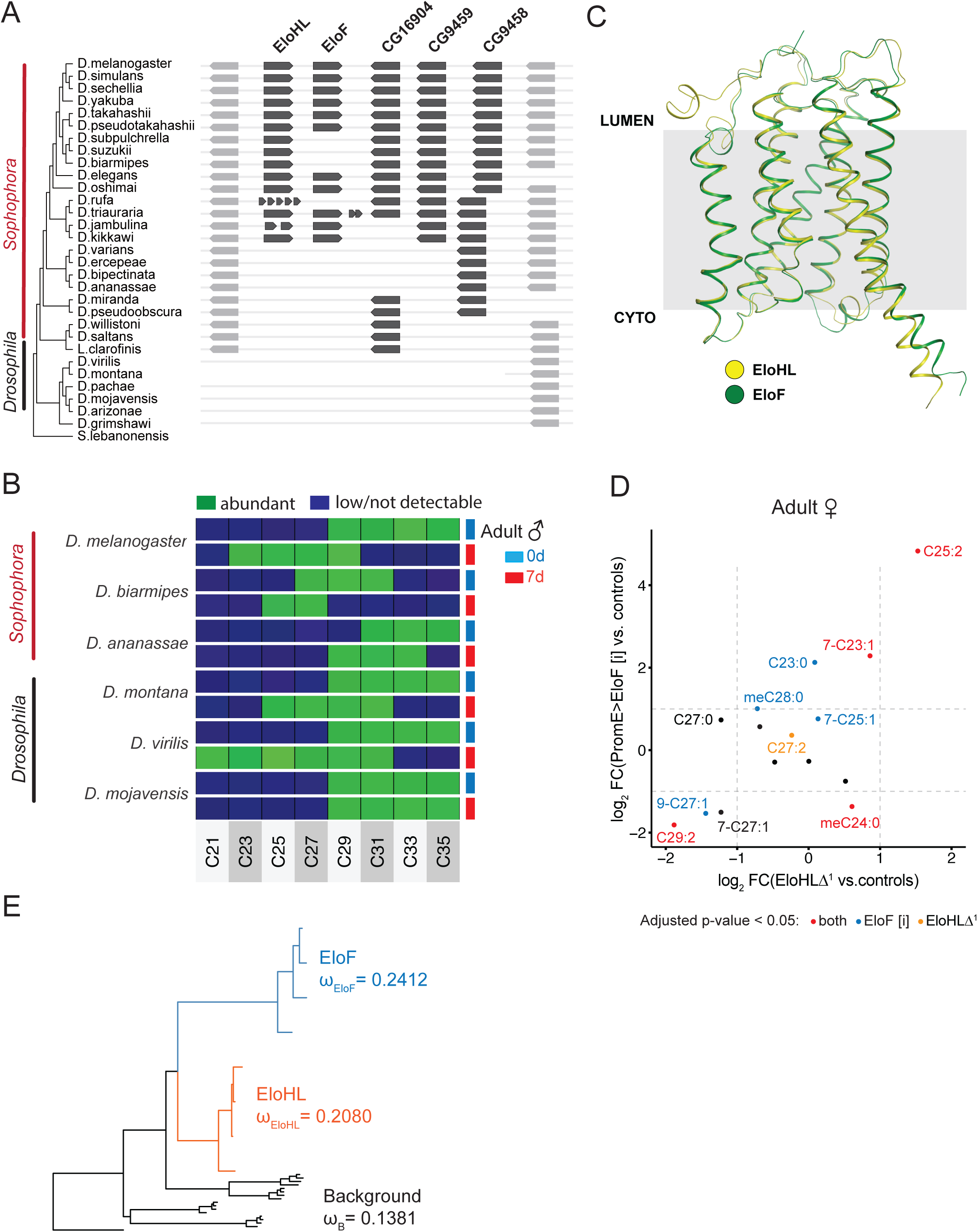
Hydrocarbon shortening in young adult male flies is widespread across *Drosophila* species. (A) Microsynteny of fatty acid elongases in the *EloHL* region is present in *Sophophora* (including *D. melanogaster*) but not in the *Drosophila* subgenus, another member of the *Drosophilidae* family. The *EloHL* loci were identified based on 3 flanking genes on each side in the *D. melanogaster EloHL* locus that are used as anchor genes in our analysis. Microsyntenic region in *S. lebanonensis* is not and in *D. montana* only partially identifiable. (B) Abundant cuticular hydrocarbon species of young (d0, light blue) and older (d7, red) male flies from species of the subgenera *Sophophora* (*D. melanogaster, D. biarmipes, D. ananassae*) and *Drosophila* (*D. montana, D. virillis, D. mojavensis*) indicated by a red and black bar respectively. Abundant (green) or low/not detected (dark blue) hydrocarbons measured by HT-GC-MS are indicated according to the number of carbon atoms (C23 to C37) and pooling alkanes, monoenes, dienes and methyl-branched alkanes. The adult transition from hyper-long to standard length cuticular hydrocarbons is observed across both *Sophophora* and *Drosophila* males except in the desertic species *D. mojavensis*. (C) Comparison of AlphaFold structures of *EloHL* and *EloF*. Predicted EloHL and EloF structures are very similar and predicted to span the ER membrane from the cytoplasm (cyto) to the lumen. (D) EloHL and EloF synthesize similar CHCs in adult females. Surface HCs from 7-day old females with adult oenocyte-specific knockdown of EloF using the PromE^TS^ driver (*PromE-GAL4, UAS-GAL80^TS^, UAS-CD8::GFP)* were compared to PromE^TS^>GFP [i] controls. Similar, surface HCs from 7-day old females EloHL mutant transheterozygotes (*EloHL*^Δ1^/Df(3R) Excel6155) were compared to both control heterozygotes (*EloHL*^Δ1^/*+* or Df(3R)Excel6155/*+*). Significance for both EloF knockdown and EloHL mutants was tested using Anova with Bonferroni correction (**Table S4**). Dots are coloured according to significance in either both EloF knockdown and EloHL mutants (red), only EloF knockdown (blue) or only EloHL mutants (yellow). EloF log2 fold changes were plotted on the y-axis and EloHL log2 fold changes on the x-axis. (E) dN/dS (ω) ratios of fatty acid elongases in the *EloHL* microsynteny. ω for *EloF* but not EloHL is significantly higher (p=0.005) ω than for the background elongases (*CG16904, CG9458, CG9459*). Calculations are based on the elongase sequences from *D. melanogaster, D.simulans, D. sechellia*, and *D.yakuba*, each of which have five fatty acid elongases in the microsynteny. Results of the different models tested are shown in **Table S5**.

Within the *EloHL* microsynteny region, one other closely related elongase (*EloF*) has been shown to synthesize cuticular hydrocarbons. *EloF* is required for adult oenocyte synthesis of pheromones in females but not males. [45, 47] Consistent with this, there is high expression of *EloF* in the adult oenocytes of females but not males, but we also find that there is no detectable *EloF* expression in the larval oenocytes of either sex (**Figure 3C**). Protein comparisons between EloF and EloHL in *D. melanogaster* show high sequence conservation (59% identity) and close superposition of their 3D structures predicted by AlphaFold (**Figure 4C**). Moreover, a comparison of sexually mature adult females one week after eclosion showed a strong correlation between the cuticular hydrocarbon molecules up or down-regulated in *EloHL^Δ1^* mutants and in *EloF* knockdowns (**Figure 4D**). Strikingly, loss-of-function of either of these two elongases leads to a strong downregulation of the female-specific pheromone C29:2 (nonacosadiene), whereas its shorter pheromonal counterpart, C27:2 (heptacosadiene), barely changes and C25:2 (pentacosadiene) is markedly upregulated (**Figure 4D**). Some monoenes and mb-alkanes are, however, differentially affected by the loss-of-function of each elongase. These findings strongly suggest that *EloHL* and *EloF* are both required in sexually mature adult females for the biosynthesis of overlapping sets of HL hydrocarbons.

To gain insight into the potential functional divergence of *EloHL* and *EloF* during evolution, we estimated selection pressures using phylogenetic analysis by maximum likelihood (PAML). [70] This method was used to calculate nonsynonymous/synonymous rate ratios (*d*_N_/*d*_S_ = *ω*) for gene sequences from *D. melanogaster, D. simulans, D. sechellia, and D. yakuba*, four species that have one copy of each of the five microsyntenic elongases. One gene, *CG31141*, outside the microsynteny, was also included due to its high sequence similarity to the five elongases. We tested the hypothesis that EloF evolves at a different rate than other elongases (model B in Table S5) compared to the hypothesis that all elongases evolve at a similar rate (Model A in Table S5), and found that EloF had a significantly higher rate of evolution than other genes (P = 0.005). We also tested whether EloHL evolves differently from other elongases (Model C in Table S5) but did not find a significant difference (P = 0.12). When comparing EloHL, EloF, and four other elongases, we observed that EloF had the highest evolutionary rate (ω = 0.24), compared to EloHL (ω = 0.20) and the other elongases (ω = 0.14) (**Figure 4E and model D in Table S5**). These results suggest that *EloHL* has experienced similar selective pressures as other genes, while *EloF* has evolved more rapidly.

### Larval synthesis of hyper-long hydrocarbons protects the pupa from desiccation

We next investigated whether an evolutionarily constrained function of *EloHL,* not shared with *EloF*, might involve its unisex larval role in HL hydrocarbon synthesis. Although *EloHL* is required for larval HL hydrocarbons, it is not required for the completion of larval development itself (**Figures 3D-3G**). Nevertheless, it remains possible that larval HL hydrocarbons synthesized during the larval stage are stored in the fat body, in order to function later on in development - during the pupal period. To address this possibility, we asked whether *EloHL*-dependent HL hydrocarbons are required for the pupal barrier to desiccation, which remains mysterious both in terms of its chemical composition and anatomical location. *EloHL^Δ1^*/*Df(3R)Excel6155* transheterozygotes, deficient in HL hydrocarbons, were subjected from pupariation onwards to varying degrees of desiccation stress induced by low relative humidity (RH) and/or high temperature. First, we established that the pupal dry weights of *EloHL^Δ1^*/*Df(3R)Excel6155* transheterozygotes, as well as oenocyte-specific *Cyp4g1* knockdowns, were not significantly different from those of genetic controls (**Figure S4A and S4B**). At 30% RH and 25°C, the pupal water content of genetic controls declines rapidly by 24 hr after puparium formation (APF) and subsequently slows, such that it is roughly halved by 72 hr APF (**Figure S4C**). Under these environmental conditions, the pupal water loss of *EloHL^Δ1^*/*Df(3R)Excel6155* transheterozygotes showed a very similar temporal pattern and magnitude to that of genetic controls (**Figure S4C**). In contrast, with drier conditions of 15-20% RH, *EloHL^Δ1^* transheterozygotes lost more water by 48 hr APF than controls, and this was observed across temperatures from 25 to 28°C (**Figure 5A**). At 29°C under these drier conditions, there was a catastrophic loss of ∼75% of pupal water by 48 hr APF, both in the transheterozygotes and the genetic controls (**Figure 5A**). At 15-20% RH and 26°C, we observed that the excessive rate of dehydration of *EloHL^Δ1^* transheterozygotes is apparent by 24 hr and continues until 48 hr APF (**Figure 5B**). The dehydration assays demonstrate that *EloHL* is required for an efficient pupal desiccation barrier that likely involves HL hydrocarbons.

**Figure 5:**
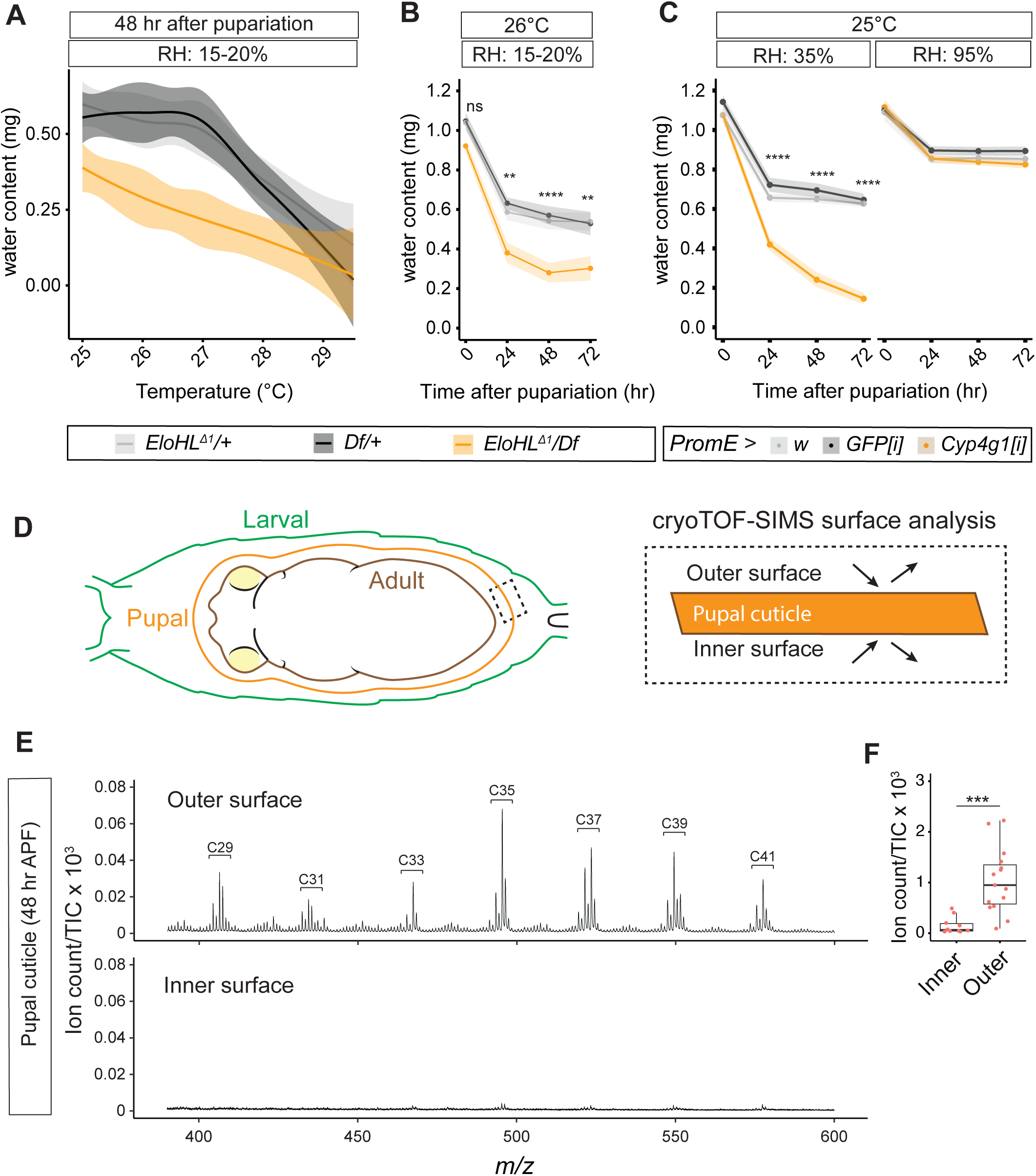
EloHL-dependent hyper-long hydrocarbons coat the external surface of the pupal cuticle and protect from desiccation. (A) Pupal water content (mg) versus temperature (°C) of *EloHL*^Δ1^/*Df(3R)Excel6155* mutants and heterozygous controls (*EloHL*^Δ1^*/+* or *Df(3R)Excel6155/+)* at 48 hr after pupariation with 15-20% relative humidity (RH). A loess curve was fitted to the data by genotype, showing that *EloHL*Δ^1^/Df has increased sensitivity to water loss at lower temperatures. (N=6, n≥5) (B) Pupal water content (mg) of *EloHL*^Δ1^/*Df(3R)Excel6155* mutants and heterozygous controls (*EloHL*^Δ1^/+; Df/+) with 15-20% relative humidity at 26°C. Statistical significance is indicated by **(p < 0.005), ****(p<0.00005) using the tests listed in **Table S4.** (N=4, n≥5) (C) Pupal water content (mg) of oenocyte-specific *Cyp4g1* knockdown (*PromE>Cyp4g1[i])* and control (*PromE/+* or *PromE>GFP [i])* animals at 25°C with either 35% or 95% relative humidity (RH). Oenocyte-specific knockdown of *Cyp4g1* leads to early water loss within 24 hr of pupariation at 35% RH but this can be prevented with 95% RH. Statistical significance is indicated by ****(p<0.00005) using the tests listed in **Table S4.** (N=2, n = 10) (D) Diagram showing the three cuticles of the late pupa. From external to internal, there is the larval cuticle, the pupal cuticle and the adult cuticle. **For more details see** Figure 6. Hydrocarbons on the outer versus the inner surfaces of the pupal cuticle can be measured using cryogenic time-of-flight secondary ion mass spectrometry (cryoTOF-SIMS). (E) Representative cryoTOF-SIMS spectra from the inner and outer surfaces of the pupal cuticle at 48 hr after puparium formation (APF). Hydrocarbons from C29 to C41 are indicated. The Y axis indicates ion count relative to the total ion count (TIC). (F) Quantification of total hydrocarbon signals on inner and outer surfaces of the pupal cuticle. Hydrocarbon signals from C29-37, normalized to the total ion count (TIC), are pooled. (N=2, n≥5)

*EloHL* mutants and larval oenocyte-specific *EloHL* knockdown both retain L hydrocarbons, such as C26 and C27 molecules, during development. This contrasts with the almost complete lack of all hydrocarbon synthesis in *Cyp4g1* knockdowns (**Figure 2E**). Using a mild desiccation stress (30% RH and 25°C) that does not elevate dehydration in *EloHL* mutants, water loss in larval oenocyte-specific *Cyp4g1* knockdown pupae is nevertheless very pronounced, almost double that of genetic controls (**Figure 5C**). Importantly, this excessive water loss is already present at 24 hr APF, three days before adult eclosion and long before adult oenocytes differentiate and synthesize hydrocarbons (**Figure 5C**). Remarkably, the excessive water loss in *Cyp4g1* knockdown pupae at 25°C could be fully rescued by raising the humidity from 30% to 95% RH (**Figure 5C**). The stronger pupal dehydration phenotype of *Cyp4g1* compared to *EloHL,* together with our biochemical analyses, strongly suggests that L and HL hydrocarbons both contribute to the pupal desiccation barrier. Nevertheless, loss of the HL class alone in *EloHL* mutants is enough to compromise barrier efficiency in high temperature and/or arid environments.

The results thus far raise important questions about the anatomical location of the pupal hydrocarbon barrier to desiccation. Multiple cuticles with barrier functions are synthesized during insect development and, in holometabolous insects like *Drosophila,* there are three cuticles present during metamorphosis i.e from pupariation until adult eclosion. From external to internal, these are the third-instar (L3) larval cuticle, which hardens to form the puparium (external pupal case), the much thinner and more flexible pupal cuticle that envelopes the forming adult, and the adult cuticle itself (**Figure 5D**). The puparium persists throughout metamorphosis but we were unable to detect significant hydrocarbons on its external surface (**Figure 1A**). Moreover, although there are HL hydrocarbons on the surface of the adult cuticle in newly eclosed flies, this structure does not form until ∼36 hr APF [71], long after the onset of excessive pupal dehydration in *EloHL* mutants and *Cyp4g1* knockdowns. However, the pupal cuticle forms at ∼12 hr APF [72], early enough to participate in the pupal desiccation barrier. Dissected pupal cuticles were therefore analysed at 48 hr APF for the presence of hydrocarbons using cryogenic time-of-flight secondary ion mass spectrometry imaging (cryoTOF-SIMS), a surface-specific analytical technique that we previously optimised for adult cuticular hydrocarbons. [73] Hydrocarbons on the pupal cuticle were largely absent from the internal surface but were detected in abundance on the external surface, ranging from C29 to C41 HL hydrocarbons (**Figure 5D-5F**). In summary, we have identified a unisex hydrocarbon barrier to desiccation during *Drosophila* metamorphosis and shown that it is synthesised by *EloHL* in larval oenocytes, stored in the larval fat body, and then mobilized to the external surface of the pupal cuticle.

## DISCUSSION

This *Drosophila* study fills a gap in our understanding of how holometabolous insects cope with the threat of dehydration during metamorphosis. It identifies the cell source, chemical basis, mechanism of synthesis, and anatomical location of a lipid barrier specialized for a developmental stage particularly vulnerable to water loss. The pupal lipid barrier we identify differs from its adult counterpart in four important ways. First, it is synthesized by larval oenocytes, a separate cell population from the adult oenocytes that produce the lipid barrier of the imago. Second, it is stored in the larval fat body until required at the pupal stage. Third, it is comprised of unisex hydrocarbons with longer average carbon chain lengths than the sex-specific hydrocarbons of the adult. And fourth, its anatomical location is on the exterior surface of the pupal cuticle, a separate structure from the adult cuticle. We now discuss how separate hydrocarbon barriers are synthesized during development and adulthood, and how their evolution may be shaped by distinct selective pressures on desiccation resistance and pheromonal signalling.

### Larval and adult oenocytes synthesize different hydrocarbon barriers

Despite their shared name, larval and adult oenocytes are different cells with distinct developmental origins and functions. [35, 36, 41, 61, 66, 74] Our study reveals a hitherto unknown role of larval oenocytes in desiccation resistance during metamorphosis. It also provides the first unifying evidence that both types of oenocytes make hydrocarbon lipid barriers, albeit chemically distinct ones. Larval oenocytes synthesize the hyper-long hydrocarbons of the pupal and immature adult cuticle, whereas adult oenocytes make the sex-specific shorter hydrocarbons of the mature adult cuticle. Both cell types utilize Cyp4g1 for hydrocarbon biosynthesis but, in contrast to previous interpretation [40], we find that it is the larval not the adult oenocytes that require *Cyp4g1* to make the hydrocarbon barrier relevant for survival at adult eclosion.

Both larval and adult oenocytes express the same terminal biosynthetic enzyme for hydrocarbons, Cyp4g1 (Ref [40]), but the different chain length ranges they produce appear to be specified by distinct fatty acid elongases. Hence, unisex *EloHL* expression in larval oenocytes enables them to produce hydrocarbons longer than C27, which are critical for an optimum pupal desiccation barrier. Larval oenocytes also express another gene (*CG6660*) encoding an elongase [66] but it is not yet clear whether or not this also contributes to the synthesis of hyper-long hydrocarbons. Adult oenocytes on the other hand express a range of sex-biased elongases, including female-specific *EloF*, which produces shorter (up to C29) blends of hydrocarbons with dual barrier and pheromonal functions. Interestingly, we found that *EloHL* is not only expressed in a unisex pattern in larval oenocytes but also redeployed in a sex-specific manner in adult oenocytes to synthesis long female pheromones such as 7,11-nonacosadiene (C29:2). [75, 76]

A key finding of this study is that larvae synthesize hyper-long hydrocarbons but do not transport them to the cuticular surface. Instead, hyper-long hydrocarbons are stored internally, within the larval fat body. This is one of very few larval tissues that does not undergo histolysis during metamorphosis, persisting through to the sexually immature adult. [77, 78] Larval oenocytes, in contrast, histolyse at the midpupal stage [41] and so cannot supply hydrocarbons to the pupal or early adult cuticles during the remainder of metamorphosis. The persistence of the larval fat body is therefore able to ensure an internal store of barrier hydrocarbons throughout metamorphosis, bridging the developmental gap between the larval and adult oenocyte sources (**Figure 6**). Our study shows that larval-oenocyte derived hyper-long hydrocarbons are required for the desiccation barrier of the pupal cuticle. Nevertheless, other cuticular components may also contribute and it is very likely that non-cuticular mechanisms of water homeostasis also play significant roles during metamorphosis.

**Figure 6:**
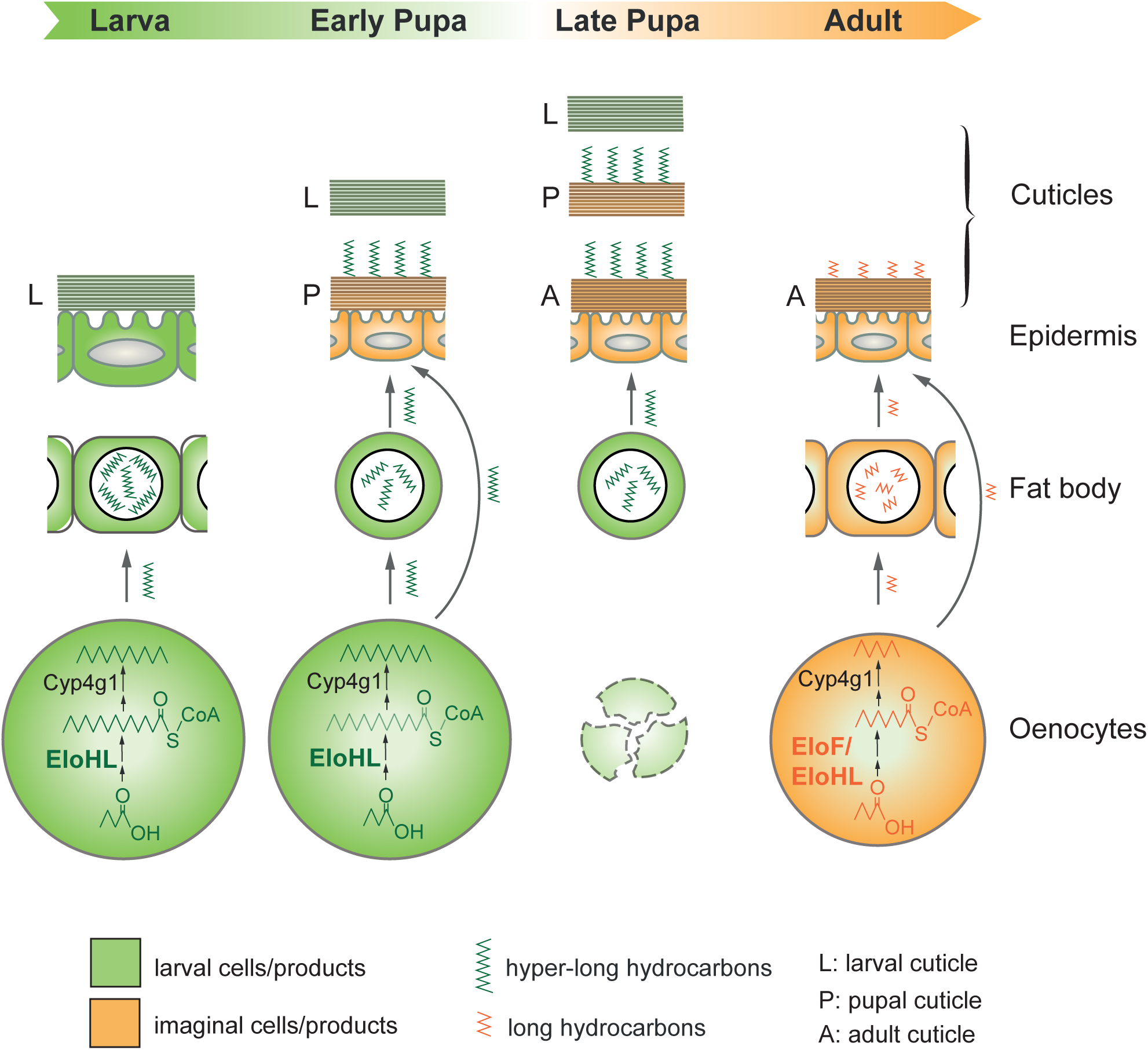
The formation of the desiccation barrier for *Drosophila* metamorphosis. *Drosophila melanogaster* undergoes three larval stages, one pupal stage, and one adult stage, with each stage characterized by a distinct cuticle that is either morphologically or chemically unique. Cuticular hydrocarbons (HCs) are absent from the surface of the larval cuticle. However, unisex hyper-long HCs (HL-HCs) are synthesized in larval oenocytes through the action of the fatty acid elongase EloHL and the cytochrome P450 enzyme Cyp4g1. These HL-HCs are then stored in the larval fat body, a tissue that does not undergo histolysis during metamorphosis. During the early pupal stage, the former larval cuticle hardens to form a rigid pupal case, while a softer pupal cuticle begins to form underneath it approximately 10 hours after puparium formation. At this point, HL-HCs are transported from the fat body - and possibly from oenocytes - to the surface of the pupal cuticle. Notably, larval oenocytes are among the last tissues to undergo histolysis during the late pupal stage, coinciding with the synthesis of the adult cuticle. The HL-HCs, which likely originate from the fat body store, are then transferred to the surface of the adult cuticle as oenocytes have already histolyzed by this stage. In mature adult stages, adult oenocytes synthesize sex-specific cuticular hydrocarbons by expressing sex-specific elongases, such as the female-specific EloF and EloHL, along with the Cyp4g1 enzyme. The transport of cuticular hydrocarbons to the surface of the adult cuticle may occur either directly from oenocytes or via the fat body, although this mechanism has not yet been fully investigated.

### Specialized lipid barriers for distinct life stages and ecological niches

*Drosophila* and other holometabolous insects undergo dramatic morphological and physiological changes during their life course, often in correlation with changes in their ecological niche. We found that the larva, which lives in a semi-aquatic environment, lacks detectable hydrocarbons on the surface of its cuticle. In contrast, the pupa occupies a drier terrestrial niche and it is coated with unisex hydrocarbons that are hyper-long. Physicochemical and biological studies of insect hydrocarbons indicate that longer chain lengths and higher proportions of methyl-branched and linear alkanes promote lower volatility and higher desiccation resistance. [5, 28, 51] Hyper-long hydrocarbons of the pupal lipid barrier are therefore likely to have evolved to minimize water loss but presumably they are not volatile enough to function well as olfactory pheromones at temperate climate. The adult flies of *Drosophila* species that communicate using sex-specific olfactory pheromones therefore need to transition to shorter more volatile hydrocarbons. These shorter hydrocarbons appear to occupy a physicochemical sweet spot, long enough to support barrier function yet short enough for some of the alkenes to work as volatile olfactory pheromones.

Our survey of six *Drosophila* species, together with previous studies [48, 79, 80] adds to the evidence that the transition from hyper-long to shorter hydrocarbons during the first day after adult eclosion is widespread across *Drosophila* species. Interestingly, however, we found that hyper-long hydrocarbons persist in one day adults of a desert-dwelling species *D. mojavensis*. We speculate that this reflects the ecological pressure for high desiccation resistance in an arid environment. *D. mojavensis*, like some other *Drosophila* species expressing hyper-long hydrocarbons does not have a recognizable microsyntenic orthologue of EloHL. It is therefore likely that other elongases fulfil a related role to EloHL in these species. Consistent with this, a recent study identified a novel elongase (mElo) in *D. mojavensis* that elongates methyl-branched hydrocarbons with up to 33 carbons. [43] *EloHL* contributes to both desiccation resistance and female pheromone biosynthesis, overlapping in the latter function with *EloF*. Sequence analysis indicates that *EloHL* has likely evolved with greater negative purifying selection than *EloF*. This suggests that *EloHL* could have important constrained function(s) that are separate from the rapidly evolving function it shares with *EloF* in female sex pheromones, which is thought to contribute to interspecies mating barriers. [16, 24, 34, 45, 53, 81] We speculate that this constrained function of EloHL could correspond to synthesis of the pupal hydrocarbon barrier.

Together, the findings in this study identify the mechanisms accounting for how hydrocarbon barrier adjust across an insect life course to suit different ecological niches. However, they also highlight that there are environmental limits beyond which hydrocarbon barriers fail. Hence, we found that the pupal hydrocarbon barrier is not able to protect metamorphosis from lethal desiccation at low humidities and high temperatures found in many areas of our planet. Given the pace of climate change, this may constrain the ability of some *Drosophila* species to adapt rapidly enough to survive in their current geographical ranges.

## METHODS

### Fly Husbandry

All flies were maintained at 25°C, 60% humidity, and a 12-hour light/dark cycle unless otherwise specified. The *Drosophila melanogaster* UAS-RNAi lines used in this study were obtained from the Vienna Drosophila RNAi Stock Centre (VDRC) or the Bloomington Drosophila Stock Centre (BDSC) (*CG8534*: 49892 (VDRC); *CG6660*: 62422 (Bloomington); *EloF*: 44660 (Bloomington); *Cyp4g1*: in-house). Note that the Bloomington RNAi line (ID: 53299) for *CG8534* has an off-target causing an eclosion defect and was therefore excluded from the manuscript. These lines were crossed with the well-described GAL4 driver lines, either *PromE* (*w*; *PromE-GAL4*, *UAS-CD8::GFP*/*CyO*) or *PromE^TS^* (*w*; *PromE-GAL4*, *UAS-CD8::GFP*, *UAS-tub-GAL80^TS^*/*CyO*). [34] For each genetic cross, larvae or eggs were collected from grape juice agar plates (25% v/v grape juice, 1.25% w/v sucrose, and 2.5% w/v agar) and transferred to vials containing standard yeast-cornmeal-agar medium (w/v: 6.63% cornmeal, 5.85% glucose, 2.34% autolysed yeast, 0.195% Nipagin, 0.00078% Bavistin, 0.7% agar, 1.95% ethanol (v/v)).

For stage-specific knockdown of *Cyp4g1* in oenocytes, the temperature-sensitive *PromE^TS^* driver was used. Larvae were maintained at either 18°C or 29°C during development, and larvae raised at 18°C were transferred to 29°C when pupariation occurred. Pupariation was assessed by transferring 20 L1 larvae of the appropriate genotype to a vial and counting the number of pupae that formed. Adult eclosion was measured by counting the number of adults that successfully emerged from the pupal case.

### Sample Preparation for GC-MS

For surface HC samples, five wandering L3 larvae or pupae were collected from the side of the vial, washed with PBS, and weighed on a microscale after drying on tissue paper. Adult flies were anesthetized with CO_2_ and weighed on a microscale. Pooled individuals were washed in 50 µl hexane with 0.1 mM octadecane (internal standard) in a GC vial for 2 minutes by actively agitating the individuals in the solution. The hexane extract was then transferred to a new GC vial and capped.

For internal HC sample preparation, five wandering L3 larvae or pupae were collected, washed with PBS, dried on tissue paper, and weighed on a microscale. Adult flies were anesthetized with CO2 and weighed on a microscale. External hydrocarbons were removed before homogenisation using a hexane dip as described above. For larval tissues, ten wL3 larvae were dissected in PBS into fat body, inverted carcass (with all internal tissues and the head removed), and remaining internal organs (gut, Malpighian tubules, crop) referred to as gut. Pooled individuals or tissues were transferred to a metal PrecellysⓇ tube with 500 µl hexane with 0.01 mM octadecane (internal standard) and homogenized using the PrecellysⓇ Evolution tissue homogenizer. Samples were homogenized for four rounds (25 seconds at 6000 rpm), transferred to an Eppendorf tube, and centrifuged for ten minutes at 14,000 rpm. The supernatant was transferred to a GC vial, and hexane was allowed to evaporate overnight in a fume hood. Once the hexane had fully evaporated, the samples were reconstituted in 50 µl hexane and capped.

### Gas Chromatography-Mass Spectrometry (GC-MS) Lipidomics

GC-MS was performed using an Agilent 7890B-7000C GC–MS system in EI mode. Data acquisition and analysis were carried out using MassHunter (Version 10.0, Agilent Technologies, Inc.). Splitless injection of 1 µl (injection temperature 310°C) onto a 30 m + 10 m × 0.25 mm BPX5 column (SGE) was used, with helium as the carrier gas. The initial oven temperature was 100°C (1 min), followed by a temperature gradient to 350°C at 10°C/min with an optional hold time of 25 minutes for internal HC samples. Sample running order was randomized using the MassHunter sample sequence randomizer. Data analysis was performed using MassHunter Qualitative and Quantitative Analysis (version 10.0, Agilent Technologies). Peak areas were first normalized to an internal standard (octadecane) spiked into the sample at the time of metabolite extraction. Compounds were identified by referencing commercial standards when available, or by retention time and fragmentation pattern when standards were unavailable. After normalization to the internal standard, data were square root transformed for visualization.

### Ribotag-RNA Isolation and qPCR

The oenocyte-specific Ribotag protocol was adapted from Huang *et al*. [68]. PromE-Gal4 driver virgins were crossed to male UAS-RPL13a-Flag flies (Bloomington: 83684). Eggs were collected from grape juice agar plates and transferred to fly vials with standard food. Thirty male or female wandering L3 larvae or adult flies were snap-frozen in liquid nitrogen and stored at -80°C. On the day of Ribotag analysis, samples were thawed on ice and transferred to plastic PrecellysⓇ tubes filled with ice-cold lysis buffer (20 mM HEPES-KOH pH 7.4, 12 mM MgCl2, 100 mM KCl, 1% NP-40, 1 mM DTT, 100 µg/ml cycloheximide, 40 U/ml RNAsin, protease inhibitor). The samples were homogenized for two rounds (25 seconds at 6000 rpm) using the PrecellysⓇ Evolution tissue homogenizer. The lysate was transferred to a new tube and centrifuged for 10 minutes at 10,000 rpm at 4°C. Dynal Protein G Magnetic beads were washed three times with wash buffer (20 mM HEPES-KOH pH 7.4, 12 mM MgCl2, 100 mM KCl, 1% NP-40, 1 mM DTT, 100 µg/ml cycloheximide, 40 U/ml RNAsin), added to the sample lysate, and incubated with end-over-end rotation for 1 hour at 4°C to pre-clean the lysate. Another set of Dynal Protein G Magnetic beads was coupled to anti-FLAG antibody by incubating washed Protein G beads with anti-FLAG antibody (Sigma Aldrich; Catalog number: F1804) with end-over-end rotation for 1 hour at room temperature, followed by five washes with wash buffer. The supernatant of the pre-cleaned sample lysate was then transferred to the anti-FLAG antibody-coupled beads and incubated by end-over-end rotation for 4 hours at 4°C. After incubation, the beads were washed three times with high salt buffer (20 mM HEPES-KOH pH 7.4, 12 mM MgCl2, 350 mM KCl, 1% NP-40, 1 mM DTT, 100 µg/ml cycloheximide, 40 U/ml RNAsin), with the last wash rotating end-over-end for 20 minutes at 4°C, followed by two additional washes. After the final wash, 500 µl TriZol was added to the beads, the sample was vortexed, and incubated for 10 minutes on ice. Finally, the beads were collected using a magnetic rack, and the supernatant was transferred to a new tube. RNA isolation was performed using the Direct-zol RNA Microprep Kit (Zymo). RNA was reverse-transcribed to cDNA using the Superscript IV cDNA Kit, and elongase expression was measured using the Roche Lightcycler with the Lightcycler Master I Mastermix. Gene expression was normalized to the housekeeping gene RPL32.

### Generation of the EloHL^Δ1^ Mutant

The EloHL^Δ1^ mutant was generated following the strategy described. [82] The TVΔattp-Pax-mCherry and CFD4 vectors were used for the targeted deletion of *CG8534*. The TVΔattp-Pax-mCherry vector features an attP site and an excisable Pax-mCherry cassette flanked by LoxP sites, enabling visual tracking of the mutation. Genomic deletions were achieved by co-injecting the TVΔattp-Pax-mCherry vector with the CFD4 vector, which expresses two guide RNAs targeting the deletion breakpoints at the ends of the 3’UTR and 5’UTR of CG8534. The vectors were co-injected into the nos-Cas9-mSA fly line with an attP2 landing site, which carries Pax-GFP as a genetic marker. Integration of the TVΔattp-Pax-mCherry cassette was confirmed by PCR analysis using Primer 1 and Primer 2. Primer 1 is located outside the homology arm used to integrate the cassette, while Primer 2 is located within the mCherry sequence. Additionally, the absence of CG8534 was confirmed in transheterozygous mutants (EloHL^Δ1^/Df(3R)Excel6155) using PCR with Primer 3 and Primer 4, which amplify part of the CG8534 cDNA.

### Desiccation Assay

For pupa desiccation assays, pupae were collected shortly after pupariation and removed from the side of the vial with a wet paintbrush. Individual pupae were transferred to a pre-weighed 0.2 ml PCR tube without a lid, and the tube was weighed again with the pupa using a microscale. Pupal desiccation resistance was tested under two conditions: (1) Tubes with pupae were kept in a pipette tip box at the specified temperature and relative humidity after pupariation. Relative humidity was measured using an RH probe, and high RH levels (∼95%) were achieved by adding wet tissues to the bottom of the pipette tip boxes. The weight of the tube plus pupa was measured every 24 hours after pupariation. (2) Tubes with pupae were transferred to a PCR machine with different temperatures set as constant incubation (25-29°C). Relative humidity was not controlled but measured, varying between 15-20%. The weight of the tube plus pupa was measured every 24 hours after pupariation. After 72 hours APF or when the pupa turned dark, the tubes with pupae were dried at 60°C overnight, and the weight of the tube plus pupa was measured again. The pupal wet weight was calculated by subtracting the empty tube weight from the ‘tube + pupa’ weight. The pupal dry weight was calculated by subtracting the empty tube weight from the dried ‘tube + pupa’ weight, and the pupal water content was calculated by subtracting the pupal dry weight from wet weight.

### cryoTOF-SIMS surface analysis

Pupae were collected from the side of the vial at 48 hours post-pupariation. The pupae were stripped of their larval cuticles by attaching them to double-sided tape and removing the cuticles with forceps. The pupae were then transferred to PBS, and a small incision was made at the anterior end, the pupal cuticle was cut longitudinally, and internal tissues were removed. The pupal cuticles were then washed twice with PBS and twice with 150 mM ammonium formate. ITO-coated glass coverslips (SPI Supplies #6462-AB, 18 mm², resistivity 70-100 Ω) were cut to size with a diamond knife, and the pupal cuticles were mounted with either the internal or external surface facing upwards.

ToF-SIMS measurements were performed on a ToF-SIMS V instrument (IONTOF, GmbH) as previously described by Newell *et al*. [73] Samples on the ITO-coated glass slides were mounted on a cooling stage. The loadlock was flooded with nitrogen gas before cooling during pump-down. During the analysis, the sample was cooled with a copper cooling finger to approximately -110°C. Analyses were conducted using a 30 keV Bi3+ analysis beam with a spot size of approximately 1 μm and a current of 0.04 pA at a cycle time of 200 μs. Secondary ions were extracted with an extraction voltage of 1 kV, with delayed extraction mode activated. For measurements used to generate graphs, data was obtained from a field of view of 250 x 250 μm at a resolution of 128 x 128 pixels. The measurements shown are a sum of 20 scans. To compensate for sample charging, a flood gun was applied during analysis with an energy of 21 eV and a current of -10 μA. The total ion dose was approximately 8.18 x 10⁷ ions/cm². The mass resolution of the resulting spectra was approximately 5,000 at m/z 200. Spectral mass calibration was performed with reference to the known ions [C]+, [CH]+, [CH2]+, and [CH3]+. For measurements used to plot mass spectra, a total of 50 scans were acquired, with a total ion dose of approximately 2.05 x 10⁸ ions/cm². For the quantification shown in Figure 5C, the [M]^+.^ ion from hydrocarbons ranging in length from C29-C37 were summed. For these hydrocarbons, presence and positive identification has been confirmed in pupal samples with GC-MS.

### Homologous Gene Identification and Evolutionary Analysis

To identify all fatty acid elongases across species, we performed TBLASTN [83] to search against the reference genomes of each species (in Figure 4A) using the protein sequences of the fatty acid elongases gene group from *D. melanogaster* (FBgg0000965) as queries, with a cutoff e-value of 1×10^-5^. Redundant hits in the same genomic regions were filtered out, and the sequences of all hits were extracted with extensions at both the 5’ and 3’ ends. Coding sequences were determined using Exonerate. [84] To identify homologous regions and fatty acid elongases in the microsynteny, we conducted TBLASTN searches using the protein sequences of flanking genes (CG9467, CG8526, FBXO11 on one side and CG42857, CG34302, Teh1 on the other side) from *D. melanogaster* as queries. The microsynteny of all the species in Figure 4A was examined, except for *S. lebanonensis*, as a microsyntenic region could not be identified. To determine the phylogenetic relationship among different genes encoding elongases, we aligned the deduced amino acid sequences using the MAFFT [85] and generated nucleotide sequence alignment based on the protein sequence alignment using PAL2NAL. [86] A gene tree was inferred using IQ-TREE [87] with 1000 bootstrap replicates, and the Mortierella alpina PUFA elongase (GenBank: AF206662.1) was used as an outgroup.

To investigate the selective pressures on different elongases in the microsyteny, we estimated nonsynonymous (dN) and synonymous (dS) substitution rates using the codon-based maximum-likelihood method implemented in the CODEML program in PAML4.9. [70] Likelihood ratio tests using the chi-square approximation were used to compare nested models.

### Protein structure comparison

The protein structures of EloHL and EloF were retrieved from the AlphaFold database (AlphaFold ID: AF-Q9VH59-F1-v4 and AF-Q9VH58-F1-v4, respectively), and compared through superposition using PyMOL. [88]

### Data analysis

The production of graphs and statistical analysis was performed in RStudio version 4.2.3 (Posit Software, PBC, Boston, MA). Heatmaps were plotted using the superheat package (version 0.1.0) and all other graphs were plotted using ggplot2 (version 3.5.1). Statistical tests were performed using the package lme (version 1.1-35.4), emmeans (version 1.10.1) and basic R. Statistical tests and results for each Figure panel are listed in **Table S4**.

## ACKNOWLEDGEMENTS

Fly stocks were obtained from the Bloomington *Drosophila* Stock Center (NIH P40OD018537), the Vienna Drosophila Research Centre and the Kyoto *Drosophila* Genetic Resource Center. We acknowledge James MacRae, Jim Ellis, Vanessa Nunes and the Crick Metabolomics STP for advice and help on metabolomics experiments. We also acknowledge Eugenio Gutierrez, Joachim Kurt and the Crick fly facility for fly constructs, injection and maintenance, as well as members of the Gould lab for advice. We also thank Joe Brock for his assistance with research illustrations. This work was supported by an Investigator Award (104566) and a Technology Development Award (223760) from the Wellcome Trust and by funding from the Francis Crick Institute, which receives its core funding from Cancer Research UK (FC001088), the UK Medical Research Council (FC001088) and the Wellcome Trust (FC001088). The ToFSIMS work in this study forms part of the Life-Sciences and Health programme of the National Measurement System of the UK Department of Science, Innovation and Technology. For the purpose of Open Access, the author has applied a CC BY public copyright licence to any Author Accepted Manuscript version arising from this submission.

## DECLARATION OF INTERESTS

The authors declare no competing interests.

## Figure legends

**Figure S1:**
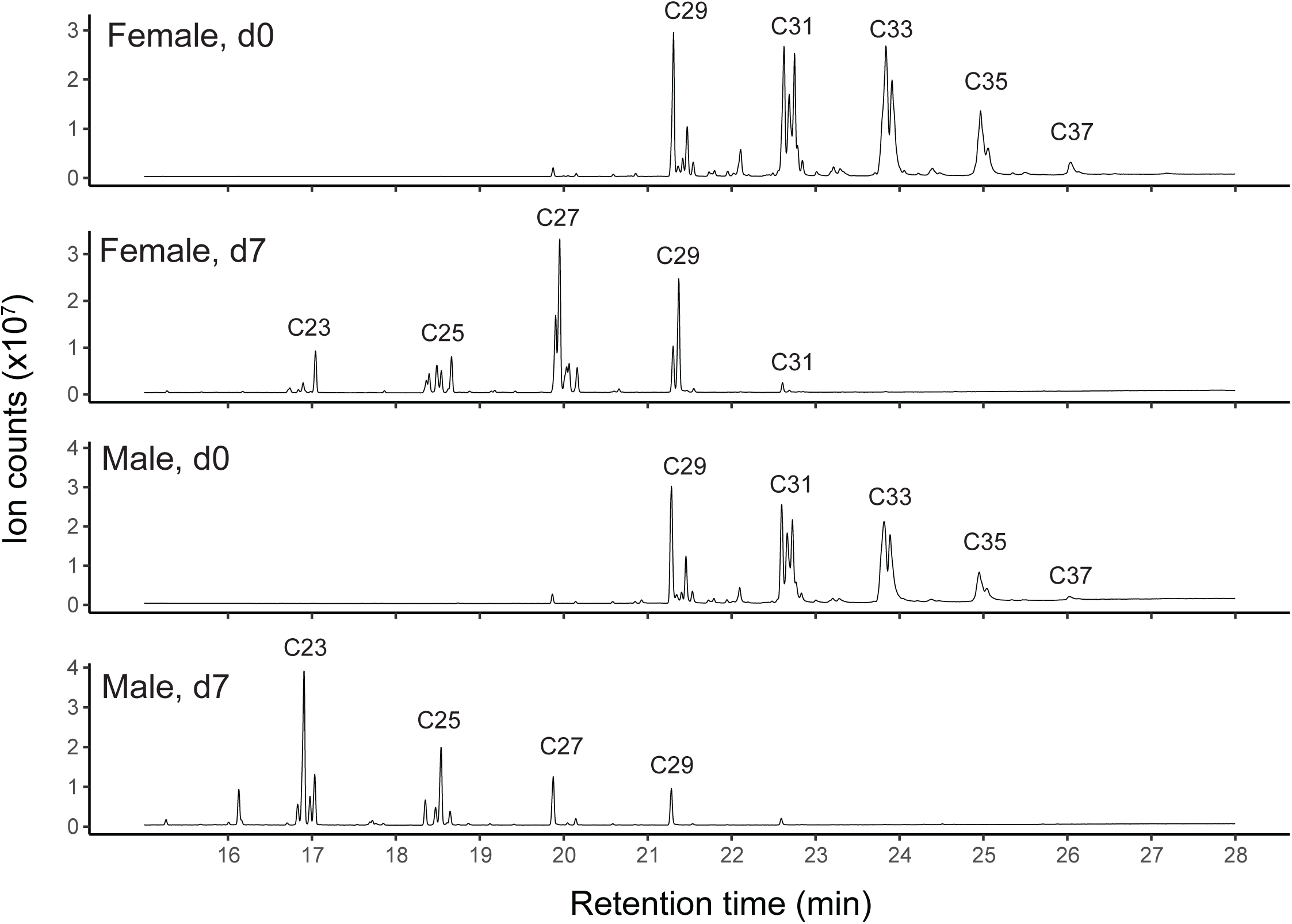
Gas chromatograms of newly emerged and 7-day old male and female flies. Representative gas chromatograms from the GC-MS analysis of newly emerged (d0) and young (d7) male and female flies. Cuticular hydrocarbon chain lengths (C23 to C37) are indicated, showing that within the first week after adult eclosion there is a transition from hyper-long to shorter hydrocarbons in both males and females.

**Figure S2:**
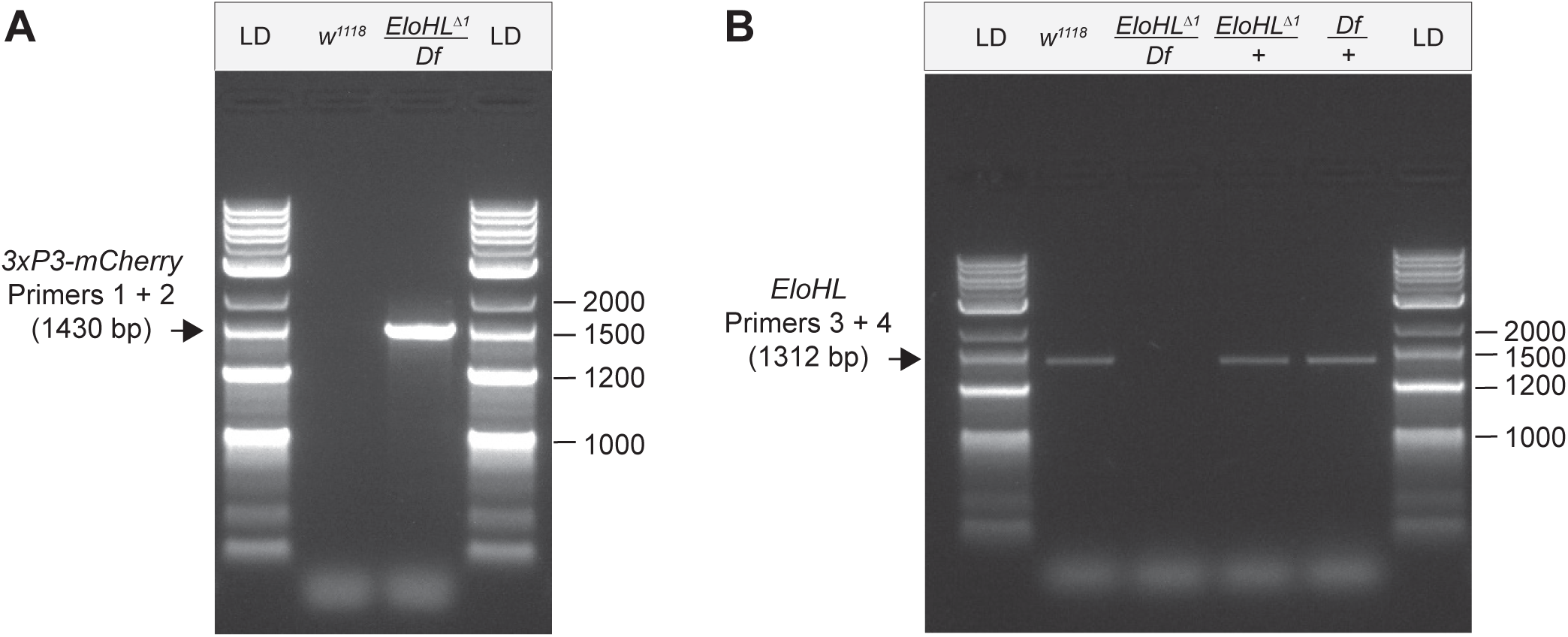
Analysis of the *EloHL*^Δ1^ mutation. The locations of Primers 1 to 4 and the strategy for replacement of the open reading frame and 3’UTR of *EloHL* (*CG8534*) with a *3xP3::mCherry* cassette are shown in Figure 3F. (A) Integration of *3xP3::mCherry* at the *EloHL* locus corresponds to a 1430 bp PCR product detected in DNA from *EloHL*^Δ1^/Df(3R)Excel6155 but not from *w^1118^ iso31* control flies. Primer 1 is located outside the 5’ homology arm and Primer 2 is located within *mCherry*. (B) Deletion of *EloHL* results in the absence of a 1312 bp PCR product in DNA from *EloHL^Δ1^*/*Df(3R)Excel6155* but not control (*w^1118^ iso31, Df(3R)Excel6155/+ or EloHL^Δ1^/+*) flies was validated using Primers 3 and 4 located within exon 1 and exon 2 of *CG8534* respectively. *EloHL*^Δ1^/Df(3R)Excel6155 lack CG8534, whereas heterozygous controls contain CG8534. The NEB 1kbp Plus DNA ladder (LD) was loaded for size estimation.

**Figure S3:**
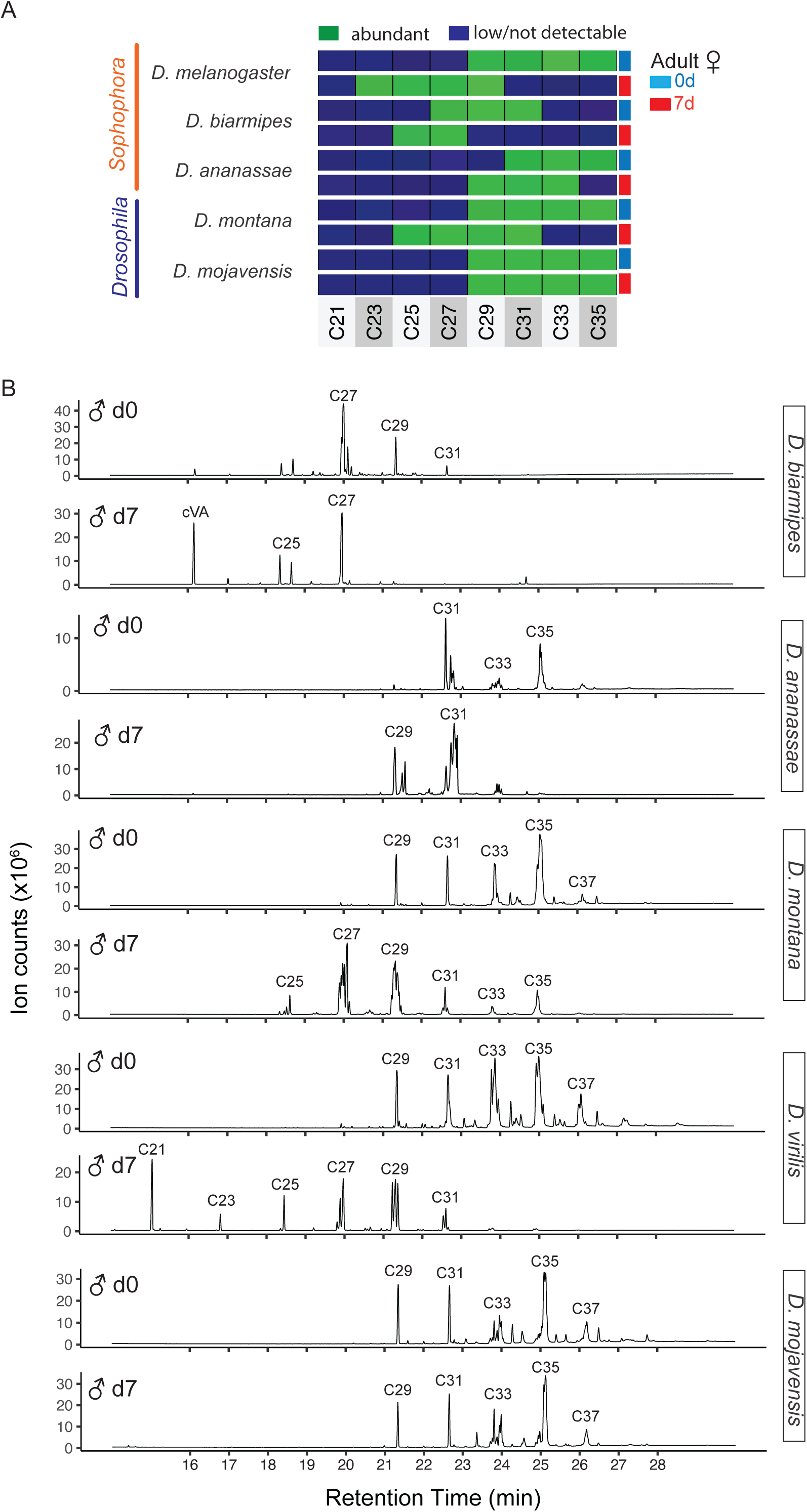
Male and female flies across many *Drosophila* species express hyper-long hydrocarbons and transition to shorter ones. (A) Abundant cuticular hydrocarbon species of newly eclosed (d0, blue) and one week old (d7, red) female flies from species of the subgenera *Sophophora* (*D. melanogaster, D. biarmipes, D. ananassae*) and *Drosophila* (*D. montana, D. mojavensis*). Abundant (green) or low/not detected (dark blue) hydrocarbons measured by HT-GC-MS are indicated according to the number of carbon atoms (C23 to C37) and pooling alkanes, monoenes, dienes and methyl-branched alkanes. Hyper-long cuticular hydrocarbons are detected across both *Sophophora* and *Drosophila* females, and *D. biarmipes* and *D. mojavensis* retain the d0 profile at d7. (B) Representative gas chromatograms of cuticular hydrocarbons from newly eclosed (d0, blue) and one week old (d7, red) male flies of the indicated species. Carbon length is indicated above peaks. These raw data were analysed for Figure 4A.

**Figure S4:**
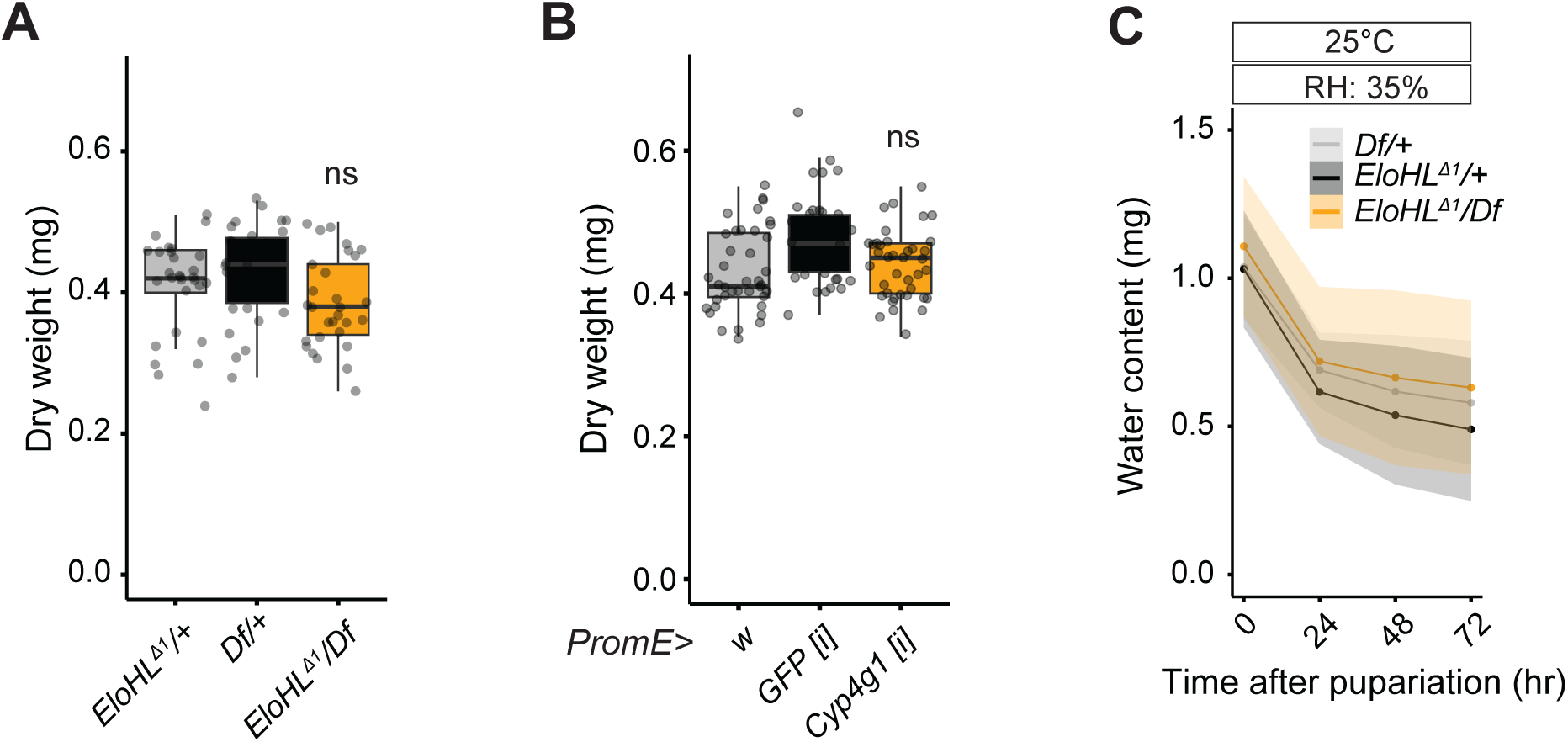
Dry weight does not significantly change in *EloHL*^Δ1^ mutant and oenocyte *Cyp4g1 [i]* pupae. (A) Pupal dry weights (mg) of *EloHL*^Δ1^/*Df(3R)Excel6155* pupa and heterozygous controls (*EloHL*^Δ1^/*+*; D*f/+)* at dark pharate stage. (N=3, n = 10) (B) Pupal dry weights (mg) of oenocyte-specific *Cyp4g1* knockdown (*PromE>Cyp4g1 [i])* and control (*PromE/+* or *PromE>GFP[i])* animals at dark pharate stage. (N=2, n ∼ 20) (C) Pupal water content (mg) versus time after pupariation for *EloHL*Δ^1^/*Df(3R)Excel6155X* mutants and heterozygous controls (*EloHL*^Δ1^/*+*; D*f/+)* at low relative humidity (35% RH) and 25°C. (N=3; n = 10)

## Notes

### Competing Interest Statement

The authors have declared no competing interest.

