## Supplemental Table S1 for "Identification of a specialized lipid barrier for *Drosophila* metamorphosis"

| Sex | Time_point | Repeat | C21:0 | C22:1 | C22:0 | 9-C23:1 | 7-C23:1 | 5-C23:1 | C23:0 | C24:1 | C24:0 | me-C24:0 | C25:2 | 9-C25:1 | 7-C25:1 | 5-C25:1 | C25:0 | C26:2 | C26:0 | me-C26:0 | C27:2 | C27:1 | C27:0 |
|---|---|---|---|---|---|---|---|---|---|---|---|---|---|---|---|---|---|---|---|---|---|---|---|
| female | 0h AE | 1.00 | 0.00 | 0.00 | 0.00 | 0.00 | 0.00 | 0.00 | 0.00 | 0.00 | 0.00 | 0.00 | 0.00 | 0.00 | 0.00 | 0.00 | 0.00 | 0.00 | 0.00 | 0.05 | 0.00 | 0.00 | 0.01 |
| female | 0h AE | 1.00 | 0.00 | 0.00 | 0.00 | 0.00 | 0.00 | 0.00 | 0.00 | 0.00 | 0.00 | 0.00 | 0.00 | 0.00 | 0.00 | 0.00 | 0.00 | 0.00 | 0.00 | 0.05 | 0.00 | 0.00 | 0.01 |
| female | 0h AE | 1.00 | 0.00 | 0.00 | 0.00 | 0.00 | 0.00 | 0.00 | 0.00 | 0.00 | 0.00 | 0.00 | 0.00 | 0.00 | 0.00 | 0.00 | 0.00 | 0.00 | 0.00 | 0.03 | 0.00 | 0.00 | 0.00 |
| female | 0h AE | 1.00 | 0.00 | 0.00 | 0.00 | 0.00 | 0.00 | 0.00 | 0.00 | 0.00 | 0.00 | 0.00 | 0.00 | 0.00 | 0.00 | 0.00 | 0.00 | 0.00 | 0.00 | 0.01 | 0.00 | 0.00 | 0.00 |
| male | 0h AE | 1.00 | 0.00 | 0.00 | 0.00 | 0.00 | 0.00 | 0.00 | 0.00 | 0.00 | 0.00 | 0.00 | 0.00 | 0.00 | 0.00 | 0.00 | 0.00 | 0.00 | 0.00 | 0.01 | 0.00 | 0.00 | 0.00 |
| male | 0h AE | 1.00 | 0.00 | 0.00 | 0.00 | 0.00 | 0.00 | 0.00 | 0.00 | 0.00 | 0.00 | 0.00 | 0.00 | 0.00 | 0.00 | 0.00 | 0.00 | 0.00 | 0.00 | 0.02 | 0.00 | 0.00 | 0.00 |
| male | 0h AE | 1.00 | 0.00 | 0.00 | 0.00 | 0.00 | 0.00 | 0.00 | 0.00 | 0.00 | 0.00 | 0.00 | 0.00 | 0.00 | 0.00 | 0.00 | 0.00 | 0.00 | 0.00 | 0.01 | 0.00 | 0.00 | 0.00 |
| male | 0h AE | 1.00 | 0.00 | 0.00 | 0.00 | 0.00 | 0.00 | 0.00 | 0.00 | 0.00 | 0.00 | 0.00 | 0.00 | 0.00 | 0.00 | 0.00 | 0.00 | 0.00 | 0.00 | 0.04 | 0.00 | 0.00 | 0.01 |
| female | 0h AE | 2.00 | 0.00 | 0.00 | 0.00 | 0.00 | 0.00 | 0.00 | 0.00 | 0.00 | 0.00 | 0.00 | 0.00 | 0.00 | 0.00 | 0.00 | 0.00 | 0.00 | 0.00 | 0.01 | 0.00 | 0.00 | 0.00 |
| female | 0h AE | 2.00 | 0.00 | 0.00 | 0.00 | 0.00 | 0.00 | 0.00 | 0.00 | 0.00 | 0.00 | 0.00 | 0.00 | 0.00 | 0.00 | 0.00 | 0.00 | 0.00 | 0.00 | 0.04 | 0.00 | 0.00 | 0.01 |
| female | 0h AE | 2.00 | 0.00 | 0.00 | 0.00 | 0.00 | 0.00 | 0.00 | 0.00 | 0.00 | 0.00 | 0.00 | 0.00 | 0.00 | 0.00 | 0.00 | 0.00 | 0.00 | 0.00 | 0.02 | 0.00 | 0.00 | 0.00 |
| female | 0h AE | 2.00 | 0.00 | 0.00 | 0.00 | 0.00 | 0.00 | 0.00 | 0.00 | 0.00 | 0.00 | 0.00 | 0.00 | 0.00 | 0.00 | 0.00 | 0.00 | 0.00 | 0.00 | 0.01 | 0.00 | 0.00 | 0.00 |
| male | 0h AE | 2.00 | 0.00 | 0.00 | 0.00 | 0.00 | 0.00 | 0.00 | 0.00 | 0.00 | 0.00 | 0.00 | 0.00 | 0.00 | 0.00 | 0.00 | 0.00 | 0.00 | 0.00 | 0.01 | 0.00 | 0.00 | 0.00 |
| male | 0h AE | 2.00 | 0.00 | 0.00 | 0.00 | 0.00 | 0.00 | 0.00 | 0.00 | 0.00 | 0.00 | 0.00 | 0.00 | 0.00 | 0.00 | 0.00 | 0.00 | 0.00 | 0.00 | 0.02 | 0.00 | 0.00 | 0.00 |
| male | 0h AE | 2.00 | 0.00 | 0.00 | 0.00 | 0.00 | 0.00 | 0.00 | 0.00 | 0.00 | 0.00 | 0.00 | 0.00 | 0.00 | 0.00 | 0.00 | 0.00 | 0.00 | 0.00 | 0.01 | 0.00 | 0.00 | 0.00 |
| female | 6h AE | 1.00 | 0.00 | 0.00 | 0.00 | 0.00 | 0.00 | 0.00 | 0.00 | 0.00 | 0.00 | 0.00 | 0.00 | 0.00 | 0.00 | 0.00 | 0.00 | 0.00 | 0.00 | 0.02 | 0.00 | 0.00 | 0.00 |
| female | 6h AE | 1.00 | 0.00 | 0.00 | 0.00 | 0.00 | 0.00 | 0.00 | 0.00 | 0.00 | 0.00 | 0.00 | 0.00 | 0.00 | 0.00 | 0.00 | 0.00 | 0.00 | 0.00 | 0.02 | 0.00 | 0.00 | 0.00 |
| female | 6h AE | 1.00 | 0.00 | 0.00 | 0.00 | 0.00 | 0.00 | 0.00 | 0.00 | 0.00 | 0.00 | 0.00 | 0.00 | 0.00 | 0.00 | 0.00 | 0.00 | 0.00 | 0.00 | 0.02 | 0.00 | 0.00 | 0.00 |
| female | 6h AE | 1.00 | 0.00 | 0.00 | 0.00 | 0.00 | 0.00 | 0.00 | 0.00 | 0.00 | 0.00 | 0.00 | 0.00 | 0.00 | 0.00 | 0.00 | 0.00 | 0.00 | 0.00 | 0.03 | 0.00 | 0.00 | 0.00 |
| male | 6h AE | 1.00 | 0.00 | 0.00 | 0.00 | 0.00 | 0.00 | 0.00 | 0.00 | 0.00 | 0.00 | 0.00 | 0.00 | 0.00 | 0.00 | 0.00 | 0.00 | 0.00 | 0.00 | 0.02 | 0.00 | 0.00 | 0.00 |
| male | 6h AE | 1.00 | 0.00 | 0.00 | 0.00 | 0.00 | 0.00 | 0.00 | 0.00 | 0.00 | 0.00 | 0.00 | 0.00 | 0.00 | 0.00 | 0.00 | 0.00 | 0.00 | 0.00 | 0.08 | 0.00 | 0.02 | 0.01 |
| male | 6h AE | 1.00 | 0.00 | 0.00 | 0.00 | 0.00 | 0.00 | 0.00 | 0.00 | 0.00 | 0.00 | 0.00 | 0.00 | 0.00 | 0.00 | 0.00 | 0.00 | 0.00 | 0.00 | 0.02 | 0.00 | 0.00 | 0.00 |
| male | 6h AE | 1.00 | 0.00 | 0.00 | 0.00 | 0.00 | 0.00 | 0.00 | 0.00 | 0.00 | 0.00 | 0.00 | 0.00 | 0.00 | 0.00 | 0.00 | 0.00 | 0.00 | 0.00 | 0.01 | 0.00 | 0.00 | 0.00 |
| female | 6h AE | 2.00 | 0.00 | 0.00 | 0.00 | 0.00 | 0.00 | 0.00 | 0.00 | 0.00 | 0.00 | 0.00 | 0.00 | 0.00 | 0.00 | 0.00 | 0.00 | 0.00 | 0.00 | 0.01 | 0.00 | 0.00 | 0.00 |
| female | 6h AE | 2.00 | 0.00 | 0.00 | 0.00 | 0.00 | 0.00 | 0.00 | 0.00 | 0.00 | 0.00 | 0.00 | 0.00 | 0.00 | 0.00 | 0.00 | 0.00 | 0.00 | 0.00 | 0.01 | 0.00 | 0.00 | 0.00 |
| female | 6h AE | 2.00 | 0.00 | 0.00 | 0.00 | 0.00 | 0.00 | 0.00 | 0.00 | 0.00 | 0.00 | 0.00 | 0.00 | 0.00 | 0.00 | 0.00 | 0.00 | 0.00 | 0.00 | 0.04 | 0.00 | 0.00 | 0.01 |
| female | 6h AE | 2.00 | 0.00 | 0.00 | 0.00 | 0.00 | 0.00 | 0.00 | 0.00 | 0.00 | 0.00 | 0.00 | 0.00 | 0.00 | 0.00 | 0.00 | 0.00 | 0.00 | 0.00 | 0.01 | 0.00 | 0.00 | 0.00 |
| male | 6h AE | 2.00 | 0.00 | 0.00 | 0.00 | 0.00 | 0.00 | 0.00 | 0.00 | 0.00 | 0.00 | 0.00 | 0.00 | 0.00 | 0.00 | 0.00 | 0.00 | 0.00 | 0.00 | 0.05 | 0.00 | 0.01 | 0.01 |
| male | 6h AE | 2.00 | 0.00 | 0.00 | 0.00 | 0.00 | 0.00 | 0.00 | 0.00 | 0.00 | 0.00 | 0.00 | 0.00 | 0.00 | 0.00 | 0.00 | 0.00 | 0.00 | 0.00 | 0.02 | 0.00 | 0.00 | 0.00 |
| male | 6h AE | 2.00 | 0.00 | 0.00 | 0.00 | 0.00 | 0.00 | 0.00 | 0.00 | 0.00 | 0.00 | 0.00 | 0.00 | 0.00 | 0.00 | 0.00 | 0.00 | 0.00 | 0.00 | 0.01 | 0.00 | 0.00 | 0.00 |
| male | 6h AE | 2.00 | 0.00 | 0.00 | 0.00 | 0.00 | 0.00 | 0.00 | 0.00 | 0.00 | 0.00 | 0.00 | 0.00 | 0.00 | 0.00 | 0.00 | 0.00 | 0.00 | 0.00 | 0.02 | 0.00 | 0.00 | 0.00 |
| female | 24h AE | 1.00 | 0.01 | 0.00 | 0.00 | 0.04 | 0.37 | 0.03 | 0.05 | 0.02 | 0.00 | 0.11 | 0.02 | 0.23 | 0.34 | 0.01 | 0.04 | 0.00 | 0.00 | 0.24 | 0.13 | 0.12 | 0.03 |
| female | 24h AE | 1.00 | 0.01 | 0.00 | 0.00 | 0.08 | 0.64 | 0.06 | 0.11 | 0.03 | 0.00 | 0.19 | 0.04 | 0.33 | 0.45 | 0.02 | 0.06 | 0.00 | 0.00 | 0.28 | 0.17 | 0.17 | 0.04 |
| female | 24h AE | 1.00 | 0.01 | 0.00 | 0.01 | 0.07 | 0.69 | 0.07 | 0.11 | 0.04 | 0.00 | 0.26 | 0.04 | 0.40 | 0.54 | 0.02 | 0.07 | 0.00 | 0.00 | 0.32 | 0.14 | 0.16 | 0.05 |
| female | 24h AE | 1.00 | 0.01 | 0.00 | 0.00 | 0.02 | 0.36 | 0.03 | 0.06 | 0.01 | 0.00 | 0.09 | 0.02 | 0.20 | 0.30 | 0.01 | 0.03 | 0.00 | 0.00 | 0.17 | 0.08 | 0.06 | 0.02 |
| male | 24h AE | 1.00 | 0.01 | 0.01 | 0.01 | 0.13 | 1.28 | 0.06 | 0.10 | 0.02 | 0.00 | 0.16 | 0.00 | 0.15 | 0.24 | 0.00 | 0.03 | 0.00 | 0.00 | 0.18 | 0.00 | 0.02 | 0.02 |
| male | 24h AE | 1.00 | 0.03 | 0.02 | 0.03 | 0.29 | 1.54 | 0.16 | 0.16 | 0.06 | 0.01 | 0.44 | 0.00 | 0.32 | 0.45 | 0.00 | 0.05 | 0.00 | 0.00 | 0.29 | 0.00 | 0.06 | 0.03 |

| Sex | Time_point | Repeat | C21:0 | C22:1 | C22:0 | 9-C23:1 | 7-C23:1 | 5-C23:1 | C23:0 | C24:1 | C24:0 | me-C24:0 | C25:2 | 9-C25:1 | 7-C25:1 | 5-C25:1 | C25:0 | C26:2 | C26:0 | me-C26:0 | C27:2 | C27:1 | C27:0 |
|---|---|---|---|---|---|---|---|---|---|---|---|---|---|---|---|---|---|---|---|---|---|---|---|
| male | 24h AE | 1.00 | 0.01 | 0.01 | 0.02 | 0.13 | 1.49 | 0.08 | 0.09 | 0.03 | 0.00 | 0.21 | 0.00 | 0.15 | 0.27 | 0.00 | 0.02 | 0.00 | 0.00 | 0.18 | 0.00 | 0.02 | 0.01 |
| male | 24h AE | 1.00 | 0.01 | 0.01 | 0.02 | 0.14 | 1.47 | 0.05 | 0.08 | 0.03 | 0.00 | 0.19 | 0.00 | 0.17 | 0.29 | 0.00 | 0.02 | 0.00 | 0.00 | 0.18 | 0.00 | 0.02 | 0.01 |
| female | 24h AE | 2.00 | 0.01 | 0.00 | 0.00 | 0.04 | 0.25 | 0.02 | 0.08 | 0.01 | 0.00 | 0.08 | 0.01 | 0.23 | 0.33 | 0.01 | 0.05 | 0.00 | 0.00 | 0.19 | 0.10 | 0.19 | 0.04 |
| female | 24h AE | 2.00 | 0.01 | 0.00 | 0.00 | 0.04 | 0.30 | 0.03 | 0.07 | 0.01 | 0.00 | 0.09 | 0.01 | 0.18 | 0.28 | 0.01 | 0.03 | 0.00 | 0.00 | 0.18 | 0.08 | 0.13 | 0.02 |
| female | 24h AE | 2.00 | 0.01 | 0.00 | 0.00 | 0.04 | 0.30 | 0.02 | 0.07 | 0.02 | 0.00 | 0.10 | 0.02 | 0.23 | 0.26 | 0.01 | 0.04 | 0.00 | 0.00 | 0.20 | 0.12 | 0.10 | 0.03 |
| female | 24h AE | 2.00 | 0.01 | 0.00 | 0.00 | 0.10 | 0.64 | 0.06 | 0.15 | 0.04 | 0.00 | 0.21 | 0.04 | 0.48 | 0.59 | 0.03 | 0.09 | 0.00 | 0.00 | 0.34 | 0.19 | 0.28 | 0.07 |
| male | 24h AE | 2.00 | 0.02 | 0.02 | 0.02 | 0.28 | 1.74 | 0.13 | 0.16 | 0.06 | 0.01 | 0.41 | 0.00 | 0.27 | 0.42 | 0.00 | 0.04 | 0.00 | 0.00 | 0.26 | 0.00 | 0.02 | 0.02 |
| male | 24h AE | 2.00 | 0.02 | 0.02 | 0.02 | 0.27 | 1.57 | 0.13 | 0.15 | 0.05 | 0.01 | 0.32 | 0.00 | 0.24 | 0.33 | 0.00 | 0.04 | 0.00 | 0.00 | 0.24 | 0.00 | 0.03 | 0.03 |
| male | 24h AE | 2.00 | 0.01 | 0.01 | 0.01 | 0.12 | 1.41 | 0.04 | 0.08 | 0.02 | 0.00 | 0.17 | 0.00 | 0.13 | 0.23 | 0.00 | 0.02 | 0.00 | 0.00 | 0.18 | 0.00 | 0.01 | 0.02 |
| male | 24h AE | 2.00 | 0.01 | 0.01 | 0.01 | 0.14 | 1.59 | 0.08 | 0.10 | 0.03 | 0.00 | 0.20 | 0.00 | 0.15 | 0.27 | 0.00 | 0.03 | 0.00 | 0.00 | 0.19 | 0.00 | 0.02 | 0.02 |
| female | 3d AE | 1.00 | 0.01 | 0.00 | 0.01 | 0.04 | 0.44 | 0.02 | 0.23 | 0.02 | 0.01 | 0.36 | 0.13 | 0.55 | 0.61 | 0.05 | 0.23 | 0.03 | 0.01 | 0.82 | 3.04 | 0.58 | 0.11 |
| female | 3d AE | 1.00 | 0.02 | 0.00 | 0.01 | 0.10 | 0.51 | 0.05 | 0.37 | 0.04 | 0.03 | 0.70 | 0.24 | 0.73 | 0.60 | 0.07 | 0.29 | 0.05 | 0.01 | 0.68 | 2.00 | 0.64 | 0.15 |
| female | 3d AE | 1.00 | 0.01 | 0.00 | 0.01 | 0.03 | 0.20 | 0.01 | 0.21 | 0.02 | 0.01 | 0.36 | 0.11 | 0.44 | 0.39 | 0.04 | 0.22 | 0.02 | 0.01 | 0.75 | 2.43 | 0.48 | 0.15 |
| female | 3d AE | 1.00 | 0.01 | 0.00 | 0.01 | 0.05 | 0.27 | 0.02 | 0.23 | 0.02 | 0.01 | 0.38 | 0.12 | 0.53 | 0.43 | 0.04 | 0.24 | 0.03 | 0.01 | 0.86 | 3.10 | 0.55 | 0.10 |
| male | 3d AE | 1.00 | 0.02 | 0.02 | 0.04 | 0.31 | 2.61 | 0.22 | 0.37 | 0.08 | 0.01 | 0.64 | 0.00 | 0.34 | 1.56 | 0.00 | 0.07 | 0.00 | 0.00 | 0.36 | 0.00 | 0.02 | 0.02 |
| male | 3d AE | 1.00 | 0.02 | 0.03 | 0.05 | 0.36 | 4.06 | 0.27 | 0.46 | 0.12 | 0.01 | 0.63 | 0.00 | 0.41 | 2.59 | 0.00 | 0.07 | 0.00 | 0.00 | 0.55 | 0.00 | 0.04 | 0.02 |
| male | 3d AE | 1.00 | 0.01 | 0.01 | 0.03 | 0.26 | 3.01 | 0.18 | 0.33 | 0.06 | 0.00 | 0.45 | 0.00 | 0.22 | 1.31 | 0.00 | 0.05 | 0.00 | 0.00 | 0.31 | 0.00 | 0.02 | 0.01 |
| male | 3d AE | 1.00 | 0.01 | 0.02 | 0.04 | 0.31 | 3.40 | 0.21 | 0.43 | 0.09 | 0.01 | 0.60 | 0.00 | 0.43 | 2.27 | 0.00 | 0.08 | 0.00 | 0.00 | 0.42 | 0.00 | 0.04 | 0.03 |
| female | 3d AE | 2.00 | 0.03 | 0.00 | 0.02 | 0.13 | 0.57 | 0.05 | 0.45 | 0.04 | 0.02 | 0.63 | 0.27 | 0.78 | 0.69 | 0.07 | 0.31 | 0.04 | 0.01 | 0.68 | 2.22 | 0.82 | 0.21 |
| female | 3d AE | 2.00 | 0.01 | 0.00 | 0.01 | 0.05 | 0.32 | 0.03 | 0.34 | 0.01 | 0.01 | 0.36 | 0.14 | 0.48 | 0.43 | 0.04 | 0.17 | 0.03 | 0.01 | 0.71 | 2.69 | 0.48 | 0.10 |
| female | 3d AE | 2.00 | 0.01 | 0.00 | 0.01 | 0.04 | 0.26 | 0.02 | 0.26 | 0.02 | 0.01 | 0.29 | 0.12 | 0.49 | 0.51 | 0.05 | 0.22 | 0.02 | 0.01 | 0.67 | 2.58 | 0.64 | 0.11 |
| female | 3d AE | 2.00 | 0.02 | 0.00 | 0.01 | 0.08 | 0.44 | 0.04 | 0.28 | 0.03 | 0.02 | 0.42 | 0.18 | 0.68 | 0.73 | 0.07 | 0.24 | 0.04 | 0.01 | 0.93 | 3.77 | 0.85 | 0.15 |
| male | 3d AE | 2.00 | 0.02 | 0.02 | 0.07 | 0.35 | 3.69 | 0.24 | 0.44 | 0.09 | 0.01 | 0.63 | 0.00 | 0.35 | 1.66 | 0.00 | 0.07 | 0.00 | 0.00 | 0.37 | 0.00 | 0.02 | 0.02 |
| male | 3d AE | 2.00 | 0.07 | 0.10 | 0.07 | 0.84 | 2.77 | 0.55 | 0.62 | 0.36 | 0.03 | 1.69 | 0.00 | 0.84 | 2.18 | 0.00 | 0.19 | 0.00 | 0.01 | 0.65 | 0.00 | 0.08 | 0.05 |
| male | 3d AE | 2.00 | 0.04 | 0.05 | 0.07 | 0.56 | 2.48 | 0.36 | 0.49 | 0.21 | 0.02 | 1.07 | 0.00 | 0.52 | 1.73 | 0.00 | 0.12 | 0.00 | 0.01 | 0.60 | 0.00 | 0.05 | 0.05 |
| male | 3d AE | 2.00 | 0.02 | 0.02 | 0.06 | 0.37 | 3.93 | 0.27 | 0.48 | 0.14 | 0.01 | 0.72 | 0.00 | 0.45 | 2.29 | 0.00 | 0.08 | 0.00 | 0.00 | 0.42 | 0.00 | 0.04 | 0.02 |
| female | 7d AE | 1.00 | 0.01 | 0.00 | 0.01 | 0.03 | 0.13 | 0.01 | 0.34 | 0.01 | 0.01 | 0.29 | 0.15 | 0.28 | 0.22 | 0.04 | 0.26 | 0.03 | 0.01 | 0.91 | 3.14 | 0.34 | 0.14 |
| female | 7d AE | 1.00 | 0.01 | 0.00 | 0.01 | 0.03 | 0.13 | 0.02 | 0.37 | 0.01 | 0.01 | 0.32 | 0.18 | 0.28 | 0.22 | 0.04 | 0.23 | 0.03 | 0.01 | 0.73 | 2.50 | 0.30 | 0.16 |
| female | 7d AE | 1.00 | 0.01 | 0.00 | 0.01 | 0.04 | 0.15 | 0.02 | 0.43 | 0.01 | 0.01 | 0.32 | 0.18 | 0.30 | 0.26 | 0.04 | 0.29 | 0.02 | 0.01 | 0.80 | 2.56 | 0.33 | 0.15 |
| female | 7d AE | 1.00 | 0.02 | 0.00 | 0.02 | 0.06 | 0.24 | 0.03 | 0.47 | 0.02 | 0.02 | 0.52 | 0.27 | 0.44 | 0.37 | 0.08 | 0.36 | 0.04 | 0.01 | 0.70 | 2.03 | 0.42 | 0.22 |
| male | 7d AE | 1.00 | 0.04 | 0.04 | 0.04 | 0.28 | 5.20 | 0.42 | 0.54 | 0.12 | 0.01 | 0.79 | 0.00 | 0.20 | 1.58 | 0.00 | 0.08 | 0.00 | 0.00 | 0.52 | 0.00 | 0.02 | 0.03 |
| male | 7d AE | 1.00 | 0.03 | 0.02 | 0.08 | 0.27 | 4.41 | 0.38 | 0.63 | 0.10 | 0.01 | 0.83 | 0.00 | 0.22 | 1.65 | 0.00 | 0.09 | 0.00 | 0.00 | 0.48 | 0.00 | 0.02 | 0.03 |
| male | 7d AE | 1.00 | 0.03 | 0.04 | 0.06 | 0.26 | 4.52 | 0.40 | 0.57 | 0.09 | 0.01 | 0.68 | 0.00 | 0.19 | 1.69 | 0.00 | 0.07 | 0.00 | 0.00 | 0.48 | 0.00 | 0.02 | 0.03 |
| male | 7d AE | 1.00 | 0.05 | 0.03 | 0.08 | 0.29 | 4.61 | 0.45 | 0.67 | 0.11 | 0.01 | 0.78 | 0.00 | 0.26 | 1.51 | 0.00 | 0.09 | 0.00 | 0.00 | 0.49 | 0.00 | 0.02 | 0.04 |
| female | 7d AE | 2.00 | 0.02 | 0.00 | 0.01 | 0.05 | 0.21 | 0.03 | 0.45 | 0.01 | 0.02 | 0.36 | 0.23 | 0.37 | 0.35 | 0.07 | 0.29 | 0.04 | 0.01 | 1.09 | 3.80 | 0.41 | 0.16 |
| female | 7d AE | 2.00 | 0.02 | 0.00 | 0.01 | 0.04 | 0.22 | 0.02 | 0.48 | 0.01 | 0.01 | 0.48 | 0.26 | 0.51 | 0.43 | 0.08 | 0.41 | 0.03 | 0.01 | 1.22 | 4.30 | 0.55 | 0.21 |
| female | 7d AE | 2.00 | 0.01 | 0.00 | 0.01 | 0.03 | 0.14 | 0.01 | 0.35 | 0.01 | 0.01 | 0.29 | 0.18 | 0.35 | 0.26 | 0.04 | 0.25 | 0.02 | 0.01 | 0.79 | 3.15 | 0.40 | 0.13 |

| Sex | Time_point | Repeat | C21:0 | C22:1 | C22:0 | 9-C23:1 | 7-C23:1 | 5-C23:1 | C23:0 | C24:1 | C24:0 | me-C24:0 | C25:2 | 9-C25:1 | 7-C25:1 | 5-C25:1 | C25:0 | C26:2 | C26:0 | me-C26:0 | C27:2 | C27:1 | C27:0 |
|---|---|---|---|---|---|---|---|---|---|---|---|---|---|---|---|---|---|---|---|---|---|---|---|
| female | 7d AE | 2.00 | 0.01 | 0.00 | 0.01 | 0.05 | 0.30 | 0.03 | 0.41 | 0.02 | 0.01 | 0.38 | 0.23 | 0.31 | 0.41 | 0.06 | 0.26 | 0.03 | 0.01 | 0.96 | 3.46 | 0.36 | 0.13 |
| male | 7d AE | 2.00 | 0.07 | 0.08 | 0.07 | 0.41 | 2.89 | 0.53 | 0.54 | 0.18 | 0.02 | 1.15 | 0.00 | 0.19 | 1.04 | 0.00 | 0.10 | 0.00 | 0.01 | 0.51 | 0.00 | 0.02 | 0.04 |
| male | 7d AE | 2.00 | 0.03 | 0.04 | 0.04 | 0.26 | 3.88 | 0.33 | 0.53 | 0.10 | 0.01 | 0.62 | 0.00 | 0.18 | 1.45 | 0.00 | 0.07 | 0.00 | 0.00 | 0.44 | 0.00 | 0.02 | 0.03 |
| male | 7d AE | 2.00 | 0.04 | 0.04 | 0.05 | 0.34 | 5.68 | 0.37 | 0.55 | 0.10 | 0.01 | 0.70 | 0.00 | 0.16 | 1.57 | 0.00 | 0.07 | 0.00 | 0.00 | 0.56 | 0.00 | 0.02 | 0.03 |
| male | 7d AE | 2.00 | 0.03 | 0.03 | 0.07 | 0.28 | 5.07 | 0.34 | 0.55 | 0.09 | 0.01 | 0.55 | 0.00 | 0.13 | 1.36 | 0.00 | 0.08 | 0.00 | 0.00 | 0.47 | 0.00 | 0.01 | 0.03 |
| mixed | wL3 | 1.00 | 0.00 | 0.00 | 0.00 | 0.00 | 0.00 | 0.00 | 0.00 | 0.00 | 0.00 | 0.00 | 0.00 | 0.00 | 0.00 | 0.00 | 0.00 | 0.00 | 0.00 | 0.00 | 0.00 | 0.00 | 0.00 |
| mixed | wL3 | 1.00 | 0.00 | 0.00 | 0.00 | 0.00 | 0.00 | 0.00 | 0.00 | 0.00 | 0.00 | 0.00 | 0.00 | 0.00 | 0.00 | 0.00 | 0.00 | 0.00 | 0.00 | 0.00 | 0.00 | 0.00 | 0.00 |
| mixed | wL3 | 1.00 | 0.00 | 0.00 | 0.00 | 0.00 | 0.00 | 0.00 | 0.00 | 0.00 | 0.00 | 0.00 | 0.00 | 0.00 | 0.00 | 0.00 | 0.00 | 0.00 | 0.00 | 0.00 | 0.00 | 0.00 | 0.00 |
| mixed | wL3 | 1.00 | 0.00 | 0.00 | 0.00 | 0.00 | 0.00 | 0.00 | 0.00 | 0.00 | 0.00 | 0.00 | 0.00 | 0.00 | 0.00 | 0.00 | 0.00 | 0.00 | 0.00 | 0.00 | 0.00 | 0.00 | 0.00 |
| mixed | wL3 | 2.00 | 0.00 | 0.00 | 0.00 | 0.00 | 0.00 | 0.00 | 0.00 | 0.00 | 0.00 | 0.00 | 0.00 | 0.00 | 0.00 | 0.00 | 0.00 | 0.00 | 0.00 | 0.00 | 0.00 | 0.00 | 0.00 |
| mixed | wL3 | 2.00 | 0.00 | 0.00 | 0.00 | 0.00 | 0.00 | 0.00 | 0.00 | 0.00 | 0.00 | 0.00 | 0.00 | 0.00 | 0.00 | 0.00 | 0.00 | 0.00 | 0.00 | 0.00 | 0.00 | 0.00 | 0.00 |
| mixed | wL3 | 2.00 | 0.00 | 0.00 | 0.00 | 0.00 | 0.00 | 0.00 | 0.00 | 0.00 | 0.00 | 0.00 | 0.00 | 0.00 | 0.00 | 0.00 | 0.00 | 0.00 | 0.00 | 0.00 | 0.00 | 0.00 | 0.00 |
| mixed | wL3 | 2.00 | 0.00 | 0.00 | 0.00 | 0.00 | 0.00 | 0.00 | 0.00 | 0.00 | 0.00 | 0.00 | 0.00 | 0.00 | 0.00 | 0.00 | 0.00 | 0.00 | 0.00 | 0.00 | 0.00 | 0.00 | 0.00 |
| mixed | 48h_APF | 1.00 | 0.00 | 0.00 | 0.00 | 0.00 | 0.00 | 0.00 | 0.00 | 0.00 | 0.00 | 0.00 | 0.00 | 0.00 | 0.00 | 0.00 | 0.00 | 0.00 | 0.00 | 0.00 | 0.00 | 0.00 | 0.00 |
| mixed | 48h_APF | 1.00 | 0.00 | 0.00 | 0.00 | 0.00 | 0.00 | 0.00 | 0.00 | 0.00 | 0.00 | 0.00 | 0.00 | 0.00 | 0.00 | 0.00 | 0.00 | 0.00 | 0.00 | 0.00 | 0.00 | 0.00 | 0.00 |
| mixed | 48h_APF | 1.00 | 0.00 | 0.00 | 0.00 | 0.00 | 0.00 | 0.00 | 0.00 | 0.00 | 0.00 | 0.00 | 0.00 | 0.00 | 0.00 | 0.00 | 0.00 | 0.00 | 0.00 | 0.00 | 0.00 | 0.00 | 0.00 |
| mixed | 48h_APF | 1.00 | 0.00 | 0.00 | 0.00 | 0.00 | 0.00 | 0.00 | 0.00 | 0.00 | 0.00 | 0.00 | 0.00 | 0.00 | 0.00 | 0.00 | 0.00 | 0.00 | 0.00 | 0.00 | 0.00 | 0.00 | 0.00 |
| mixed | 48h_APF | 2.00 | 0.00 | 0.00 | 0.00 | 0.00 | 0.00 | 0.00 | 0.00 | 0.00 | 0.00 | 0.00 | 0.00 | 0.00 | 0.00 | 0.00 | 0.00 | 0.00 | 0.00 | 0.00 | 0.00 | 0.00 | 0.00 |
| mixed | 48h_APF | 2.00 | 0.00 | 0.00 | 0.00 | 0.00 | 0.00 | 0.00 | 0.00 | 0.00 | 0.00 | 0.00 | 0.00 | 0.00 | 0.00 | 0.00 | 0.00 | 0.00 | 0.00 | 0.00 | 0.00 | 0.00 | 0.00 |
| mixed | 48h_APF | 2.00 | 0.00 | 0.00 | 0.00 | 0.00 | 0.00 | 0.00 | 0.00 | 0.00 | 0.00 | 0.00 | 0.00 | 0.00 | 0.00 | 0.00 | 0.00 | 0.00 | 0.00 | 0.00 | 0.00 | 0.00 | 0.00 |
| mixed | 48h_APF | 2.00 | 0.00 | 0.00 | 0.00 | 0.00 | 0.00 | 0.00 | 0.00 | 0.00 | 0.00 | 0.00 | 0.00 | 0.00 | 0.00 | 0.00 | 0.00 | 0.00 | 0.00 | 0.00 | 0.00 | 0.00 | 0.00 |

| Sex | Time_point | Repeat | me-C28:0 | C29:2 | C29:1 (A) | C29:1 (B) | C29:1 (C) | C29:0 | C30:1 | me-C30:0 | C31:2 | C31:1 (A) | C31:1 (B) | C31:1 (C) | C31:0 | me-C32:0 | C32:2 | C32:1 | C33:2 | C33:1 | C35:2 | C35:1 | C37:2 |
|---|---|---|---|---|---|---|---|---|---|---|---|---|---|---|---|---|---|---|---|---|---|---|---|
| female | 0h AE | 1.00 | 2.30 | 0.00 | 0.05 | 0.07 | 0.31 | 0.03 | 0.19 | 0.47 | 0.20 | 0.41 | 0.58 | 0.10 | 0.03 | 0.15 | 0.10 | 0.06 | 1.94 | 0.64 | 0.97 | 0.12 | 0.14 |
| female | 0h AE | 1.00 | 2.26 | 0.00 | 0.07 | 0.05 | 0.25 | 0.03 | 0.13 | 0.49 | 0.18 | 0.44 | 0.53 | 0.09 | 0.02 | 0.16 | 0.10 | 0.05 | 1.99 | 0.62 | 0.85 | 0.13 | 0.11 |
| female | 0h AE | 1.00 | 1.77 | 0.00 | 0.03 | 0.03 | 0.15 | 0.01 | 0.08 | 0.35 | 0.11 | 0.24 | 0.33 | 0.05 | 0.01 | 0.08 | 0.04 | 0.02 | 1.05 | 0.32 | 0.33 | 0.06 | 0.05 |
| female | 0h AE | 1.00 | 1.62 | 0.00 | 0.02 | 0.02 | 0.12 | 0.01 | 0.05 | 0.32 | 0.11 | 0.24 | 0.28 | 0.05 | 0.01 | 0.09 | 0.03 | 0.02 | 0.96 | 0.29 | 0.39 | 0.06 | 0.04 |
| male | 0h AE | 1.00 | 1.31 | 0.00 | 0.02 | 0.02 | 0.10 | 0.01 | 0.05 | 0.23 | 0.10 | 0.19 | 0.20 | 0.03 | 0.01 | 0.09 | 0.03 | 0.02 | 0.84 | 0.22 | 0.24 | 0.03 | 0.03 |
| male | 0h AE | 1.00 | 1.59 | 0.00 | 0.02 | 0.03 | 0.13 | 0.02 | 0.06 | 0.26 | 0.10 | 0.22 | 0.26 | 0.06 | 0.01 | 0.07 | 0.03 | 0.02 | 0.91 | 0.24 | 0.31 | 0.04 | 0.04 |
| male | 0h AE | 1.00 | 1.28 | 0.00 | 0.01 | 0.02 | 0.08 | 0.01 | 0.04 | 0.21 | 0.09 | 0.15 | 0.21 | 0.02 | 0.01 | 0.06 | 0.01 | 0.01 | 0.65 | 0.20 | 0.24 | 0.03 | 0.02 |
| male | 0h AE | 1.00 | 1.79 | 0.00 | 0.04 | 0.06 | 0.23 | 0.02 | 0.11 | 0.33 | 0.19 | 0.35 | 0.40 | 0.06 | 0.02 | 0.10 | 0.07 | 0.03 | 1.43 | 0.44 | 0.60 | 0.07 | 0.10 |
| female | 0h AE | 2.00 | 0.99 | 0.00 | 0.01 | 0.01 | 0.08 | 0.01 | 0.04 | 0.23 | 0.06 | 0.16 | 0.26 | 0.04 | 0.01 | 0.07 | 0.02 | 0.01 | 0.72 | 0.23 | 0.33 | 0.05 | 0.04 |
| female | 0h AE | 2.00 | 1.82 | 0.00 | 0.06 | 0.06 | 0.29 | 0.04 | 0.15 | 0.39 | 0.18 | 0.42 | 0.58 | 0.08 | 0.02 | 0.13 | 0.11 | 0.05 | 1.92 | 0.51 | 0.77 | 0.09 | 0.17 |
| female | 0h AE | 2.00 | 1.46 | 0.00 | 0.02 | 0.02 | 0.14 | 0.02 | 0.09 | 0.36 | 0.09 | 0.29 | 0.40 | 0.08 | 0.02 | 0.14 | 0.05 | 0.03 | 1.42 | 0.44 | 0.51 | 0.08 | 0.08 |
| female | 0h AE | 2.00 | 1.93 | 0.00 | 0.02 | 0.03 | 0.15 | 0.01 | 0.07 | 0.45 | 0.13 | 0.32 | 0.45 | 0.04 | 0.01 | 0.14 | 0.05 | 0.03 | 1.45 | 0.45 | 0.52 | 0.06 | 0.07 |
| male | 0h AE | 2.00 | 1.19 | 0.00 | 0.01 | 0.01 | 0.08 | 0.01 | 0.04 | 0.22 | 0.07 | 0.14 | 0.21 | 0.03 | 0.01 | 0.06 | 0.03 | 0.01 | 0.71 | 0.21 | 0.26 | 0.03 | 0.03 |
| male | 0h AE | 2.00 | 1.27 | 0.00 | 0.01 | 0.02 | 0.11 | 0.01 | 0.05 | 0.27 | 0.07 | 0.18 | 0.28 | 0.05 | 0.01 | 0.08 | 0.03 | 0.02 | 0.82 | 0.28 | 0.35 | 0.06 | 0.05 |
| male | 0h AE | 2.00 | 0.98 | 0.00 | 0.01 | 0.01 | 0.09 | 0.01 | 0.03 | 0.18 | 0.06 | 0.14 | 0.18 | 0.03 | 0.01 | 0.05 | 0.02 | 0.01 | 0.64 | 0.17 | 0.19 | 0.03 | 0.03 |
| female | 6h AE | 1.00 | 2.37 | 0.00 | 0.07 | 0.07 | 0.23 | 0.03 | 0.11 | 0.46 | 0.27 | 0.42 | 0.41 | 0.04 | 0.02 | 0.10 | 0.08 | 0.03 | 1.54 | 0.37 | 0.55 | 0.07 | 0.07 |
| female | 6h AE | 1.00 | 1.69 | 0.00 | 0.03 | 0.05 | 0.17 | 0.02 | 0.07 | 0.38 | 0.16 | 0.30 | 0.30 | 0.04 | 0.01 | 0.10 | 0.05 | 0.02 | 1.04 | 0.26 | 0.36 | 0.05 | 0.05 |
| female | 6h AE | 1.00 | 1.74 | 0.00 | 0.04 | 0.05 | 0.18 | 0.01 | 0.07 | 0.33 | 0.15 | 0.36 | 0.27 | 0.04 | 0.01 | 0.09 | 0.06 | 0.03 | 1.01 | 0.31 | 0.36 | 0.04 | 0.06 |
| female | 6h AE | 1.00 | 1.78 | 0.00 | 0.05 | 0.06 | 0.17 | 0.02 | 0.07 | 0.35 | 0.16 | 0.35 | 0.29 | 0.04 | 0.01 | 0.10 | 0.06 | 0.03 | 1.10 | 0.32 | 0.36 | 0.05 | 0.05 |
| male | 6h AE | 1.00 | 1.55 | 0.00 | 0.05 | 0.03 | 0.15 | 0.01 | 0.05 | 0.26 | 0.20 | 0.25 | 0.22 | 0.04 | 0.01 | 0.06 | 0.03 | 0.01 | 0.85 | 0.28 | 0.31 | 0.04 | 0.03 |
| male | 6h AE | 1.00 | 2.30 | 0.00 | 0.15 | 0.14 | 0.46 | 0.03 | 0.21 | 0.45 | 0.43 | 0.59 | 0.50 | 0.07 | 0.02 | 0.11 | 0.16 | 0.05 | 2.34 | 0.66 | 0.99 | 0.11 | 0.16 |
| male | 6h AE | 1.00 | 1.61 | 0.00 | 0.04 | 0.04 | 0.18 | 0.01 | 0.05 | 0.26 | 0.18 | 0.25 | 0.24 | 0.03 | 0.01 | 0.07 | 0.04 | 0.02 | 1.00 | 0.26 | 0.32 | 0.04 | 0.05 |
| male | 6h AE | 1.00 | 1.41 | 0.00 | 0.03 | 0.03 | 0.15 | 0.01 | 0.06 | 0.27 | 0.15 | 0.25 | 0.27 | 0.03 | 0.01 | 0.06 | 0.05 | 0.01 | 0.78 | 0.19 | 0.30 | 0.03 | 0.04 |
| female | 6h AE | 2.00 | 1.83 | 0.00 | 0.05 | 0.04 | 0.24 | 0.02 | 0.07 | 0.44 | 0.18 | 0.42 | 0.42 | 0.06 | 0.01 | 0.13 | 0.06 | 0.03 | 1.51 | 0.42 | 0.49 | 0.06 | 0.07 |
| female | 6h AE | 2.00 | 1.99 | 0.00 | 0.04 | 0.04 | 0.18 | 0.01 | 0.06 | 0.38 | 0.16 | 0.27 | 0.29 | 0.05 | 0.01 | 0.10 | 0.04 | 0.02 | 1.06 | 0.30 | 0.50 | 0.06 | 0.05 |
| female | 6h AE | 2.00 | 2.02 | 0.00 | 0.07 | 0.08 | 0.31 | 0.02 | 0.12 | 0.40 | 0.18 | 0.38 | 0.44 | 0.07 | 0.01 | 0.16 | 0.08 | 0.03 | 1.67 | 0.46 | 0.71 | 0.08 | 0.10 |
| female | 6h AE | 2.00 | 1.64 | 0.00 | 0.06 | 0.05 | 0.18 | 0.01 | 0.09 | 0.38 | 0.16 | 0.29 | 0.33 | 0.03 | 0.02 | 0.10 | 0.04 | 0.02 | 1.08 | 0.34 | 0.44 | 0.07 | 0.07 |
| male | 6h AE | 2.00 | 1.80 | 0.00 | 0.11 | 0.10 | 0.35 | 0.03 | 0.15 | 0.34 | 0.29 | 0.41 | 0.43 | 0.06 | 0.02 | 0.10 | 0.09 | 0.04 | 1.48 | 0.40 | 0.65 | 0.09 | 0.13 |
| male | 6h AE | 2.00 | 1.39 | 0.00 | 0.05 | 0.05 | 0.17 | 0.01 | 0.06 | 0.23 | 0.15 | 0.22 | 0.22 | 0.04 | 0.01 | 0.06 | 0.05 | 0.02 | 0.86 | 0.23 | 0.32 | 0.04 | 0.04 |
| male | 6h AE | 2.00 | 1.33 | 0.00 | 0.04 | 0.04 | 0.15 | 0.01 | 0.05 | 0.24 | 0.14 | 0.20 | 0.22 | 0.03 | 0.01 | 0.06 | 0.03 | 0.01 | 0.75 | 0.23 | 0.28 | 0.03 | 0.03 |
| male | 6h AE | 2.00 | 1.46 | 0.00 | 0.03 | 0.04 | 0.14 | 0.01 | 0.05 | 0.21 | 0.14 | 0.19 | 0.20 | 0.03 | 0.01 | 0.06 | 0.03 | 0.01 | 0.69 | 0.20 | 0.23 | 0.05 | 0.03 |
| female | 24h AE | 1.00 | 2.67 | 0.00 | 0.13 | 0.05 | 0.17 | 0.02 | 0.05 | 0.44 | 0.21 | 0.26 | 0.20 | 0.04 | 0.01 | 0.10 | 0.05 | 0.02 | 0.92 | 0.21 | 0.34 | 0.03 | 0.04 |
| female | 24h AE | 1.00 | 2.65 | 0.00 | 0.18 | 0.07 | 0.24 | 0.03 | 0.07 | 0.49 | 0.30 | 0.40 | 0.32 | 0.07 | 0.02 | 0.12 | 0.09 | 0.03 | 1.06 | 0.30 | 0.38 | 0.04 | 0.06 |
| female | 24h AE | 1.00 | 2.32 | 0.00 | 0.19 | 0.12 | 0.30 | 0.04 | 0.14 | 0.41 | 0.26 | 0.39 | 0.33 | 0.06 | 0.02 | 0.13 | 0.09 | 0.03 | 1.26 | 0.35 | 0.49 | 0.07 | 0.11 |
| female | 24h AE | 1.00 | 2.13 | 0.00 | 0.09 | 0.07 | 0.16 | 0.02 | 0.05 | 0.32 | 0.15 | 0.24 | 0.19 | 0.03 | 0.01 | 0.08 | 0.03 | 0.01 | 0.64 | 0.18 | 0.26 | 0.04 | 0.04 |
| male | 24h AE | 1.00 | 1.77 | 0.00 | 0.05 | 0.03 | 0.13 | 0.01 | 0.03 | 0.21 | 0.12 | 0.14 | 0.13 | 0.02 | 0.01 | 0.05 | 0.02 | 0.01 | 0.43 | 0.14 | 0.13 | 0.02 | 0.02 |
| male | 24h AE | 1.00 | 2.41 | 0.00 | 0.10 | 0.09 | 0.28 | 0.02 | 0.08 | 0.34 | 0.23 | 0.29 | 0.25 | 0.05 | 0.02 | 0.09 | 0.06 | 0.02 | 0.96 | 0.25 | 0.31 | 0.03 | 0.06 |

| Sex | Time_point | Repeat | me-C28:0 | C29:2 | C29:1 (A) | C29:1 (B) | C29:1 (C) | C29:0 | C30:1 | me-C30:0 | C31:2 | C31:1 (A) | C31:1 (B) | C31:1 (C) | C31:0 | me-C32:0 | C32:2 | C32:1 | C33:2 | C33:1 | C35:2 | C35:1 | C37:2 |
|---|---|---|---|---|---|---|---|---|---|---|---|---|---|---|---|---|---|---|---|---|---|---|---|
| male | 24h AE | 1.00 | 1.77 | 0.00 | 0.04 | 0.04 | 0.12 | 0.00 | 0.03 | 0.17 | 0.10 | 0.11 | 0.12 | 0.01 | 0.01 | 0.05 | 0.02 | 0.01 | 0.44 | 0.11 | 0.16 | 0.02 | 0.02 |
| male | 24h AE | 1.00 | 1.83 | 0.00 | 0.04 | 0.04 | 0.12 | 0.01 | 0.04 | 0.23 | 0.12 | 0.13 | 0.14 | 0.03 | 0.01 | 0.06 | 0.02 | 0.01 | 0.46 | 0.12 | 0.17 | 0.02 | 0.03 |
| female | 24h AE | 2.00 | 2.22 | 0.00 | 0.13 | 0.06 | 0.18 | 0.02 | 0.05 | 0.37 | 0.22 | 0.26 | 0.25 | 0.04 | 0.01 | 0.09 | 0.03 | 0.01 | 0.69 | 0.21 | 0.29 | 0.03 | 0.03 |
| female | 24h AE | 2.00 | 2.35 | 0.00 | 0.11 | 0.06 | 0.18 | 0.02 | 0.05 | 0.40 | 0.22 | 0.28 | 0.24 | 0.02 | 0.01 | 0.09 | 0.06 | 0.02 | 0.89 | 0.24 | 0.29 | 0.03 | 0.04 |
| female | 24h AE | 2.00 | 2.32 | 0.00 | 0.09 | 0.05 | 0.16 | 0.02 | 0.07 | 0.31 | 0.16 | 0.20 | 0.21 | 0.03 | 0.01 | 0.08 | 0.05 | 0.02 | 0.71 | 0.19 | 0.27 | 0.04 | 0.04 |
| female | 24h AE | 2.00 | 2.56 | 0.00 | 0.20 | 0.11 | 0.29 | 0.04 | 0.12 | 0.50 | 0.28 | 0.46 | 0.48 | 0.05 | 0.02 | 0.16 | 0.08 | 0.03 | 1.26 | 0.37 | 0.50 | 0.06 | 0.09 |
| male | 24h AE | 2.00 | 2.09 | 0.00 | 0.08 | 0.05 | 0.17 | 0.01 | 0.07 | 0.29 | 0.15 | 0.19 | 0.19 | 0.03 | 0.01 | 0.09 | 0.04 | 0.02 | 0.66 | 0.17 | 0.30 | 0.04 | 0.04 |
| male | 24h AE | 2.00 | 2.30 | 0.00 | 0.07 | 0.06 | 0.19 | 0.02 | 0.05 | 0.27 | 0.16 | 0.19 | 0.20 | 0.03 | 0.01 | 0.08 | 0.03 | 0.02 | 0.62 | 0.16 | 0.25 | 0.05 | 0.04 |
| male | 24h AE | 2.00 | 1.83 | 0.00 | 0.05 | 0.03 | 0.11 | 0.02 | 0.04 | 0.23 | 0.09 | 0.11 | 0.13 | 0.02 | 0.01 | 0.06 | 0.02 | 0.00 | 0.40 | 0.11 | 0.17 | 0.03 | 0.01 |
| male | 24h AE | 2.00 | 2.28 | 0.00 | 0.05 | 0.04 | 0.13 | 0.01 | 0.05 | 0.24 | 0.12 | 0.13 | 0.13 | 0.02 | 0.01 | 0.06 | 0.03 | 0.00 | 0.47 | 0.12 | 0.19 | 0.02 | 0.02 |
| female | 3d AE | 1.00 | 2.22 | 1.57 | 0.00 | 0.03 | 0.08 | 0.01 | 0.01 | 0.18 | 0.08 | 0.06 | 0.06 | 0.01 | 0.01 | 0.04 | 0.01 | 0.00 | 0.22 | 0.06 | 0.08 | 0.00 | 0.00 |
| female | 3d AE | 1.00 | 2.27 | 1.77 | 0.00 | 0.04 | 0.09 | 0.02 | 0.02 | 0.18 | 0.06 | 0.06 | 0.06 | 0.02 | 0.01 | 0.06 | 0.01 | 0.00 | 0.21 | 0.05 | 0.06 | 0.00 | 0.00 |
| female | 3d AE | 1.00 | 1.84 | 1.55 | 0.00 | 0.03 | 0.06 | 0.01 | 0.01 | 0.16 | 0.05 | 0.05 | 0.05 | 0.01 | 0.01 | 0.04 | 0.01 | 0.00 | 0.20 | 0.06 | 0.06 | 0.00 | 0.00 |
| female | 3d AE | 1.00 | 1.87 | 1.65 | 0.00 | 0.03 | 0.06 | 0.01 | 0.00 | 0.15 | 0.04 | 0.07 | 0.04 | 0.01 | 0.01 | 0.04 | 0.01 | 0.00 | 0.16 | 0.04 | 0.05 | 0.00 | 0.00 |
| male | 3d AE | 1.00 | 0.75 | 0.00 | 0.00 | 0.00 | 0.01 | 0.00 | 0.00 | 0.05 | 0.00 | 0.01 | 0.00 | 0.00 | 0.00 | 0.00 | 0.00 | 0.00 | 0.01 | 0.00 | 0.00 | 0.00 | 0.00 |
| male | 3d AE | 1.00 | 1.18 | 0.00 | 0.00 | 0.00 | 0.01 | 0.00 | 0.00 | 0.06 | 0.00 | 0.01 | 0.01 | 0.00 | 0.00 | 0.00 | 0.00 | 0.00 | 0.03 | 0.00 | 0.00 | 0.00 | 0.00 |
| male | 3d AE | 1.00 | 0.61 | 0.00 | 0.00 | 0.00 | 0.01 | 0.01 | 0.00 | 0.03 | 0.00 | 0.01 | 0.00 | 0.00 | 0.00 | 0.00 | 0.00 | 0.00 | 0.01 | 0.00 | 0.00 | 0.00 | 0.00 |
| male | 3d AE | 1.00 | 1.04 | 0.00 | 0.00 | 0.00 | 0.01 | 0.00 | 0.00 | 0.06 | 0.00 | 0.01 | 0.01 | 0.00 | 0.00 | 0.00 | 0.00 | 0.00 | 0.04 | 0.00 | 0.00 | 0.00 | 0.00 |
| female | 3d AE | 2.00 | 2.35 | 1.87 | 0.00 | 0.05 | 0.10 | 0.04 | 0.02 | 0.21 | 0.08 | 0.08 | 0.07 | 0.02 | 0.01 | 0.06 | 0.01 | 0.00 | 0.22 | 0.07 | 0.11 | 0.00 | 0.00 |
| female | 3d AE | 2.00 | 1.77 | 1.41 | 0.00 | 0.04 | 0.05 | 0.01 | 0.01 | 0.16 | 0.05 | 0.05 | 0.04 | 0.01 | 0.00 | 0.05 | 0.00 | 0.00 | 0.15 | 0.04 | 0.05 | 0.00 | 0.00 |
| female | 3d AE | 2.00 | 1.82 | 1.70 | 0.00 | 0.02 | 0.06 | 0.02 | 0.01 | 0.15 | 0.04 | 0.04 | 0.05 | 0.01 | 0.00 | 0.04 | 0.01 | 0.00 | 0.19 | 0.05 | 0.05 | 0.00 | 0.00 |
| female | 3d AE | 2.00 | 2.60 | 2.34 | 0.00 | 0.04 | 0.12 | 0.02 | 0.03 | 0.24 | 0.10 | 0.12 | 0.09 | 0.01 | 0.01 | 0.06 | 0.02 | 0.00 | 0.35 | 0.09 | 0.11 | 0.00 | 0.00 |
| male | 3d AE | 2.00 | 0.91 | 0.00 | 0.00 | 0.00 | 0.01 | 0.01 | 0.00 | 0.04 | 0.00 | 0.01 | 0.00 | 0.00 | 0.00 | 0.00 | 0.00 | 0.00 | 0.02 | 0.00 | 0.00 | 0.00 | 0.00 |
| male | 3d AE | 2.00 | 2.07 | 0.00 | 0.00 | 0.01 | 0.04 | 0.01 | 0.00 | 0.17 | 0.00 | 0.03 | 0.03 | 0.01 | 0.00 | 0.00 | 0.01 | 0.00 | 0.11 | 0.03 | 0.00 | 0.00 | 0.00 |
| male | 3d AE | 2.00 | 1.54 | 0.00 | 0.00 | 0.01 | 0.02 | 0.01 | 0.00 | 0.10 | 0.00 | 0.01 | 0.02 | 0.00 | 0.01 | 0.00 | 0.00 | 0.00 | 0.05 | 0.00 | 0.00 | 0.00 | 0.00 |
| male | 3d AE | 2.00 | 1.01 | 0.00 | 0.00 | 0.00 | 0.02 | 0.01 | 0.00 | 0.06 | 0.00 | 0.02 | 0.01 | 0.00 | 0.00 | 0.00 | 0.00 | 0.00 | 0.04 | 0.00 | 0.00 | 0.00 | 0.00 |
| female | 7d AE | 1.00 | 1.46 | 2.03 | 0.00 | 0.01 | 0.01 | 0.02 | 0.00 | 0.05 | 0.03 | 0.00 | 0.00 | 0.00 | 0.00 | 0.00 | 0.00 | 0.00 | 0.00 | 0.00 | 0.00 | 0.00 | 0.00 |
| female | 7d AE | 1.00 | 1.46 | 1.43 | 0.00 | 0.01 | 0.01 | 0.01 | 0.00 | 0.05 | 0.02 | 0.00 | 0.00 | 0.00 | 0.00 | 0.00 | 0.00 | 0.00 | 0.00 | 0.00 | 0.00 | 0.00 | 0.00 |
| female | 7d AE | 1.00 | 1.52 | 1.56 | 0.00 | 0.00 | 0.01 | 0.01 | 0.00 | 0.05 | 0.02 | 0.00 | 0.00 | 0.00 | 0.00 | 0.00 | 0.00 | 0.00 | 0.00 | 0.00 | 0.00 | 0.00 | 0.00 |
| female | 7d AE | 1.00 | 1.85 | 1.51 | 0.00 | 0.01 | 0.02 | 0.02 | 0.00 | 0.09 | 0.04 | 0.00 | 0.00 | 0.00 | 0.01 | 0.00 | 0.00 | 0.00 | 0.00 | 0.00 | 0.00 | 0.00 | 0.00 |
| male | 7d AE | 1.00 | 1.25 | 0.00 | 0.00 | 0.00 | 0.00 | 0.01 | 0.00 | 0.04 | 0.00 | 0.00 | 0.00 | 0.00 | 0.01 | 0.00 | 0.00 | 0.00 | 0.00 | 0.00 | 0.00 | 0.00 | 0.00 |
| male | 7d AE | 1.00 | 1.49 | 0.00 | 0.00 | 0.00 | 0.00 | 0.01 | 0.00 | 0.04 | 0.00 | 0.00 | 0.00 | 0.00 | 0.00 | 0.00 | 0.00 | 0.00 | 0.00 | 0.00 | 0.00 | 0.00 | 0.00 |
| male | 7d AE | 1.00 | 1.26 | 0.00 | 0.00 | 0.00 | 0.00 | 0.00 | 0.00 | 0.04 | 0.00 | 0.00 | 0.00 | 0.00 | 0.00 | 0.00 | 0.00 | 0.00 | 0.00 | 0.00 | 0.00 | 0.00 | 0.00 |
| male | 7d AE | 1.00 | 1.48 | 0.00 | 0.00 | 0.00 | 0.00 | 0.00 | 0.00 | 0.04 | 0.00 | 0.00 | 0.00 | 0.00 | 0.00 | 0.00 | 0.00 | 0.00 | 0.00 | 0.00 | 0.00 | 0.00 | 0.00 |
| female | 7d AE | 2.00 | 1.91 | 1.88 | 0.00 | 0.02 | 0.02 | 0.02 | 0.00 | 0.07 | 0.03 | 0.00 | 0.00 | 0.00 | 0.00 | 0.00 | 0.00 | 0.00 | 0.00 | 0.00 | 0.00 | 0.00 | 0.00 |
| female | 7d AE | 2.00 | 1.82 | 2.30 | 0.00 | 0.02 | 0.02 | 0.01 | 0.00 | 0.05 | 0.02 | 0.00 | 0.00 | 0.00 | 0.00 | 0.00 | 0.00 | 0.00 | 0.00 | 0.00 | 0.00 | 0.00 | 0.00 |
| female | 7d AE | 2.00 | 1.58 | 1.81 | 0.00 | 0.01 | 0.01 | 0.01 | 0.00 | 0.06 | 0.02 | 0.00 | 0.00 | 0.00 | 0.00 | 0.00 | 0.00 | 0.00 | 0.00 | 0.00 | 0.00 | 0.00 | 0.00 |

| Sex | Time_point | Repeat | me-C28:0 | C29:2 | C29:1 (A) | C29:1 (B) | C29:1 (C) | C29:0 | C30:1 | me-C30:0 | C31:2 | C31:1 (A) | C31:1 (B) | C31:1 (C) | C31:0 | me-C32:0 | C32:2 | C32:1 | C33:2 | C33:1 | C35:2 | C35:1 | C37:2 |
|---|---|---|---|---|---|---|---|---|---|---|---|---|---|---|---|---|---|---|---|---|---|---|---|
| female | 7d AE | 2.00 | 1.69 | 1.62 | 0.00 | 0.01 | 0.01 | 0.02 | 0.00 | 0.07 | 0.02 | 0.00 | 0.00 | 0.00 | 0.00 | 0.00 | 0.00 | 0.00 | 0.00 | 0.00 | 0.00 | 0.00 | 0.00 |
| male | 7d AE | 2.00 | 1.78 | 0.00 | 0.00 | 0.00 | 0.00 | 0.01 | 0.00 | 0.06 | 0.00 | 0.00 | 0.00 | 0.00 | 0.00 | 0.00 | 0.00 | 0.00 | 0.00 | 0.00 | 0.00 | 0.00 | 0.00 |
| male | 7d AE | 2.00 | 1.23 | 0.00 | 0.00 | 0.00 | 0.00 | 0.01 | 0.00 | 0.03 | 0.00 | 0.00 | 0.00 | 0.00 | 0.00 | 0.00 | 0.00 | 0.00 | 0.00 | 0.00 | 0.00 | 0.00 | 0.00 |
| male | 7d AE | 2.00 | 1.46 | 0.00 | 0.00 | 0.00 | 0.00 | 0.00 | 0.00 | 0.05 | 0.00 | 0.00 | 0.00 | 0.00 | 0.00 | 0.00 | 0.00 | 0.00 | 0.00 | 0.00 | 0.00 | 0.00 | 0.00 |
| male | 7d AE | 2.00 | 1.42 | 0.00 | 0.00 | 0.00 | 0.00 | 0.01 | 0.00 | 0.04 | 0.00 | 0.00 | 0.00 | 0.00 | 0.00 | 0.00 | 0.00 | 0.00 | 0.00 | 0.00 | 0.00 | 0.00 | 0.00 |
| mixed | wL3 | 1.00 | 0.00 | 0.00 | 0.00 | 0.00 | 0.00 | 0.00 | 0.00 | 0.00 | 0.00 | 0.00 | 0.00 | 0.00 | 0.00 | 0.00 | 0.00 | 0.00 | 0.00 | 0.00 | 0.00 | 0.00 | 0.00 |
| mixed | wL3 | 1.00 | 0.00 | 0.00 | 0.00 | 0.00 | 0.00 | 0.00 | 0.00 | 0.00 | 0.00 | 0.00 | 0.00 | 0.00 | 0.00 | 0.00 | 0.00 | 0.00 | 0.00 | 0.00 | 0.00 | 0.00 | 0.00 |
| mixed | wL3 | 1.00 | 0.00 | 0.00 | 0.00 | 0.00 | 0.00 | 0.00 | 0.00 | 0.00 | 0.00 | 0.00 | 0.00 | 0.00 | 0.00 | 0.00 | 0.00 | 0.00 | 0.00 | 0.00 | 0.00 | 0.00 | 0.00 |
| mixed | wL3 | 1.00 | 0.00 | 0.00 | 0.00 | 0.00 | 0.00 | 0.00 | 0.00 | 0.00 | 0.00 | 0.00 | 0.00 | 0.00 | 0.00 | 0.00 | 0.00 | 0.00 | 0.00 | 0.00 | 0.00 | 0.00 | 0.00 |
| mixed | wL3 | 2.00 | 0.00 | 0.00 | 0.00 | 0.00 | 0.00 | 0.00 | 0.00 | 0.00 | 0.00 | 0.00 | 0.00 | 0.00 | 0.00 | 0.00 | 0.00 | 0.00 | 0.00 | 0.00 | 0.00 | 0.00 | 0.00 |
| mixed | wL3 | 2.00 | 0.01 | 0.00 | 0.00 | 0.00 | 0.00 | 0.00 | 0.00 | 0.00 | 0.00 | 0.00 | 0.00 | 0.00 | 0.00 | 0.00 | 0.00 | 0.00 | 0.00 | 0.00 | 0.00 | 0.00 | 0.00 |
| mixed | wL3 | 2.00 | 0.01 | 0.00 | 0.00 | 0.00 | 0.00 | 0.00 | 0.00 | 0.00 | 0.00 | 0.00 | 0.00 | 0.00 | 0.00 | 0.00 | 0.00 | 0.00 | 0.00 | 0.00 | 0.00 | 0.00 | 0.00 |
| mixed | wL3 | 2.00 | 0.00 | 0.00 | 0.00 | 0.00 | 0.00 | 0.00 | 0.00 | 0.00 | 0.00 | 0.00 | 0.00 | 0.00 | 0.00 | 0.00 | 0.00 | 0.00 | 0.00 | 0.00 | 0.00 | 0.00 | 0.00 |
| mixed | 48h_APF | 1.00 | 0.00 | 0.00 | 0.00 | 0.00 | 0.00 | 0.00 | 0.00 | 0.00 | 0.00 | 0.00 | 0.00 | 0.00 | 0.00 | 0.00 | 0.00 | 0.00 | 0.00 | 0.00 | 0.00 | 0.00 | 0.00 |
| mixed | 48h_APF | 1.00 | 0.00 | 0.00 | 0.00 | 0.00 | 0.00 | 0.00 | 0.00 | 0.00 | 0.00 | 0.00 | 0.00 | 0.00 | 0.00 | 0.00 | 0.00 | 0.00 | 0.00 | 0.00 | 0.00 | 0.00 | 0.00 |
| mixed | 48h_APF | 1.00 | 0.00 | 0.00 | 0.00 | 0.00 | 0.00 | 0.00 | 0.00 | 0.00 | 0.00 | 0.00 | 0.00 | 0.00 | 0.00 | 0.00 | 0.00 | 0.00 | 0.00 | 0.00 | 0.00 | 0.00 | 0.00 |
| mixed | 48h_APF | 1.00 | 0.00 | 0.00 | 0.00 | 0.00 | 0.00 | 0.00 | 0.00 | 0.00 | 0.00 | 0.00 | 0.00 | 0.00 | 0.00 | 0.00 | 0.00 | 0.00 | 0.00 | 0.00 | 0.00 | 0.00 | 0.00 |
| mixed | 48h_APF | 2.00 | 0.00 | 0.00 | 0.00 | 0.00 | 0.00 | 0.00 | 0.00 | 0.00 | 0.00 | 0.00 | 0.00 | 0.00 | 0.00 | 0.00 | 0.00 | 0.00 | 0.00 | 0.00 | 0.00 | 0.00 | 0.00 |
| mixed | 48h_APF | 2.00 | 0.00 | 0.00 | 0.00 | 0.00 | 0.00 | 0.00 | 0.00 | 0.00 | 0.00 | 0.00 | 0.00 | 0.00 | 0.00 | 0.00 | 0.00 | 0.00 | 0.00 | 0.00 | 0.00 | 0.00 | 0.00 |
| mixed | 48h_APF | 2.00 | 0.00 | 0.00 | 0.00 | 0.00 | 0.00 | 0.00 | 0.00 | 0.00 | 0.00 | 0.00 | 0.00 | 0.00 | 0.00 | 0.00 | 0.00 | 0.00 | 0.00 | 0.00 | 0.00 | 0.00 | 0.00 |
| mixed | 48h_APF | 2.00 | 0.00 | 0.00 | 0.00 | 0.00 | 0.00 | 0.00 | 0.00 | 0.00 | 0.00 | 0.00 | 0.00 | 0.00 | 0.00 | 0.00 | 0.00 | 0.00 | 0.00 | 0.00 | 0.00 | 0.00 | 0.00 |

| Sex | Time_point | Repeat | C21:0 | C22:1 | C22:0 | 9-C23:1 | 7-C23:1 | 5-C23:1 | C23:0 | C24:1 | C24:0 | me-C24:( | C25:2 | 9-C25:1 | 7-C25:1 | 5-C25:1 | C25:0 | C26:2 | C26:0 | me-C26:0 | C27:2 | C27:1 | C27:0 |
|---|---|---|---|---|---|---|---|---|---|---|---|---|---|---|---|---|---|---|---|---|---|---|---|
| female | 0h AE | 1 | 0.0000 | 0.0000 | 0.0000 | 0.0000 | 0.0000 | 0.0000 | 0.0000 | 0.0000 | 0.0000 | 0.0000 | 0.0000 | 0.0000 | 0.0000 | 0.0000 | 0.0000 | 0.0000 | 0.0000 | 0.0503 | 0.0000 | 0.0000 | 0.0090 |
| female | 0h AE | 1 | 0.0000 | 0.0000 | 0.0000 | 0.0000 | 0.0000 | 0.0000 | 0.0000 | 0.0000 | 0.0000 | 0.0000 | 0.0000 | 0.0000 | 0.0000 | 0.0000 | 0.0000 | 0.0000 | 0.0000 | 0.0474 | 0.0000 | 0.0000 | 0.0086 |
| female | 0h AE | 1 | 0.0000 | 0.0000 | 0.0000 | 0.0000 | 0.0000 | 0.0000 | 0.0000 | 0.0000 | 0.0000 | 0.0000 | 0.0000 | 0.0000 | 0.0000 | 0.0000 | 0.0000 | 0.0000 | 0.0000 | 0.0314 | 0.0000 | 0.0000 | 0.0000 |
| female | 0h AE | 1 | 0.0000 | 0.0000 | 0.0000 | 0.0000 | 0.0000 | 0.0000 | 0.0000 | 0.0000 | 0.0000 | 0.0000 | 0.0000 | 0.0000 | 0.0000 | 0.0000 | 0.0000 | 0.0000 | 0.0000 | 0.0140 | 0.0000 | 0.0000 | 0.0000 |
| male | 0h AE | 1 | 0.0000 | 0.0000 | 0.0000 | 0.0000 | 0.0000 | 0.0000 | 0.0000 | 0.0000 | 0.0000 | 0.0000 | 0.0000 | 0.0000 | 0.0000 | 0.0000 | 0.0000 | 0.0000 | 0.0000 | 0.0127 | 0.0000 | 0.0000 | 0.0000 |
| male | 0h AE | 1 | 0.0000 | 0.0000 | 0.0000 | 0.0000 | 0.0000 | 0.0000 | 0.0000 | 0.0000 | 0.0000 | 0.0000 | 0.0000 | 0.0000 | 0.0000 | 0.0000 | 0.0000 | 0.0000 | 0.0000 | 0.0218 | 0.0000 | 0.0000 | 0.0000 |
| male | 0h AE | 1 | 0.0000 | 0.0000 | 0.0000 | 0.0000 | 0.0000 | 0.0000 | 0.0000 | 0.0000 | 0.0000 | 0.0000 | 0.0000 | 0.0000 | 0.0000 | 0.0000 | 0.0000 | 0.0000 | 0.0000 | 0.0096 | 0.0000 | 0.0000 | 0.0000 |
| male | 0h AE | 1 | 0.0000 | 0.0000 | 0.0000 | 0.0000 | 0.0000 | 0.0000 | 0.0000 | 0.0000 | 0.0000 | 0.0000 | 0.0000 | 0.0000 | 0.0000 | 0.0000 | 0.0000 | 0.0000 | 0.0000 | 0.0431 | 0.0000 | 0.0000 | 0.0058 |
| female | 0h AE | 2 | 0.0000 | 0.0000 | 0.0000 | 0.0000 | 0.0000 | 0.0000 | 0.0000 | 0.0000 | 0.0000 | 0.0000 | 0.0000 | 0.0000 | 0.0000 | 0.0000 | 0.0000 | 0.0000 | 0.0000 | 0.0105 | 0.0000 | 0.0000 | 0.0000 |
| female | 0h AE | 2 | 0.0000 | 0.0000 | 0.0000 | 0.0000 | 0.0000 | 0.0000 | 0.0000 | 0.0000 | 0.0000 | 0.0000 | 0.0000 | 0.0000 | 0.0000 | 0.0000 | 0.0000 | 0.0000 | 0.0000 | 0.0378 | 0.0000 | 0.0000 | 0.0078 |
| female | 0h AE | 2 | 0.0000 | 0.0000 | 0.0000 | 0.0000 | 0.0000 | 0.0000 | 0.0000 | 0.0000 | 0.0000 | 0.0000 | 0.0000 | 0.0000 | 0.0000 | 0.0000 | 0.0000 | 0.0000 | 0.0000 | 0.0152 | 0.0000 | 0.0000 | 0.0045 |
| female | 0h AE | 2 | 0.0000 | 0.0000 | 0.0000 | 0.0000 | 0.0000 | 0.0000 | 0.0000 | 0.0000 | 0.0000 | 0.0000 | 0.0000 | 0.0000 | 0.0000 | 0.0000 | 0.0000 | 0.0000 | 0.0000 | 0.0098 | 0.0000 | 0.0000 | 0.0037 |
| male | 0h AE | 2 | 0.0000 | 0.0000 | 0.0000 | 0.0000 | 0.0000 | 0.0000 | 0.0000 | 0.0000 | 0.0000 | 0.0000 | 0.0000 | 0.0000 | 0.0000 | 0.0000 | 0.0000 | 0.0000 | 0.0000 | 0.0093 | 0.0000 | 0.0000 | 0.0025 |
| male | 0h AE | 2 | 0.0000 | 0.0000 | 0.0000 | 0.0000 | 0.0000 | 0.0000 | 0.0000 | 0.0000 | 0.0000 | 0.0000 | 0.0000 | 0.0000 | 0.0000 | 0.0000 | 0.0000 | 0.0000 | 0.0000 | 0.0174 | 0.0000 | 0.0000 | 0.0033 |
| male | 0h AE | 2 | 0.0000 | 0.0000 | 0.0000 | 0.0000 | 0.0000 | 0.0000 | 0.0000 | 0.0000 | 0.0000 | 0.0000 | 0.0000 | 0.0000 | 0.0000 | 0.0000 | 0.0000 | 0.0000 | 0.0000 | 0.0085 | 0.0000 | 0.0000 | 0.0044 |
| female | 6h AE | 1 | 0.0000 | 0.0000 | 0.0000 | 0.0000 | 0.0000 | 0.0000 | 0.0000 | 0.0000 | 0.0000 | 0.0000 | 0.0000 | 0.0000 | 0.0000 | 0.0000 | 0.0000 | 0.0000 | 0.0000 | 0.0205 | 0.0000 | 0.0000 | 0.0000 |
| female | 6h AE | 1 | 0.0000 | 0.0000 | 0.0000 | 0.0000 | 0.0000 | 0.0000 | 0.0000 | 0.0000 | 0.0000 | 0.0000 | 0.0000 | 0.0000 | 0.0000 | 0.0000 | 0.0000 | 0.0000 | 0.0000 | 0.0171 | 0.0000 | 0.0000 | 0.0000 |
| female | 6h AE | 1 | 0.0000 | 0.0000 | 0.0000 | 0.0000 | 0.0000 | 0.0000 | 0.0000 | 0.0000 | 0.0000 | 0.0000 | 0.0000 | 0.0000 | 0.0000 | 0.0000 | 0.0000 | 0.0000 | 0.0000 | 0.0170 | 0.0000 | 0.0000 | 0.0000 |
| female | 6h AE | 1 | 0.0000 | 0.0000 | 0.0000 | 0.0000 | 0.0000 | 0.0000 | 0.0000 | 0.0000 | 0.0000 | 0.0000 | 0.0000 | 0.0000 | 0.0000 | 0.0000 | 0.0000 | 0.0000 | 0.0000 | 0.0269 | 0.0000 | 0.0000 | 0.0000 |
| male | 6h AE | 1 | 0.0000 | 0.0000 | 0.0000 | 0.0000 | 0.0000 | 0.0000 | 0.0000 | 0.0000 | 0.0000 | 0.0000 | 0.0000 | 0.0000 | 0.0000 | 0.0000 | 0.0000 | 0.0000 | 0.0000 | 0.0184 | 0.0000 | 0.0000 | 0.0000 |
| male | 6h AE | 1 | 0.0000 | 0.0000 | 0.0000 | 0.0000 | 0.0000 | 0.0000 | 0.0000 | 0.0000 | 0.0000 | 0.0000 | 0.0000 | 0.0000 | 0.0000 | 0.0000 | 0.0000 | 0.0000 | 0.0000 | 0.0760 | 0.0000 | 0.0164 | 0.0122 |
| male | 6h AE | 1 | 0.0000 | 0.0000 | 0.0000 | 0.0000 | 0.0000 | 0.0000 | 0.0000 | 0.0000 | 0.0000 | 0.0000 | 0.0000 | 0.0000 | 0.0000 | 0.0000 | 0.0000 | 0.0000 | 0.0000 | 0.0152 | 0.0000 | 0.0032 | 0.0030 |
| male | 6h AE | 1 | 0.0000 | 0.0000 | 0.0000 | 0.0000 | 0.0000 | 0.0000 | 0.0000 | 0.0000 | 0.0000 | 0.0000 | 0.0000 | 0.0000 | 0.0000 | 0.0000 | 0.0000 | 0.0000 | 0.0000 | 0.0130 | 0.0000 | 0.0000 | 0.0042 |
| female | 6h AE | 2 | 0.0000 | 0.0000 | 0.0000 | 0.0000 | 0.0000 | 0.0000 | 0.0000 | 0.0000 | 0.0000 | 0.0000 | 0.0000 | 0.0000 | 0.0000 | 0.0000 | 0.0000 | 0.0000 | 0.0000 | 0.0146 | 0.0000 | 0.0039 | 0.0037 |
| female | 6h AE | 2 | 0.0000 | 0.0000 | 0.0000 | 0.0000 | 0.0000 | 0.0000 | 0.0000 | 0.0000 | 0.0000 | 0.0000 | 0.0000 | 0.0000 | 0.0000 | 0.0000 | 0.0000 | 0.0000 | 0.0000 | 0.0148 | 0.0000 | 0.0000 | 0.0000 |
| female | 6h AE | 2 | 0.0000 | 0.0000 | 0.0000 | 0.0000 | 0.0000 | 0.0000 | 0.0000 | 0.0000 | 0.0000 | 0.0000 | 0.0000 | 0.0000 | 0.0000 | 0.0000 | 0.0000 | 0.0000 | 0.0000 | 0.0353 | 0.0000 | 0.0000 | 0.0055 |
| female | 6h AE | 2 | 0.0000 | 0.0000 | 0.0000 | 0.0000 | 0.0000 | 0.0000 | 0.0000 | 0.0000 | 0.0000 | 0.0000 | 0.0000 | 0.0000 | 0.0000 | 0.0000 | 0.0000 | 0.0000 | 0.0000 | 0.0128 | 0.0000 | 0.0000 | 0.0000 |
| male | 6h AE | 2 | 0.0000 | 0.0000 | 0.0000 | 0.0000 | 0.0000 | 0.0000 | 0.0000 | 0.0000 | 0.0000 | 0.0000 | 0.0000 | 0.0000 | 0.0000 | 0.0000 | 0.0000 | 0.0000 | 0.0000 | 0.0474 | 0.0000 | 0.0134 | 0.0110 |
| male | 6h AE | 2 | 0.0000 | 0.0000 | 0.0000 | 0.0000 | 0.0000 | 0.0000 | 0.0000 | 0.0000 | 0.0000 | 0.0000 | 0.0000 | 0.0000 | 0.0000 | 0.0000 | 0.0000 | 0.0000 | 0.0000 | 0.0158 | 0.0000 | 0.0000 | 0.0000 |
| male | 6h AE | 2 | 0.0000 | 0.0000 | 0.0000 | 0.0000 | 0.0000 | 0.0000 | 0.0000 | 0.0000 | 0.0000 | 0.0000 | 0.0000 | 0.0000 | 0.0000 | 0.0000 | 0.0000 | 0.0000 | 0.0000 | 0.0116 | 0.0000 | 0.0000 | 0.0000 |
| male | 6h AE | 2 | 0.0000 | 0.0000 | 0.0000 | 0.0000 | 0.0000 | 0.0000 | 0.0000 | 0.0000 | 0.0000 | 0.0000 | 0.0000 | 0.0000 | 0.0000 | 0.0000 | 0.0000 | 0.0000 | 0.0000 | 0.0179 | 0.0000 | 0.0000 | 0.0000 |
| female | 24h AE | 1 | 0.0051 | 0.0000 | 0.0032 | 0.0402 | 0.3657 | 0.0275 | 0.0538 | 0.0173 | 0.0021 | 0.1098 | 0.0210 | 0.2266 | 0.3410 | 0.0101 | 0.0392 | 0.0000 | 0.0000 | 0.2377 | 0.1330 | 0.1228 | 0.0272 |
| female | 24h AE | 1 | 0.0070 | 0.0000 | 0.0032 | 0.0755 | 0.6356 | 0.0645 | 0.1122 | 0.0300 | 0.0034 | 0.1899 | 0.0379 | 0.3326 | 0.4509 | 0.0240 | 0.0629 | 0.0000 | 0.0000 | 0.2812 | 0.1738 | 0.1663 | 0.0422 |
| female | 24h AE | 1 | 0.0111 | 0.0000 | 0.0070 | 0.0723 | 0.6886 | 0.0721 | 0.1116 | 0.0410 | 0.0024 | 0.2648 | 0.0420 | 0.4010 | 0.5378 | 0.0246 | 0.0660 | 0.0000 | 0.0000 | 0.3153 | 0.1447 | 0.1607 | 0.0472 |
| female | 24h AE | 1 | 0.0059 | 0.0000 | 0.0018 | 0.0241 | 0.3621 | 0.0308 | 0.0633 | 0.0105 | 0.0030 | 0.0939 | 0.0187 | 0.1989 | 0.2985 | 0.0054 | 0.0350 | 0.0000 | 0.0000 | 0.1694 | 0.0809 | 0.0598 | 0.0209 |
| male | 24h AE | 1 | 0.0099 | 0.0077 | 0.0120 | 0.1313 | 1.2825 | 0.0649 | 0.1012 | 0.0231 | 0.0020 | 0.1554 | 0.0000 | 0.1456 | 0.2419 | 0.0000 | 0.0255 | 0.0000 | 0.0000 | 0.1826 | 0.0000 | 0.0153 | 0.0175 |
| male | 24h AE | 1 | 0.0254 | 0.0219 | 0.0281 | 0.2903 | 1.5363 | 0.1639 | 0.1564 | 0.0572 | 0.0051 | 0.4415 | 0.0000 | 0.3239 | 0.4461 | 0.0000 | 0.0457 | 0.0000 | 0.0000 | 0.2923 | 0.0000 | 0.0574 | 0.0282 |
| male | 24h AE | 1 | 0.0083 | 0.0087 | 0.0182 | 0.1297 | 1.4850 | 0.0782 | 0.0928 | 0.0252 | 0.0018 | 0.2145 | 0.0000 | 0.1478 | 0.2743 | 0.0000 | 0.0242 | 0.0000 | 0.0000 | 0.1782 | 0.0000 | 0.0229 | 0.0132 |
| male | 24h AE | 1 | 0.0069 | 0.0079 | 0.0178 | 0.1405 | 1.4709 | 0.0516 | 0.0815 | 0.0303 | 0.0030 | 0.1853 | 0.0000 | 0.1660 | 0.2868 | 0.0000 | 0.0208 | 0.0000 | 0.0000 | 0.1841 | 0.0000 | 0.0244 | 0.0126 |
| female | 24h AE | 2 | 0.0076 | 0.0000 | 0.0000 | 0.0380 | 0.2511 | 0.0239 | 0.0771 | 0.0115 | 0.0000 | 0.0801 | 0.0092 | 0.2270 | 0.3256 | 0.0108 | 0.0540 | 0.0000 | 0.0000 | 0.1924 | 0.1050 | 0.1865 | 0.0443 |

| Sex | Time_point | Repeat | C21:0 | C22:1 | C22:0 | 9-C23:1 | 7-C23:1 | 5-C23:1 | C23:0 | C24:1 | C24:0 | me-C24:( | C25:2 | 9-C25:1 | 7-C25:1 | 5-C25:1 | C25:0 | C26:2 | C26:0 | me-C26:0 | C27:2 | C27:1 | C27:0 |
|---|---|---|---|---|---|---|---|---|---|---|---|---|---|---|---|---|---|---|---|---|---|---|---|
| female | 24h AE | 2 | 0.0051 | 0.0000 | 0.0025 | 0.0372 | 0.3018 | 0.0252 | 0.0698 | 0.0146 | 0.0000 | 0.0852 | 0.0135 | 0.1767 | 0.2848 | 0.0079 | 0.0323 | 0.0000 | 0.0000 | 0.1827 | 0.0802 | 0.1342 | 0.0230 |
| female | 24h AE | 2 | 0.0074 | 0.0000 | 0.0000 | 0.0423 | 0.2963 | 0.0210 | 0.0706 | 0.0202 | 0.0000 | 0.1024 | 0.0232 | 0.2287 | 0.2633 | 0.0148 | 0.0360 | 0.0000 | 0.0000 | 0.1981 | 0.1189 | 0.1022 | 0.0272 |
| female | 24h AE | 2 | 0.0111 | 0.0000 | 0.0049 | 0.1032 | 0.6363 | 0.0647 | 0.1546 | 0.0401 | 0.0000 | 0.2057 | 0.0364 | 0.4846 | 0.5905 | 0.0332 | 0.0858 | 0.0000 | 0.0000 | 0.3428 | 0.1874 | 0.2817 | 0.0656 |
| male | 24h AE | 2 | 0.0216 | 0.0203 | 0.0189 | 0.2803 | 1.7378 | 0.1270 | 0.1551 | 0.0551 | 0.0052 | 0.4077 | 0.0000 | 0.2650 | 0.4194 | 0.0000 | 0.0435 | 0.0000 | 0.0000 | 0.2556 | 0.0000 | 0.0244 | 0.0243 |
| male | 24h AE | 2 | 0.0213 | 0.0240 | 0.0246 | 0.2733 | 1.5694 | 0.1280 | 0.1513 | 0.0537 | 0.0053 | 0.3169 | 0.0000 | 0.2444 | 0.3301 | 0.0000 | 0.0383 | 0.0000 | 0.0000 | 0.2449 | 0.0000 | 0.0324 | 0.0288 |
| male | 24h AE | 2 | 0.0128 | 0.0083 | 0.0101 | 0.1219 | 1.4057 | 0.0413 | 0.0842 | 0.0226 | 0.0000 | 0.1704 | 0.0000 | 0.1304 | 0.2277 | 0.0000 | 0.0243 | 0.0000 | 0.0000 | 0.1805 | 0.0000 | 0.0106 | 0.0154 |
| male | 24h AE | 2 | 0.0092 | 0.0096 | 0.0148 | 0.1395 | 1.5936 | 0.0836 | 0.0974 | 0.0318 | 0.0000 | 0.2030 | 0.0000 | 0.1544 | 0.2737 | 0.0000 | 0.0250 | 0.0000 | 0.0000 | 0.1934 | 0.0000 | 0.0242 | 0.0155 |
| female | 3d AE | 1 | 0.0068 | 0.0000 | 0.0054 | 0.0408 | 0.4398 | 0.0241 | 0.2271 | 0.0203 | 0.0106 | 0.3588 | 0.1270 | 0.5478 | 0.6066 | 0.0501 | 0.2279 | 0.0288 | 0.0072 | 0.8178 | 3.0447 | 0.5795 | 0.1054 |
| female | 3d AE | 1 | 0.0240 | 0.0000 | 0.0120 | 0.0988 | 0.5122 | 0.0459 | 0.3705 | 0.0443 | 0.0254 | 0.7030 | 0.2400 | 0.7299 | 0.6007 | 0.0733 | 0.2887 | 0.0456 | 0.0146 | 0.6801 | 2.0014 | 0.6400 | 0.1486 |
| female | 3d AE | 1 | 0.0087 | 0.0000 | 0.0058 | 0.0329 | 0.1958 | 0.0127 | 0.2122 | 0.0184 | 0.0084 | 0.3604 | 0.1112 | 0.4441 | 0.3904 | 0.0410 | 0.2175 | 0.0209 | 0.0072 | 0.7511 | 2.4275 | 0.4785 | 0.1465 |
| female | 3d AE | 1 | 0.0079 | 0.0000 | 0.0090 | 0.0482 | 0.2715 | 0.0199 | 0.2277 | 0.0182 | 0.0085 | 0.3843 | 0.1230 | 0.5298 | 0.4331 | 0.0401 | 0.2384 | 0.0289 | 0.0071 | 0.8631 | 3.1009 | 0.5466 | 0.1019 |
| male | 3d AE | 1 | 0.0235 | 0.0159 | 0.0450 | 0.3081 | 2.6121 | 0.2181 | 0.3666 | 0.0849 | 0.0086 | 0.6375 | 0.0000 | 0.3441 | 1.5645 | 0.0000 | 0.0669 | 0.0000 | 0.0045 | 0.3618 | 0.0000 | 0.0237 | 0.0231 |
| male | 3d AE | 1 | 0.0212 | 0.0276 | 0.0529 | 0.3563 | 4.0566 | 0.2665 | 0.4555 | 0.1154 | 0.0092 | 0.6294 | 0.0000 | 0.4145 | 2.5884 | 0.0000 | 0.0706 | 0.0000 | 0.0037 | 0.5495 | 0.0000 | 0.0379 | 0.0232 |
| male | 3d AE | 1 | 0.0126 | 0.0124 | 0.0335 | 0.2646 | 3.0064 | 0.1772 | 0.3346 | 0.0560 | 0.0045 | 0.4499 | 0.0000 | 0.2248 | 1.3056 | 0.0000 | 0.0460 | 0.0000 | 0.0016 | 0.3061 | 0.0000 | 0.0240 | 0.0108 |
| male | 3d AE | 1 | 0.0139 | 0.0176 | 0.0440 | 0.3052 | 3.4042 | 0.2136 | 0.4308 | 0.0919 | 0.0070 | 0.5996 | 0.0000 | 0.4308 | 2.2661 | 0.0000 | 0.0783 | 0.0000 | 0.0017 | 0.4208 | 0.0000 | 0.0398 | 0.0270 |
| female | 3d AE | 2 | 0.0288 | 0.0000 | 0.0178 | 0.1296 | 0.5677 | 0.0486 | 0.4453 | 0.0422 | 0.0233 | 0.6312 | 0.2737 | 0.7833 | 0.6892 | 0.0722 | 0.3075 | 0.0449 | 0.0143 | 0.6846 | 2.2212 | 0.8167 | 0.2053 |
| female | 3d AE | 2 | 0.0095 | 0.0000 | 0.0077 | 0.0499 | 0.3245 | 0.0254 | 0.3353 | 0.0146 | 0.0100 | 0.3591 | 0.1357 | 0.4793 | 0.4301 | 0.0364 | 0.1706 | 0.0266 | 0.0060 | 0.7142 | 2.6861 | 0.4803 | 0.0974 |
| female | 3d AE | 2 | 0.0111 | 0.0000 | 0.0089 | 0.0437 | 0.2592 | 0.0214 | 0.2627 | 0.0153 | 0.0117 | 0.2875 | 0.1205 | 0.4943 | 0.5079 | 0.0466 | 0.2177 | 0.0201 | 0.0060 | 0.6693 | 2.5781 | 0.6402 | 0.1085 |
| female | 3d AE | 2 | 0.0171 | 0.0000 | 0.0087 | 0.0772 | 0.4387 | 0.0360 | 0.2793 | 0.0254 | 0.0160 | 0.4244 | 0.1782 | 0.6776 | 0.7287 | 0.0680 | 0.2406 | 0.0361 | 0.0136 | 0.9289 | 3.7698 | 0.8519 | 0.1491 |
| male | 3d AE | 2 | 0.0166 | 0.0187 | 0.0675 | 0.3550 | 3.6945 | 0.2387 | 0.4379 | 0.0939 | 0.0105 | 0.6269 | 0.0000 | 0.3457 | 1.6570 | 0.0000 | 0.0703 | 0.0000 | 0.0016 | 0.3704 | 0.0000 | 0.0173 | 0.0185 |
| male | 3d AE | 2 | 0.0725 | 0.1020 | 0.0705 | 0.8390 | 2.7680 | 0.5506 | 0.6173 | 0.3581 | 0.0284 | 1.6877 | 0.0000 | 0.8397 | 2.1773 | 0.0000 | 0.1858 | 0.0000 | 0.0077 | 0.6505 | 0.0000 | 0.0838 | 0.0530 |
| male | 3d AE | 2 | 0.0413 | 0.0470 | 0.0667 | 0.5640 | 2.4796 | 0.3609 | 0.4942 | 0.2065 | 0.0170 | 1.0708 | 0.0000 | 0.5166 | 1.7344 | 0.0000 | 0.1165 | 0.0000 | 0.0057 | 0.5980 | 0.0000 | 0.0518 | 0.0461 |
| male | 3d AE | 2 | 0.0209 | 0.0240 | 0.0578 | 0.3652 | 3.9293 | 0.2740 | 0.4801 | 0.1365 | 0.0089 | 0.7183 | 0.0000 | 0.4457 | 2.2876 | 0.0000 | 0.0809 | 0.0000 | 0.0029 | 0.4237 | 0.0000 | 0.0416 | 0.0218 |
| female | 7d AE | 1 | 0.0117 | 0.0000 | 0.0077 | 0.0316 | 0.1342 | 0.0144 | 0.3385 | 0.0119 | 0.0128 | 0.2905 | 0.1471 | 0.2835 | 0.2168 | 0.0378 | 0.2557 | 0.0254 | 0.0070 | 0.9127 | 3.1392 | 0.3443 | 0.1387 |
| female | 7d AE | 1 | 0.0140 | 0.0000 | 0.0077 | 0.0315 | 0.1299 | 0.0156 | 0.3658 | 0.0087 | 0.0104 | 0.3170 | 0.1793 | 0.2806 | 0.2242 | 0.0384 | 0.2293 | 0.0267 | 0.0062 | 0.7269 | 2.5005 | 0.2999 | 0.1579 |
| female | 7d AE | 1 | 0.0117 | 0.0000 | 0.0096 | 0.0376 | 0.1501 | 0.0151 | 0.4343 | 0.0105 | 0.0104 | 0.3199 | 0.1776 | 0.3034 | 0.2580 | 0.0354 | 0.2868 | 0.0205 | 0.0077 | 0.8000 | 2.5587 | 0.3265 | 0.1474 |
| female | 7d AE | 1 | 0.0245 | 0.0000 | 0.0153 | 0.0580 | 0.2368 | 0.0268 | 0.4682 | 0.0227 | 0.0234 | 0.5195 | 0.2660 | 0.4366 | 0.3657 | 0.0765 | 0.3609 | 0.0394 | 0.0115 | 0.6982 | 2.0295 | 0.4163 | 0.2215 |
| male | 7d AE | 1 | 0.0419 | 0.0409 | 0.0384 | 0.2832 | 5.2016 | 0.4158 | 0.5398 | 0.1182 | 0.0109 | 0.7866 | 0.0000 | 0.2031 | 1.5830 | 0.0000 | 0.0828 | 0.0000 | 0.0016 | 0.5177 | 0.0000 | 0.0203 | 0.0272 |
| male | 7d AE | 1 | 0.0297 | 0.0222 | 0.0842 | 0.2722 | 4.4062 | 0.3830 | 0.6292 | 0.0997 | 0.0096 | 0.8274 | 0.0000 | 0.2154 | 1.6481 | 0.0000 | 0.0902 | 0.0000 | 0.0028 | 0.4849 | 0.0000 | 0.0162 | 0.0317 |
| male | 7d AE | 1 | 0.0316 | 0.0405 | 0.0630 | 0.2577 | 4.5205 | 0.4041 | 0.5727 | 0.0884 | 0.0077 | 0.6840 | 0.0000 | 0.1901 | 1.6936 | 0.0000 | 0.0680 | 0.0000 | 0.0031 | 0.4763 | 0.0000 | 0.0211 | 0.0263 |
| male | 7d AE | 1 | 0.0465 | 0.0348 | 0.0796 | 0.2869 | 4.6122 | 0.4454 | 0.6717 | 0.1113 | 0.0116 | 0.7772 | 0.0000 | 0.2621 | 1.5116 | 0.0000 | 0.0915 | 0.0000 | 0.0000 | 0.4901 | 0.0000 | 0.0176 | 0.0350 |
| female | 7d AE | 2 | 0.0152 | 0.0000 | 0.0114 | 0.0489 | 0.2051 | 0.0257 | 0.4479 | 0.0139 | 0.0155 | 0.3646 | 0.2344 | 0.3728 | 0.3512 | 0.0669 | 0.2916 | 0.0353 | 0.0092 | 1.0879 | 3.8050 | 0.4062 | 0.1598 |
| female | 7d AE | 2 | 0.0181 | 0.0000 | 0.0113 | 0.0444 | 0.2246 | 0.0218 | 0.4812 | 0.0149 | 0.0139 | 0.4843 | 0.2579 | 0.5083 | 0.4297 | 0.0839 | 0.4090 | 0.0350 | 0.0142 | 1.2206 | 4.2982 | 0.5521 | 0.2086 |
| female | 7d AE | 2 | 0.0120 | 0.0000 | 0.0067 | 0.0296 | 0.1415 | 0.0123 | 0.3491 | 0.0102 | 0.0111 | 0.2942 | 0.1804 | 0.3460 | 0.2563 | 0.0415 | 0.2476 | 0.0193 | 0.0107 | 0.7885 | 3.1476 | 0.3960 | 0.1300 |
| female | 7d AE | 2 | 0.0139 | 0.0000 | 0.0140 | 0.0536 | 0.2963 | 0.0300 | 0.4099 | 0.0192 | 0.0125 | 0.3829 | 0.2263 | 0.3137 | 0.4130 | 0.0607 | 0.2575 | 0.0346 | 0.0123 | 0.9621 | 3.4588 | 0.3554 | 0.1323 |
| male | 7d AE | 2 | 0.0723 | 0.0813 | 0.0744 | 0.4115 | 2.8877 | 0.5294 | 0.5431 | 0.1783 | 0.0197 | 1.1451 | 0.0000 | 0.1866 | 1.0360 | 0.0000 | 0.0971 | 0.0000 | 0.0105 | 0.5082 | 0.0000 | 0.0181 | 0.0367 |
| male | 7d AE | 2 | 0.0318 | 0.0361 | 0.0424 | 0.2616 | 3.8805 | 0.3301 | 0.5260 | 0.0976 | 0.0118 | 0.6232 | 0.0000 | 0.1793 | 1.4485 | 0.0000 | 0.0740 | 0.0000 | 0.0047 | 0.4429 | 0.0000 | 0.0157 | 0.0326 |
| male | 7d AE | 2 | 0.0386 | 0.0353 | 0.0529 | 0.3368 | 5.6830 | 0.3697 | 0.5528 | 0.0995 | 0.0085 | 0.6971 | 0.0000 | 0.1558 | 1.5687 | 0.0000 | 0.0661 | 0.0000 | 0.0033 | 0.5594 | 0.0000 | 0.0173 | 0.0303 |
| male | 7d AE | 2 | 0.0299 | 0.0325 | 0.0656 | 0.2800 | 5.0685 | 0.3419 | 0.5490 | 0.0948 | 0.0076 | 0.5451 | 0.0000 | 0.1309 | 1.3644 | 0.0000 | 0.0791 | 0.0000 | 0.0020 | 0.4673 | 0.0000 | 0.0146 | 0.0285 |
| mixed | wL3 | 1 | 0.0000 | 0.0000 | 0.0000 | 0.0000 | 0.0000 | 0.0000 | 0.0000 | 0.0000 | 0.0000 | 0.0000 | 0.0000 | 0.0000 | 0.0000 | 0.0000 | 0.0000 | 0.0000 | 0.0000 | 0.0000 | 0.0000 | 0.0000 | 0.0000 |

| Sex | Time_point | Repeat | C21:0 | C22:1 | C22:0 | 9-C23:1 | 7-C23:1 | 5-C23:1 | C23:0 | C24:1 | C24:0 | me-C24:0 | C25:2 | 9-C25:1 | 7-C25:1 | 5-C25:1 | C25:0 | C26:2 | C26:0 | me-C26:0 | C27:2 | C27:1 | C27:0 |
|---|---|---|---|---|---|---|---|---|---|---|---|---|---|---|---|---|---|---|---|---|---|---|---|
| mixed | wL3 | 1 | 0.0000 | 0.0000 | 0.0000 | 0.0000 | 0.0000 | 0.0000 | 0.0000 | 0.0000 | 0.0000 | 0.0000 | 0.0000 | 0.0000 | 0.0000 | 0.0000 | 0.0000 | 0.0000 | 0.0000 | 0.0000 | 0.0000 | 0.0000 | 0.0000 |
| mixed | wL3 | 1 | 0.0000 | 0.0000 | 0.0000 | 0.0000 | 0.0000 | 0.0000 | 0.0000 | 0.0000 | 0.0000 | 0.0000 | 0.0000 | 0.0000 | 0.0000 | 0.0000 | 0.0000 | 0.0000 | 0.0000 | 0.0000 | 0.0000 | 0.0000 | 0.0000 |
| mixed | wL3 | 1 | 0.0000 | 0.0000 | 0.0000 | 0.0000 | 0.0000 | 0.0000 | 0.0000 | 0.0000 | 0.0000 | 0.0000 | 0.0000 | 0.0000 | 0.0000 | 0.0000 | 0.0000 | 0.0000 | 0.0000 | 0.0000 | 0.0000 | 0.0000 | 0.0000 |
| mixed | wL3 | 2 | 0.0000 | 0.0000 | 0.0000 | 0.0000 | 0.0000 | 0.0000 | 0.0000 | 0.0000 | 0.0000 | 0.0000 | 0.0000 | 0.0000 | 0.0000 | 0.0000 | 0.0000 | 0.0000 | 0.0000 | 0.0000 | 0.0000 | 0.0000 | 0.0000 |
| mixed | wL3 | 2 | 0.0000 | 0.0000 | 0.0000 | 0.0000 | 0.0000 | 0.0000 | 0.0000 | 0.0000 | 0.0000 | 0.0000 | 0.0000 | 0.0000 | 0.0000 | 0.0000 | 0.0000 | 0.0000 | 0.0000 | 0.0000 | 0.0000 | 0.0000 | 0.0000 |
| mixed | wL3 | 2 | 0.0000 | 0.0000 | 0.0000 | 0.0000 | 0.0000 | 0.0000 | 0.0000 | 0.0000 | 0.0000 | 0.0000 | 0.0000 | 0.0000 | 0.0000 | 0.0000 | 0.0000 | 0.0000 | 0.0000 | 0.0000 | 0.0000 | 0.0000 | 0.0000 |
| mixed | wL3 | 2 | 0.0000 | 0.0000 | 0.0000 | 0.0000 | 0.0000 | 0.0000 | 0.0000 | 0.0000 | 0.0000 | 0.0000 | 0.0000 | 0.0000 | 0.0000 | 0.0000 | 0.0000 | 0.0000 | 0.0000 | 0.0000 | 0.0000 | 0.0000 | 0.0000 |
| mixed | 48h_APF | 1 | 0.0000 | 0.0000 | 0.0000 | 0.0000 | 0.0000 | 0.0000 | 0.0000 | 0.0000 | 0.0000 | 0.0000 | 0.0000 | 0.0000 | 0.0000 | 0.0000 | 0.0000 | 0.0000 | 0.0000 | 0.0000 | 0.0000 | 0.0000 | 0.0000 |
| mixed | 48h_APF | 1 | 0.0000 | 0.0000 | 0.0000 | 0.0000 | 0.0000 | 0.0000 | 0.0000 | 0.0000 | 0.0000 | 0.0000 | 0.0000 | 0.0000 | 0.0000 | 0.0000 | 0.0000 | 0.0000 | 0.0000 | 0.0000 | 0.0000 | 0.0000 | 0.0000 |
| mixed | 48h_APF | 1 | 0.0000 | 0.0000 | 0.0000 | 0.0000 | 0.0000 | 0.0000 | 0.0000 | 0.0000 | 0.0000 | 0.0000 | 0.0000 | 0.0000 | 0.0000 | 0.0000 | 0.0000 | 0.0000 | 0.0000 | 0.0000 | 0.0000 | 0.0000 | 0.0000 |
| mixed | 48h_APF | 1 | 0.0000 | 0.0000 | 0.0000 | 0.0000 | 0.0000 | 0.0000 | 0.0000 | 0.0000 | 0.0000 | 0.0000 | 0.0000 | 0.0000 | 0.0000 | 0.0000 | 0.0000 | 0.0000 | 0.0000 | 0.0000 | 0.0000 | 0.0000 | 0.0000 |
| mixed | 48h_APF | 2 | 0.0000 | 0.0000 | 0.0000 | 0.0000 | 0.0000 | 0.0000 | 0.0000 | 0.0000 | 0.0000 | 0.0000 | 0.0000 | 0.0000 | 0.0000 | 0.0000 | 0.0000 | 0.0000 | 0.0000 | 0.0000 | 0.0000 | 0.0000 | 0.0000 |
| mixed | 48h_APF | 2 | 0.0000 | 0.0000 | 0.0000 | 0.0000 | 0.0000 | 0.0000 | 0.0000 | 0.0000 | 0.0000 | 0.0000 | 0.0000 | 0.0000 | 0.0000 | 0.0000 | 0.0000 | 0.0000 | 0.0000 | 0.0000 | 0.0000 | 0.0000 | 0.0000 |
| mixed | 48h_APF | 2 | 0.0000 | 0.0000 | 0.0000 | 0.0000 | 0.0000 | 0.0000 | 0.0000 | 0.0000 | 0.0000 | 0.0000 | 0.0000 | 0.0000 | 0.0000 | 0.0000 | 0.0000 | 0.0000 | 0.0000 | 0.0000 | 0.0000 | 0.0000 | 0.0000 |
| mixed | 48h_APF | 2 | 0.0000 | 0.0000 | 0.0000 | 0.0000 | 0.0000 | 0.0000 | 0.0000 | 0.0000 | 0.0000 | 0.0000 | 0.0000 | 0.0000 | 0.0000 | 0.0000 | 0.0000 | 0.0000 | 0.0000 | 0.0000 | 0.0000 | 0.0000 | 0.0000 |

| Sex | Time_point | Repeat | me-C28:0 | C29:2 | C29:1 (A) | C29:1 (B) | C29:1 (C) | C29:0 | C30:1 | me-C30:0 | C31:2 | C31:1 (A) | C31:1 (B) | C31:1 (C) | C31:0 | me-C32:0 | C32:2 | C32:1 | C33:2 | C33:1 | C35:2 | C35:1 | C37:2 |
|---|---|---|---|---|---|---|---|---|---|---|---|---|---|---|---|---|---|---|---|---|---|---|---|
| female | 0h AE | 1 | 2.2995 | 0.0000 | 0.0546 | 0.0722 | 0.3119 | 0.0315 | 0.1946 | 0.4702 | 0.2033 | 0.4069 | 0.5756 | 0.1009 | 0.0332 | 0.1518 | 0.1028 | 0.0555 | 1.9386 | 0.6432 | 0.9652 | 0.1238 | 0.1430 |
| female | 0h AE | 1 | 2.2607 | 0.0000 | 0.0665 | 0.0486 | 0.2530 | 0.0307 | 0.1326 | 0.4926 | 0.1827 | 0.4386 | 0.5263 | 0.0899 | 0.0241 | 0.1608 | 0.0954 | 0.0491 | 1.9857 | 0.6171 | 0.8471 | 0.1314 | 0.1137 |
| female | 0h AE | 1 | 1.7688 | 0.0000 | 0.0250 | 0.0271 | 0.1483 | 0.0128 | 0.0794 | 0.3501 | 0.1137 | 0.2356 | 0.3292 | 0.0501 | 0.0135 | 0.0817 | 0.0381 | 0.0214 | 1.0501 | 0.3204 | 0.3347 | 0.0620 | 0.0483 |
| female | 0h AE | 1 | 1.6240 | 0.0000 | 0.0212 | 0.0183 | 0.1173 | 0.0130 | 0.0530 | 0.3218 | 0.1146 | 0.2355 | 0.2790 | 0.0505 | 0.0102 | 0.0904 | 0.0290 | 0.0204 | 0.9592 | 0.2922 | 0.3897 | 0.0603 | 0.0386 |
| male | 0h AE | 1 | 1.3146 | 0.0000 | 0.0170 | 0.0182 | 0.0986 | 0.0083 | 0.0471 | 0.2252 | 0.0981 | 0.1897 | 0.1991 | 0.0278 | 0.0073 | 0.0896 | 0.0257 | 0.0168 | 0.8382 | 0.2244 | 0.2397 | 0.0316 | 0.0300 |
| male | 0h AE | 1 | 1.5886 | 0.0000 | 0.0240 | 0.0251 | 0.1293 | 0.0169 | 0.0562 | 0.2574 | 0.1031 | 0.2158 | 0.2556 | 0.0643 | 0.0136 | 0.0701 | 0.0312 | 0.0203 | 0.9121 | 0.2428 | 0.3075 | 0.0427 | 0.0361 |
| male | 0h AE | 1 | 1.2798 | 0.0000 | 0.0101 | 0.0174 | 0.0823 | 0.0110 | 0.0382 | 0.2080 | 0.0850 | 0.1488 | 0.2066 | 0.0221 | 0.0055 | 0.0602 | 0.0144 | 0.0133 | 0.6463 | 0.1954 | 0.2387 | 0.0325 | 0.0180 |
| male | 0h AE | 1 | 1.7922 | 0.0000 | 0.0385 | 0.0606 | 0.2254 | 0.0195 | 0.1052 | 0.3287 | 0.1934 | 0.3523 | 0.3992 | 0.0565 | 0.0181 | 0.0983 | 0.0722 | 0.0319 | 1.4342 | 0.4361 | 0.5995 | 0.0683 | 0.0952 |
| female | 0h AE | 2 | 0.9855 | 0.0000 | 0.0113 | 0.0115 | 0.0819 | 0.0136 | 0.0407 | 0.2346 | 0.0643 | 0.1609 | 0.2560 | 0.0432 | 0.0115 | 0.0719 | 0.0227 | 0.0149 | 0.7217 | 0.2344 | 0.3255 | 0.0481 | 0.0425 |
| female | 0h AE | 2 | 1.8233 | 0.0000 | 0.0578 | 0.0638 | 0.2920 | 0.0353 | 0.1541 | 0.3915 | 0.1777 | 0.4192 | 0.5772 | 0.0843 | 0.0242 | 0.1310 | 0.1109 | 0.0549 | 1.9166 | 0.5134 | 0.7660 | 0.0879 | 0.1694 |
| female | 0h AE | 2 | 1.4564 | 0.0000 | 0.0223 | 0.0193 | 0.1368 | 0.0238 | 0.0925 | 0.3566 | 0.0904 | 0.2887 | 0.3951 | 0.0814 | 0.0199 | 0.1375 | 0.0540 | 0.0277 | 1.4217 | 0.4393 | 0.5133 | 0.0817 | 0.0798 |
| female | 0h AE | 2 | 1.9254 | 0.0000 | 0.0234 | 0.0251 | 0.1482 | 0.0114 | 0.0689 | 0.4524 | 0.1265 | 0.3224 | 0.4549 | 0.0420 | 0.0112 | 0.1419 | 0.0508 | 0.0271 | 1.4528 | 0.4518 | 0.5155 | 0.0605 | 0.0674 |
| male | 0h AE | 2 | 1.1948 | 0.0000 | 0.0121 | 0.0104 | 0.0783 | 0.0082 | 0.0371 | 0.2231 | 0.0734 | 0.1432 | 0.2055 | 0.0304 | 0.0083 | 0.0588 | 0.0284 | 0.0142 | 0.7119 | 0.2141 | 0.2603 | 0.0323 | 0.0336 |
| male | 0h AE | 2 | 1.2742 | 0.0000 | 0.0131 | 0.0195 | 0.1144 | 0.0138 | 0.0526 | 0.2651 | 0.0673 | 0.1810 | 0.2821 | 0.0525 | 0.0133 | 0.0753 | 0.0306 | 0.0164 | 0.8222 | 0.2815 | 0.3490 | 0.0575 | 0.0462 |
| male | 0h AE | 2 | 0.9799 | 0.0000 | 0.0141 | 0.0133 | 0.0862 | 0.0080 | 0.0307 | 0.1848 | 0.0561 | 0.1444 | 0.1835 | 0.0282 | 0.0083 | 0.0488 | 0.0175 | 0.0123 | 0.6365 | 0.1724 | 0.1949 | 0.0271 | 0.0274 |
| female | 6h AE | 1 | 2.3697 | 0.0000 | 0.0666 | 0.0654 | 0.2318 | 0.0281 | 0.1074 | 0.4563 | 0.2693 | 0.4162 | 0.4134 | 0.0413 | 0.0195 | 0.0989 | 0.0801 | 0.0333 | 1.5444 | 0.3693 | 0.5487 | 0.0709 | 0.0722 |
| female | 6h AE | 1 | 1.6949 | 0.0000 | 0.0340 | 0.0482 | 0.1708 | 0.0152 | 0.0717 | 0.3793 | 0.1639 | 0.2964 | 0.2982 | 0.0404 | 0.0121 | 0.0979 | 0.0456 | 0.0196 | 1.0391 | 0.2650 | 0.3621 | 0.0481 | 0.0519 |
| female | 6h AE | 1 | 1.7439 | 0.0000 | 0.0402 | 0.0477 | 0.1786 | 0.0126 | 0.0660 | 0.3286 | 0.1493 | 0.3564 | 0.2745 | 0.0439 | 0.0142 | 0.0891 | 0.0615 | 0.0269 | 1.0074 | 0.3078 | 0.3635 | 0.0425 | 0.0596 |
| female | 6h AE | 1 | 1.7821 | 0.0000 | 0.0520 | 0.0562 | 0.1655 | 0.0151 | 0.0676 | 0.3510 | 0.1590 | 0.3466 | 0.2925 | 0.0369 | 0.0113 | 0.0992 | 0.0599 | 0.0276 | 1.1012 | 0.3193 | 0.3638 | 0.0516 | 0.0504 |
| male | 6h AE | 1 | 1.5508 | 0.0000 | 0.0523 | 0.0349 | 0.1498 | 0.0133 | 0.0526 | 0.2567 | 0.2004 | 0.2469 | 0.2220 | 0.0408 | 0.0071 | 0.0606 | 0.0343 | 0.0129 | 0.8507 | 0.2817 | 0.3095 | 0.0440 | 0.0319 |
| male | 6h AE | 1 | 2.3011 | 0.0000 | 0.1473 | 0.1408 | 0.4563 | 0.0340 | 0.2123 | 0.4453 | 0.4327 | 0.5883 | 0.5033 | 0.0725 | 0.0215 | 0.1132 | 0.1575 | 0.0539 | 2.3426 | 0.6565 | 0.9916 | 0.1088 | 0.1650 |
| male | 6h AE | 1 | 1.6129 | 0.0000 | 0.0384 | 0.0421 | 0.1808 | 0.0106 | 0.0476 | 0.2571 | 0.1756 | 0.2529 | 0.2409 | 0.0293 | 0.0066 | 0.0680 | 0.0371 | 0.0161 | 0.9969 | 0.2564 | 0.3208 | 0.0380 | 0.0512 |
| male | 6h AE | 1 | 1.4106 | 0.0000 | 0.0317 | 0.0341 | 0.1469 | 0.0097 | 0.0618 | 0.2676 | 0.1508 | 0.2461 | 0.2726 | 0.0341 | 0.0076 | 0.0617 | 0.0501 | 0.0140 | 0.7764 | 0.1949 | 0.2958 | 0.0336 | 0.0355 |
| female | 6h AE | 2 | 1.8322 | 0.0000 | 0.0451 | 0.0434 | 0.2400 | 0.0178 | 0.0701 | 0.4382 | 0.1776 | 0.4228 | 0.4150 | 0.0598 | 0.0139 | 0.1277 | 0.0618 | 0.0285 | 1.5059 | 0.4244 | 0.4917 | 0.0588 | 0.0683 |
| female | 6h AE | 2 | 1.9884 | 0.0000 | 0.0393 | 0.0435 | 0.1840 | 0.0101 | 0.0627 | 0.3816 | 0.1644 | 0.2744 | 0.2865 | 0.0488 | 0.0128 | 0.0981 | 0.0424 | 0.0210 | 1.0565 | 0.3048 | 0.5006 | 0.0612 | 0.0513 |
| female | 6h AE | 2 | 2.0199 | 0.0000 | 0.0725 | 0.0751 | 0.3081 | 0.0211 | 0.1229 | 0.3988 | 0.1827 | 0.3828 | 0.4390 | 0.0681 | 0.0133 | 0.1576 | 0.0801 | 0.0274 | 1.6713 | 0.4551 | 0.7082 | 0.0819 | 0.1014 |
| female | 6h AE | 2 | 1.6379 | 0.0000 | 0.0586 | 0.0482 | 0.1809 | 0.0142 | 0.0850 | 0.3834 | 0.1554 | 0.2856 | 0.3278 | 0.0344 | 0.0151 | 0.1044 | 0.0443 | 0.0227 | 1.0750 | 0.3359 | 0.4438 | 0.0683 | 0.0700 |
| male | 6h AE | 2 | 1.7962 | 0.0000 | 0.1099 | 0.1035 | 0.3490 | 0.0256 | 0.1491 | 0.3371 | 0.2897 | 0.4119 | 0.4295 | 0.0647 | 0.0171 | 0.1012 | 0.0938 | 0.0409 | 1.4838 | 0.4007 | 0.6516 | 0.0899 | 0.1257 |
| male | 6h AE | 2 | 1.3873 | 0.0000 | 0.0482 | 0.0461 | 0.1717 | 0.0096 | 0.0570 | 0.2325 | 0.1504 | 0.2226 | 0.2220 | 0.0360 | 0.0087 | 0.0617 | 0.0469 | 0.0194 | 0.8599 | 0.2317 | 0.3169 | 0.0418 | 0.0410 |
| male | 6h AE | 2 | 1.3271 | 0.0000 | 0.0396 | 0.0394 | 0.1472 | 0.0092 | 0.0472 | 0.2360 | 0.1449 | 0.1985 | 0.2167 | 0.0323 | 0.0084 | 0.0628 | 0.0265 | 0.0110 | 0.7504 | 0.2279 | 0.2750 | 0.0304 | 0.0332 |
| male | 6h AE | 2 | 1.4556 | 0.0000 | 0.0274 | 0.0370 | 0.1439 | 0.0121 | 0.0548 | 0.2102 | 0.1393 | 0.1861 | 0.1961 | 0.0258 | 0.0057 | 0.0625 | 0.0278 | 0.0092 | 0.6854 | 0.1996 | 0.2338 | 0.0450 | 0.0305 |
| female | 24h AE | 1 | 2.6738 | 0.0000 | 0.1293 | 0.0490 | 0.1659 | 0.0200 | 0.0483 | 0.4419 | 0.2107 | 0.2586 | 0.2041 | 0.0412 | 0.0069 | 0.0994 | 0.0528 | 0.0174 | 0.9250 | 0.2113 | 0.3361 | 0.0288 | 0.0421 |
| female | 24h AE | 1 | 2.6460 | 0.0000 | 0.1816 | 0.0699 | 0.2366 | 0.0318 | 0.0683 | 0.4852 | 0.2985 | 0.3989 | 0.3210 | 0.0683 | 0.0182 | 0.1192 | 0.0890 | 0.0311 | 1.0606 | 0.3001 | 0.3805 | 0.0390 | 0.0616 |
| female | 24h AE | 1 | 2.3249 | 0.0000 | 0.1885 | 0.1210 | 0.2950 | 0.0380 | 0.1357 | 0.4072 | 0.2578 | 0.3864 | 0.3326 | 0.0605 | 0.0208 | 0.1301 | 0.0942 | 0.0307 | 1.2568 | 0.3531 | 0.4922 | 0.0669 | 0.1083 |
| female | 24h AE | 1 | 2.1288 | 0.0000 | 0.0919 | 0.0653 | 0.1601 | 0.0203 | 0.0525 | 0.3158 | 0.1506 | 0.2358 | 0.1916 | 0.0273 | 0.0145 | 0.0766 | 0.0331 | 0.0119 | 0.6381 | 0.1818 | 0.2587 | 0.0364 | 0.0408 |
| male | 24h AE | 1 | 1.7666 | 0.0000 | 0.0503 | 0.0282 | 0.1302 | 0.0119 | 0.0326 | 0.2120 | 0.1164 | 0.1360 | 0.1343 | 0.0232 | 0.0068 | 0.0548 | 0.0186 | 0.0087 | 0.4321 | 0.1361 | 0.1287 | 0.0244 | 0.0176 |
| male | 24h AE | 1 | 2.4112 | 0.0000 | 0.1020 | 0.0885 | 0.2770 | 0.0244 | 0.0811 | 0.3351 | 0.2300 | 0.2910 | 0.2504 | 0.0475 | 0.0186 | 0.0884 | 0.0552 | 0.0179 | 0.9575 | 0.2503 | 0.3089 | 0.0323 | 0.0559 |
| male | 24h AE | 1 | 1.7701 | 0.0000 | 0.0406 | 0.0392 | 0.1223 | 0.0045 | 0.0279 | 0.1736 | 0.1049 | 0.1122 | 0.1180 | 0.0126 | 0.0061 | 0.0519 | 0.0217 | 0.0089 | 0.4436 | 0.1086 | 0.1566 | 0.0228 | 0.0210 |
| male | 24h AE | 1 | 1.8251 | 0.0000 | 0.0425 | 0.0356 | 0.1241 | 0.0108 | 0.0400 | 0.2266 | 0.1179 | 0.1341 | 0.1433 | 0.0280 | 0.0071 | 0.0573 | 0.0247 | 0.0095 | 0.4583 | 0.1185 | 0.1731 | 0.0205 | 0.0305 |
| female | 24h AE | 2 | 2.2230 | 0.0000 | 0.1261 | 0.0557 | 0.1761 | 0.0225 | 0.0528 | 0.3676 | 0.2167 | 0.2565 | 0.2478 | 0.0357 | 0.0138 | 0.0929 | 0.0281 | 0.0134 | 0.6935 | 0.2123 | 0.2915 | 0.0335 | 0.0329 |

| Sex | Time_point | Repeat | me-C28:0 | C29:2 | C29:1 (A) | C29:1 (B) | C29:1 (C) | C29:0 | C30:1 | me-C30:0 | C31:2 | C31:1 (A) | C31:1 (B) | C31:1 (C) | C31:0 | me-C32:0 | C32:2 | C32:1 | C33:2 | C33:1 | C35:2 | C35:1 | C37:2 |
|---|---|---|---|---|---|---|---|---|---|---|---|---|---|---|---|---|---|---|---|---|---|---|---|
| female | 24h AE | 2 | 2.3470 | 0.0000 | 0.1094 | 0.0557 | 0.1780 | 0.0173 | 0.0539 | 0.4043 | 0.2192 | 0.2794 | 0.2442 | 0.0240 | 0.0117 | 0.0941 | 0.0593 | 0.0210 | 0.8865 | 0.2351 | 0.2903 | 0.0275 | 0.0398 |
| female | 24h AE | 2 | 2.3195 | 0.0000 | 0.0899 | 0.0508 | 0.1611 | 0.0172 | 0.0661 | 0.3080 | 0.1641 | 0.2016 | 0.2071 | 0.0324 | 0.0053 | 0.0837 | 0.0480 | 0.0196 | 0.7064 | 0.1927 | 0.2730 | 0.0389 | 0.0406 |
| female | 24h AE | 2 | 2.5607 | 0.0000 | 0.1955 | 0.1052 | 0.2871 | 0.0432 | 0.1243 | 0.4997 | 0.2757 | 0.4589 | 0.4845 | 0.0521 | 0.0239 | 0.1568 | 0.0844 | 0.0336 | 1.2597 | 0.3746 | 0.5010 | 0.0567 | 0.0893 |
| male | 24h AE | 2 | 2.0885 | 0.0000 | 0.0803 | 0.0465 | 0.1693 | 0.0149 | 0.0653 | 0.2882 | 0.1549 | 0.1912 | 0.1864 | 0.0345 | 0.0106 | 0.0880 | 0.0394 | 0.0202 | 0.6625 | 0.1675 | 0.2970 | 0.0389 | 0.0392 |
| male | 24h AE | 2 | 2.2961 | 0.0000 | 0.0746 | 0.0551 | 0.1889 | 0.0180 | 0.0496 | 0.2701 | 0.1644 | 0.1887 | 0.1977 | 0.0269 | 0.0103 | 0.0806 | 0.0296 | 0.0162 | 0.6244 | 0.1612 | 0.2488 | 0.0487 | 0.0377 |
| male | 24h AE | 2 | 1.8326 | 0.0000 | 0.0456 | 0.0283 | 0.1059 | 0.0208 | 0.0355 | 0.2345 | 0.0881 | 0.1095 | 0.1346 | 0.0233 | 0.0052 | 0.0608 | 0.0169 | 0.0000 | 0.3976 | 0.1121 | 0.1709 | 0.0269 | 0.0146 |
| male | 24h AE | 2 | 2.2800 | 0.0000 | 0.0491 | 0.0351 | 0.1328 | 0.0110 | 0.0483 | 0.2403 | 0.1225 | 0.1350 | 0.1286 | 0.0250 | 0.0079 | 0.0550 | 0.0265 | 0.0000 | 0.4701 | 0.1212 | 0.1896 | 0.0226 | 0.0209 |
| female | 3d AE | 1 | 2.2228 | 1.5659 | 0.0000 | 0.0312 | 0.0750 | 0.0115 | 0.0131 | 0.1759 | 0.0766 | 0.0642 | 0.0571 | 0.0116 | 0.0065 | 0.0389 | 0.0087 | 0.0000 | 0.2178 | 0.0618 | 0.0753 | 0.0000 | 0.0000 |
| female | 3d AE | 1 | 2.2712 | 1.7658 | 0.0000 | 0.0362 | 0.0877 | 0.0218 | 0.0211 | 0.1816 | 0.0634 | 0.0615 | 0.0606 | 0.0194 | 0.0085 | 0.0558 | 0.0076 | 0.0000 | 0.2110 | 0.0523 | 0.0596 | 0.0000 | 0.0000 |
| female | 3d AE | 1 | 1.8363 | 1.5490 | 0.0000 | 0.0259 | 0.0631 | 0.0131 | 0.0110 | 0.1590 | 0.0523 | 0.0504 | 0.0451 | 0.0092 | 0.0067 | 0.0434 | 0.0081 | 0.0000 | 0.2011 | 0.0607 | 0.0589 | 0.0000 | 0.0000 |
| female | 3d AE | 1 | 1.8680 | 1.6534 | 0.0000 | 0.0303 | 0.0583 | 0.0141 | 0.0000 | 0.1521 | 0.0443 | 0.0698 | 0.0418 | 0.0110 | 0.0073 | 0.0415 | 0.0096 | 0.0000 | 0.1640 | 0.0391 | 0.0453 | 0.0000 | 0.0000 |
| male | 3d AE | 1 | 0.7469 | 0.0000 | 0.0000 | 0.0023 | 0.0104 | 0.0042 | 0.0000 | 0.0453 | 0.0000 | 0.0059 | 0.0000 | 0.0000 | 0.0024 | 0.0000 | 0.0000 | 0.0000 | 0.0127 | 0.0000 | 0.0000 | 0.0000 | 0.0000 |
| male | 3d AE | 1 | 1.1766 | 0.0000 | 0.0000 | 0.0036 | 0.0101 | 0.0041 | 0.0000 | 0.0559 | 0.0000 | 0.0078 | 0.0116 | 0.0000 | 0.0038 | 0.0000 | 0.0000 | 0.0000 | 0.0300 | 0.0000 | 0.0000 | 0.0000 | 0.0000 |
| male | 3d AE | 1 | 0.6070 | 0.0000 | 0.0000 | 0.0025 | 0.0077 | 0.0058 | 0.0000 | 0.0332 | 0.0000 | 0.0060 | 0.0040 | 0.0013 | 0.0018 | 0.0000 | 0.0000 | 0.0000 | 0.0136 | 0.0000 | 0.0000 | 0.0000 | 0.0000 |
| male | 3d AE | 1 | 1.0391 | 0.0000 | 0.0000 | 0.0019 | 0.0070 | 0.0036 | 0.0000 | 0.0647 | 0.0000 | 0.0091 | 0.0131 | 0.0025 | 0.0020 | 0.0000 | 0.0000 | 0.0000 | 0.0417 | 0.0000 | 0.0000 | 0.0000 | 0.0000 |
| female | 3d AE | 2 | 2.3521 | 1.8704 | 0.0000 | 0.0543 | 0.0981 | 0.0368 | 0.0166 | 0.2083 | 0.0810 | 0.0750 | 0.0654 | 0.0186 | 0.0089 | 0.0571 | 0.0134 | 0.0000 | 0.2241 | 0.0682 | 0.1074 | 0.0000 | 0.0000 |
| female | 3d AE | 2 | 1.7727 | 1.4131 | 0.0000 | 0.0376 | 0.0498 | 0.0135 | 0.0137 | 0.1600 | 0.0450 | 0.0509 | 0.0440 | 0.0063 | 0.0050 | 0.0484 | 0.0039 | 0.0000 | 0.1541 | 0.0393 | 0.0517 | 0.0000 | 0.0000 |
| female | 3d AE | 2 | 1.8201 | 1.7037 | 0.0000 | 0.0248 | 0.0553 | 0.0192 | 0.0129 | 0.1490 | 0.0440 | 0.0421 | 0.0548 | 0.0078 | 0.0021 | 0.0390 | 0.0055 | 0.0000 | 0.1894 | 0.0465 | 0.0509 | 0.0000 | 0.0000 |
| female | 3d AE | 2 | 2.6017 | 2.3394 | 0.0000 | 0.0408 | 0.1179 | 0.0234 | 0.0291 | 0.2366 | 0.1031 | 0.1169 | 0.0949 | 0.0121 | 0.0095 | 0.0560 | 0.0156 | 0.0000 | 0.3516 | 0.0947 | 0.1121 | 0.0000 | 0.0000 |
| male | 3d AE | 2 | 0.9117 | 0.0000 | 0.0000 | 0.0000 | 0.0096 | 0.0052 | 0.0000 | 0.0425 | 0.0000 | 0.0073 | 0.0000 | 0.0000 | 0.0015 | 0.0000 | 0.0000 | 0.0000 | 0.0193 | 0.0000 | 0.0000 | 0.0000 | 0.0000 |
| male | 3d AE | 2 | 2.0706 | 0.0000 | 0.0000 | 0.0126 | 0.0374 | 0.0143 | 0.0000 | 0.1682 | 0.0000 | 0.0284 | 0.0335 | 0.0088 | 0.0048 | 0.0000 | 0.0077 | 0.0000 | 0.1067 | 0.0280 | 0.0000 | 0.0000 | 0.0000 |
| male | 3d AE | 2 | 1.5396 | 0.0000 | 0.0000 | 0.0068 | 0.0174 | 0.0093 | 0.0000 | 0.1008 | 0.0000 | 0.0140 | 0.0174 | 0.0020 | 0.0055 | 0.0000 | 0.0000 | 0.0000 | 0.0461 | 0.0000 | 0.0000 | 0.0000 | 0.0000 |
| male | 3d AE | 2 | 1.0071 | 0.0000 | 0.0000 | 0.0036 | 0.0249 | 0.0098 | 0.0000 | 0.0553 | 0.0000 | 0.0152 | 0.0112 | 0.0011 | 0.0025 | 0.0000 | 0.0000 | 0.0000 | 0.0365 | 0.0000 | 0.0000 | 0.0000 | 0.0000 |
| female | 7d AE | 1 | 1.4605 | 2.0256 | 0.0000 | 0.0079 | 0.0141 | 0.0151 | 0.0000 | 0.0543 | 0.0254 | 0.0000 | 0.0017 | 0.0000 | 0.0026 | 0.0000 | 0.0000 | 0.0000 | 0.0000 | 0.0000 | 0.0000 | 0.0000 | 0.0000 |
| female | 7d AE | 1 | 1.4640 | 1.4319 | 0.0000 | 0.0062 | 0.0123 | 0.0136 | 0.0000 | 0.0544 | 0.0180 | 0.0000 | 0.0000 | 0.0000 | 0.0000 | 0.0000 | 0.0000 | 0.0000 | 0.0000 | 0.0000 | 0.0000 | 0.0000 | 0.0000 |
| female | 7d AE | 1 | 1.5190 | 1.5628 | 0.0000 | 0.0000 | 0.0113 | 0.0127 | 0.0000 | 0.0511 | 0.0195 | 0.0000 | 0.0000 | 0.0000 | 0.0032 | 0.0000 | 0.0000 | 0.0000 | 0.0000 | 0.0000 | 0.0000 | 0.0000 | 0.0000 |
| female | 7d AE | 1 | 1.8529 | 1.5086 | 0.0000 | 0.0085 | 0.0183 | 0.0176 | 0.0000 | 0.0945 | 0.0356 | 0.0000 | 0.0000 | 0.0000 | 0.0054 | 0.0000 | 0.0000 | 0.0000 | 0.0000 | 0.0000 | 0.0000 | 0.0000 | 0.0000 |
| male | 7d AE | 1 | 1.2456 | 0.0000 | 0.0000 | 0.0000 | 0.0000 | 0.0052 | 0.0000 | 0.0410 | 0.0000 | 0.0000 | 0.0000 | 0.0000 | 0.0054 | 0.0000 | 0.0000 | 0.0000 | 0.0000 | 0.0000 | 0.0000 | 0.0000 | 0.0000 |
| male | 7d AE | 1 | 1.4944 | 0.0000 | 0.0000 | 0.0000 | 0.0000 | 0.0056 | 0.0000 | 0.0391 | 0.0000 | 0.0000 | 0.0000 | 0.0000 | 0.0000 | 0.0000 | 0.0000 | 0.0000 | 0.0000 | 0.0000 | 0.0000 | 0.0000 | 0.0000 |
| male | 7d AE | 1 | 1.2579 | 0.0000 | 0.0000 | 0.0000 | 0.0000 | 0.0035 | 0.0000 | 0.0394 | 0.0000 | 0.0000 | 0.0000 | 0.0000 | 0.0000 | 0.0000 | 0.0000 | 0.0000 | 0.0000 | 0.0000 | 0.0000 | 0.0000 | 0.0000 |
| male | 7d AE | 1 | 1.4790 | 0.0000 | 0.0000 | 0.0000 | 0.0000 | 0.0041 | 0.0000 | 0.0422 | 0.0000 | 0.0000 | 0.0000 | 0.0000 | 0.0000 | 0.0000 | 0.0000 | 0.0000 | 0.0000 | 0.0000 | 0.0000 | 0.0000 | 0.0000 |
| female | 7d AE | 2 | 1.9133 | 1.8829 | 0.0000 | 0.0164 | 0.0201 | 0.0174 | 0.0000 | 0.0677 | 0.0250 | 0.0000 | 0.0000 | 0.0000 | 0.0000 | 0.0000 | 0.0000 | 0.0000 | 0.0000 | 0.0000 | 0.0000 | 0.0000 | 0.0000 |
| female | 7d AE | 2 | 1.8206 | 2.2985 | 0.0000 | 0.0172 | 0.0207 | 0.0138 | 0.0000 | 0.0510 | 0.0235 | 0.0000 | 0.0000 | 0.0000 | 0.0000 | 0.0000 | 0.0000 | 0.0000 | 0.0000 | 0.0000 | 0.0000 | 0.0000 | 0.0000 |
| female | 7d AE | 2 | 1.5783 | 1.8098 | 0.0000 | 0.0108 | 0.0140 | 0.0113 | 0.0000 | 0.0597 | 0.0249 | 0.0000 | 0.0000 | 0.0000 | 0.0000 | 0.0000 | 0.0000 | 0.0000 | 0.0000 | 0.0000 | 0.0000 | 0.0000 | 0.0000 |
| female | 7d AE | 2 | 1.6893 | 1.6153 | 0.0000 | 0.0075 | 0.0146 | 0.0175 | 0.0000 | 0.0669 | 0.0170 | 0.0000 | 0.0000 | 0.0000 | 0.0000 | 0.0000 | 0.0000 | 0.0000 | 0.0000 | 0.0000 | 0.0000 | 0.0000 | 0.0000 |
| male | 7d AE | 2 | 1.7843 | 0.0000 | 0.0000 | 0.0000 | 0.0000 | 0.0058 | 0.0000 | 0.0572 | 0.0000 | 0.0000 | 0.0000 | 0.0000 | 0.0000 | 0.0000 | 0.0000 | 0.0000 | 0.0000 | 0.0000 | 0.0000 | 0.0000 | 0.0000 |
| male | 7d AE | 2 | 1.2291 | 0.0000 | 0.0000 | 0.0000 | 0.0000 | 0.0056 | 0.0000 | 0.0310 | 0.0000 | 0.0000 | 0.0000 | 0.0000 | 0.0000 | 0.0000 | 0.0000 | 0.0000 | 0.0000 | 0.0000 | 0.0000 | 0.0000 | 0.0000 |
| male | 7d AE | 2 | 1.4621 | 0.0000 | 0.0000 | 0.0000 | 0.0000 | 0.0043 | 0.0000 | 0.0477 | 0.0000 | 0.0000 | 0.0000 | 0.0000 | 0.0000 | 0.0000 | 0.0000 | 0.0000 | 0.0000 | 0.0000 | 0.0000 | 0.0000 | 0.0000 |
| male | 7d AE | 2 | 1.4207 | 0.0000 | 0.0000 | 0.0000 | 0.0000 | 0.0073 | 0.0000 | 0.0374 | 0.0000 | 0.0000 | 0.0000 | 0.0000 | 0.0000 | 0.0000 | 0.0000 | 0.0000 | 0.0000 | 0.0000 | 0.0000 | 0.0000 | 0.0000 |
| mixed | wL3 | 1 | 0.0000 | 0.0000 | 0.0000 | 0.0000 | 0.0000 | 0.0000 | 0.0000 | 0.0000 | 0.0000 | 0.0000 | 0.0000 | 0.0000 | 0.0000 | 0.0000 | 0.0000 | 0.0000 | 0.0000 | 0.0000 | 0.0000 | 0.0000 | 0.0000 |

| Sex | Time_point | Repeat | me-C28:0 | C29:2 | C29:1 (A) | C29:1 (B) | C29:1 (C) | C29:0 | C30:1 | me-C30:0 | C31:2 | C31:1 (A) | C31:1 (B) | C31:1 (C) | C31:0 | me-C32:0 | C32:2 | C32:1 | C33:2 | C33:1 | C35:2 | C35:1 | C37:2 |
|---|---|---|---|---|---|---|---|---|---|---|---|---|---|---|---|---|---|---|---|---|---|---|---|
| mixed | wL3 | 1 | 0.0000 | 0.0000 | 0.0000 | 0.0000 | 0.0000 | 0.0000 | 0.0000 | 0.0000 | 0.0000 | 0.0000 | 0.0000 | 0.0000 | 0.0000 | 0.0000 | 0.0000 | 0.0000 | 0.0000 | 0.0000 | 0.0000 | 0.0000 | 0.0000 |
| mixed | wL3 | 1 | 0.0000 | 0.0000 | 0.0000 | 0.0000 | 0.0000 | 0.0000 | 0.0000 | 0.0000 | 0.0000 | 0.0000 | 0.0000 | 0.0000 | 0.0000 | 0.0000 | 0.0000 | 0.0000 | 0.0000 | 0.0000 | 0.0000 | 0.0000 | 0.0000 |
| mixed | wL3 | 1 | 0.0000 | 0.0000 | 0.0000 | 0.0000 | 0.0000 | 0.0000 | 0.0000 | 0.0000 | 0.0000 | 0.0000 | 0.0000 | 0.0000 | 0.0000 | 0.0000 | 0.0000 | 0.0000 | 0.0000 | 0.0000 | 0.0000 | 0.0000 | 0.0000 |
| mixed | wL3 | 2 | 0.0000 | 0.0000 | 0.0000 | 0.0000 | 0.0000 | 0.0000 | 0.0000 | 0.0000 | 0.0000 | 0.0000 | 0.0000 | 0.0000 | 0.0000 | 0.0000 | 0.0000 | 0.0000 | 0.0000 | 0.0000 | 0.0000 | 0.0000 | 0.0000 |
| mixed | wL3 | 2 | 0.0086 | 0.0000 | 0.0000 | 0.0000 | 0.0000 | 0.0000 | 0.0000 | 0.0000 | 0.0000 | 0.0000 | 0.0000 | 0.0000 | 0.0000 | 0.0000 | 0.0000 | 0.0000 | 0.0000 | 0.0000 | 0.0000 | 0.0000 | 0.0000 |
| mixed | wL3 | 2 | 0.0058 | 0.0000 | 0.0000 | 0.0000 | 0.0000 | 0.0000 | 0.0000 | 0.0000 | 0.0000 | 0.0000 | 0.0000 | 0.0000 | 0.0000 | 0.0000 | 0.0000 | 0.0000 | 0.0000 | 0.0000 | 0.0000 | 0.0000 | 0.0000 |
| mixed | wL3 | 2 | 0.0000 | 0.0000 | 0.0000 | 0.0000 | 0.0000 | 0.0000 | 0.0000 | 0.0000 | 0.0000 | 0.0000 | 0.0000 | 0.0000 | 0.0000 | 0.0000 | 0.0000 | 0.0000 | 0.0000 | 0.0000 | 0.0000 | 0.0000 | 0.0000 |
| mixed | 48h_APF | 1 | 0.0000 | 0.0000 | 0.0000 | 0.0000 | 0.0000 | 0.0000 | 0.0000 | 0.0000 | 0.0000 | 0.0000 | 0.0000 | 0.0000 | 0.0000 | 0.0000 | 0.0000 | 0.0000 | 0.0000 | 0.0000 | 0.0000 | 0.0000 | 0.0000 |
| mixed | 48h_APF | 1 | 0.0000 | 0.0000 | 0.0000 | 0.0000 | 0.0000 | 0.0000 | 0.0000 | 0.0000 | 0.0000 | 0.0000 | 0.0000 | 0.0000 | 0.0000 | 0.0000 | 0.0000 | 0.0000 | 0.0000 | 0.0000 | 0.0000 | 0.0000 | 0.0000 |
| mixed | 48h_APF | 1 | 0.0000 | 0.0000 | 0.0000 | 0.0000 | 0.0000 | 0.0000 | 0.0000 | 0.0000 | 0.0000 | 0.0000 | 0.0000 | 0.0000 | 0.0000 | 0.0000 | 0.0000 | 0.0000 | 0.0000 | 0.0000 | 0.0000 | 0.0000 | 0.0000 |
| mixed | 48h_APF | 1 | 0.0000 | 0.0000 | 0.0000 | 0.0000 | 0.0000 | 0.0000 | 0.0000 | 0.0000 | 0.0000 | 0.0000 | 0.0000 | 0.0000 | 0.0000 | 0.0000 | 0.0000 | 0.0000 | 0.0000 | 0.0000 | 0.0000 | 0.0000 | 0.0000 |
| mixed | 48h_APF | 2 | 0.0000 | 0.0000 | 0.0000 | 0.0000 | 0.0000 | 0.0000 | 0.0000 | 0.0000 | 0.0000 | 0.0000 | 0.0000 | 0.0000 | 0.0000 | 0.0000 | 0.0000 | 0.0000 | 0.0000 | 0.0000 | 0.0000 | 0.0000 | 0.0000 |
| mixed | 48h_APF | 2 | 0.0000 | 0.0000 | 0.0000 | 0.0000 | 0.0000 | 0.0000 | 0.0000 | 0.0000 | 0.0000 | 0.0000 | 0.0000 | 0.0000 | 0.0000 | 0.0000 | 0.0000 | 0.0000 | 0.0000 | 0.0000 | 0.0000 | 0.0000 | 0.0000 |
| mixed | 48h_APF | 2 | 0.0000 | 0.0000 | 0.0000 | 0.0000 | 0.0000 | 0.0000 | 0.0000 | 0.0000 | 0.0000 | 0.0000 | 0.0000 | 0.0000 | 0.0000 | 0.0000 | 0.0000 | 0.0000 | 0.0000 | 0.0000 | 0.0000 | 0.0000 | 0.0000 |
| mixed | 48h_APF | 2 | 0.0000 | 0.0000 | 0.0000 | 0.0000 | 0.0000 | 0.0000 | 0.0000 | 0.0000 | 0.0000 | 0.0000 | 0.0000 | 0.0000 | 0.0000 | 0.0000 | 0.0000 | 0.0000 | 0.0000 | 0.0000 | 0.0000 | 0.0000 | 0.0000 |
