## Supplemental Table S2 for "Identification of a specialized lipid barrier for *Drosophila* metamorphosis"

| Repeat | Time_point | me-C26:0 | C27:1 | C27:0 | me-C28:0 | C29:1 | C29:0 | C30:1 | me-C30:0 | C31:1 | C30:0 | me-C32:0 | C33:2 | C33:1 | C35:2 | C37:2 |
|---|---|---|---|---|---|---|---|---|---|---|---|---|---|---|---|---|
| 1 | wL3 | 0.1928031 | 0.0362783 | 0.0045002 | 1.11641 | 0.1180382 | 0.0055172 | 0.0184495 | 0.8033963 | 0.2058571 | 0 | 0.0682474 | 0.3080098 | 0 | 0.0585762 | 0 |
| 1 | wL3 | 0.2375624 | 0.0418431 | 0.0052572 | 1.2394089 | 0.1130716 | 0.0054653 | 0.0170822 | 0.7714579 | 0.1915399 | 0 | 0.0689245 | 0.3911809 | 0 | 0.0914762 | 0 |
| 1 | wL3 | 0.1876003 | 0.0380133 | 0.0026438 | 1.0942774 | 0.1290953 | 0.0089583 | 0.0169031 | 0.7557729 | 0.2403349 | 0 | 0.0723043 | 0.3425055 | 0 | 0.0574399 | 0 |
| 1 | wL3 | 0.2337252 | 0.0314363 | 0.0050997 | 1.4963919 | 0.13709 | 0.0062024 | 0.0208498 | 0.9827512 | 0.2699203 | 0 | 0.0754035 | 0.2954046 | 0 | 0.1324165 | 0 |
| 1 | wL3 | 0.1418511 | 0.0283191 | 0.0030041 | 0.8148544 | 0.0893865 | 0.004513 | 0.0107704 | 0.6037664 | 0.1645953 | 0 | 0.0478651 | 0.2304462 | 0 | 0.060852 | 0 |
| 1 | wL3 | 0.1882749 | 0.0277893 | 0.0048684 | 0.7932682 | 0.0931375 | 0.0067033 | 0.0141432 | 0.6181582 | 0.1739689 | 0 | 0.0549233 | 0.2529034 | 0 | 0.0773097 | 0 |
| 1 | 0h_AE | 0.0433093 | 0.0097186 | 0.0007467 | 0.0864219 | 0.0335391 | 0.0008553 | 0.0066319 | 0.0398855 | 0.0262702 | 0.0004676 | 0.0059399 | 0.0447536 | 0.0069091 | 0.0273616 | 0 |
| 1 | 0h_AE | 0.0328694 | 0.0083283 | 0.001171 | 0.063107 | 0.0337852 | 0.0008401 | 0.0057314 | 0.0290566 | 0.0193956 | 0.0003488 | 0.0040759 | 0.0348832 | 0.0065506 | 0.0233917 | 0 |
| 1 | 0h_AE | 0.0341596 | 0.0099422 | 0.0011273 | 0.0589986 | 0.0240972 | 0.0003638 | 0.0037898 | 0.0271106 | 0.0146525 | 0.000778 | 0.003422 | 0.0373897 | 0.0052246 | 0.0207275 | 0 |
| 1 | 0h_AE | 0.024207 | 0.0064889 | 0.0010644 | 0.0475483 | 0.0192542 | 0.0004091 | 0.0031401 | 0.0213381 | 0.0115538 | 0.0003111 | 0.0027387 | 0.0283847 | 0.0047571 | 0.019455 | 0 |
| 1 | 0h_AE | 0.0246293 | 0.0065663 | 0.0009314 | 0.045356 | 0.021137 | 0.000312 | 0.0036107 | 0.0236373 | 0.0130614 | 0.0004235 | 0.0037811 | 0.0344546 | 0.0046777 | 0.0248647 | 0 |
| 2 | wL3 | 0.2174937 | 0.0606458 | 0.0084627 | 0.6841979 | 0.0826585 | 0.0056018 | 0.0101492 | 0.4303985 | 0.1032963 | 0.0026804 | 0.1769738 | 0.0336524 | 0 | 0.0484834 | 0.0160769 |
| 2 | wL3 | 0.1702502 | 0.0474154 | 0.004894 | 0.8193372 | 0.0974936 | 0.0078988 | 0.0120552 | 0.4637974 | 0.1210606 | 0.0022908 | 0.1721229 | 0.0414356 | 0 | 0.0398875 | 0.0090073 |
| 2 | wL3 | 0.1780779 | 0.0478226 | 0.0097639 | 0.7033266 | 0.0805509 | 0.0062637 | 0.0100989 | 0.3912764 | 0.1206579 | 0.0023661 | 0.1549955 | 0.0311856 | 0 | 0.0381768 | 0.0091585 |
| 2 | wL3 | 0.1464224 | 0.0439539 | 0.0059771 | 0.542642 | 0.0666441 | 0.0052223 | 0.0058842 | 0.3310501 | 0.0854748 | 0.0014436 | 0.1534856 | 0.0230926 | 0 | 0.0370134 | 0.0068696 |
| 2 | wL3 | 0.1954483 | 0.0572701 | 0.0072682 | 0.7627822 | 0.0868003 | 0.0068699 | 0.0098766 | 0.3465832 | 0.0883166 | 0.0012323 | 0.1349882 | 0.0210522 | 0 | 0.0289519 | 0.0076179 |
| 2 | wL3 | 0.1556819 | 0.0490855 | 0.0052456 | 0.5885703 | 0.0653455 | 0.0041579 | 0.0076181 | 0.3418384 | 0.0966832 | 0.0010181 | 0.1479338 | 0.0313765 | 0 | 0.0297325 | 0.0104387 |
| 2 | wL3 | 0.1611108 | 0.0447972 | 0.0059021 | 0.4824659 | 0.0524955 | 0.0045688 | 0.0051663 | 0.2949098 | 0.0639614 | 0.0016197 | 0.1393801 | 0.0290167 | 0 | 0.026502 | 0.0083439 |
| 2 | wL3 | 0.1468982 | 0.0329118 | 0.0050787 | 0.5643395 | 0.0680596 | 0.0054159 | 0.0072041 | 0.3405635 | 0.1041176 | 0.0019608 | 0.1389686 | 0.0229459 | 0 | 0.0401334 | 0.0072418 |
| 2 | 0h_AE | 0.1881605 | 0.0440512 | 0.0060823 | 0.3495214 | 0.077558 | 0.0031748 | 0.0137818 | 0.1372081 | 0.0746692 | 0.0026978 | 0.1448911 | 0.0165296 | 0.0280985 | 0.1015942 | 0.0331895 |
| 2 | 0h_AE | 0.1773988 | 0.047611 | 0.0048986 | 0.2809353 | 0.0853885 | 0.003335 | 0.0226087 | 0.1318021 | 0.0694321 | 0.0021083 | 0.1396952 | 0.0155596 | 0.0311498 | 0.0872749 | 0.0329458 |
| 2 | 0h_AE | 0.1712023 | 0.0385404 | 0.0056442 | 0.3123695 | 0.0752206 | 0.003379 | 0.0144177 | 0.1160659 | 0.0820172 | 0.0015138 | 0.1645262 | 0.0122408 | 0.0263456 | 0.0665662 | 0.0248815 |
| 2 | 0h_AE | 0.1286763 | 0.0329068 | 0.0039238 | 0.2720749 | 0.0864473 | 0.0027476 | 0.0111647 | 0.1118583 | 0.0687742 | 0.0016424 | 0.1469948 | 0.0130024 | 0.0192075 | 0.0704779 | 0.0161936 |
| 3 | wL3 | 0.0848645 | 0.076315 | 0 | 1.5298398 | 0.1576062 | 0.0047009 | 0.0222441 | 0.2560276 | 0.2249274 | 0.0769669 | 0.2902013 | 0 | 0.0818199 | 0 | 0 |
| 3 | wL3 | 0.0664914 | 0.0480654 | 0 | 1.1306963 | 0.1369535 | 0.0039368 | 0.0162746 | 0.220309 | 0.1682055 | 0.0710083 | 0.2318189 | 0 | 0.0792404 | 0 | 0 |
| 3 | wL3 | 0.07042 | 0.0631662 | 0 | 1.3895156 | 0.1569693 | 0.0027404 | 0.0200673 | 0.2068724 | 0.1819102 | 0.07068 | 0.2243229 | 0 | 0.0835127 | 0 | 0 |
| 3 | wL3 | 0.018888 | 0.0407016 | 0 | 0.1001273 | 0.0192816 | 0 | 0 | 0.0210297 | 0 | 0 | 0 | 0 | 0 | 0 | 0 |
| 3 | 0h_AE | 0.0102974 | 0.0091058 | 0 | 0.4896275 | 0.0511091 | 0.0025713 | 0.0127605 | 0.0910413 | 0.1248898 | 0.0204722 | 0.2125588 | 0.0507734 | 0.0887947 | 0.0162453 | 0 |
| 3 | 0h_AE | 0.0065642 | 0.006782 | 0 | 0.405415 | 0.0500331 | 0.002755 | 0.0104102 | 0.0770707 | 0.1126172 | 0.0185332 | 0.1773061 | 0.0426038 | 0.0689388 | 0.012034 | 0 |
| 3 | 0h_AE | 0.0105433 | 0.0080467 | 0 | 0.5386722 | 0.0661623 | 0.0016896 | 0.0171774 | 0.097091 | 0.1120962 | 0.0229568 | 0.2242042 | 0.0543888 | 0.0867087 | 0.0160084 | 0 |
| 3 | 0h_AE | 0.005768 | 0.0068653 | 0 | 0.349441 | 0.047136 | 0.0019164 | 0.0122779 | 0.0680146 | 0.0998536 | 0.0157681 | 0.1745711 | 0.0387063 | 0.0661474 | 0.0105158 | 0 |
