## Supplemental Table S3 for "Identification of a specialized lipid barrier for *Drosophila* metamorphosis"

| Tissue | Repeat | me-C26:0 | C27:1 | me-C28:0 | C29:1 | C30:1 | me-C30:0 | C31:1 | me-C32:0 | C33:2 | C35:2 | C37:2 |
|---|---|---|---|---|---|---|---|---|---|---|---|---|
| Carcass | 1 | 0.00992176 | 0 | 0.07727179 | 0 | 0 | 0.05402681 | 0 | 0.00684661 | 0 | 0 | 0 |
| Carcass | 1 | 0.00888441 | 0 | 0.06741822 | 0 | 0 | 0.05307108 | 0 | 0.00853896 | 0 | 0 | 0 |
| Carcass | 1 | 0.00775168 | 0 | 0.0752758 | 0 | 0 | 0.0581839 | 0 | 0.0077798 | 0 | 0 | 0 |
| Fatbody | 1 | 0.15718009 | 0.0146401 | 0.66032858 | 0.0588748 | 0.00819402 | 0.30805841 | 0.09874681 | 0.02431733 | 0.15610112 | 0.04030494 | 0.01159147 |
| Fatbody | 1 | 0.24013259 | 0.02781366 | 0.77842819 | 0.07402476 | 0.00803667 | 0.42433515 | 0.12470916 | 0.03178894 | 0.19939794 | 0.05235334 | 0.01882799 |
| Fatbody | 1 | 0.2049617 | 0.02508032 | 0.81760762 | 0.07609273 | 0.01203637 | 0.43701547 | 0.13506343 | 0.02855681 | 0.19775399 | 0.04663865 | 0.01357503 |
| Rest | 1 | 0.00747118 | 0 | 0.01179537 | 0 | 0 | 0.0031419 | 0 | 0 | 0 | 0 | 0 |
| Rest | 1 | 0.01327569 | 0 | 0.02894715 | 0 | 0 | 0.0187496 | 0 | 0 | 0 | 0 | 0 |
| Rest | 1 | 0.00538866 | 0 | 0.01630431 | 0 | 0 | 0.01040502 | 0 | 0 | 0 | 0 | 0 |
| Carcass | 2 | 0.01688371 | 0 | 0.1179189 | 0 | 0 | 0.08530225 | 0 | 0 | 0 | 0 | 0 |
| Carcass | 2 | 0.01310669 | 0 | 0.14297927 | 0 | 0 | 0.08928512 | 0 | 0.01789602 | 0 | 0 | 0 |
| Carcass | 2 | 0 | 0 | 0.17120437 | 0 | 0 | 0.09574546 | 0 | 0.01675688 | 0 | 0 | 0 |
| Carcass | 2 | 0.01491076 | 0 | 0.14376933 | 0 | 0 | 0.11082041 | 0 | 0.01170297 | 0 | 0 | 0 |
| Fatbody | 2 | 0.32331369 | 0.03899438 | 1.54697148 | 0.13692967 | 0.01583585 | 0.83305808 | 0.20956822 | 0.06344144 | 0.28721384 | 0.07061005 | 0.01898248 |
| Fatbody | 2 | 0.36446487 | 0.05792011 | 1.91068834 | 0.15954295 | 0.01162746 | 0.91357148 | 0.23020487 | 0.05379797 | 0.26768661 | 0.06927371 | 0.01302276 |
| Fatbody | 2 | 0.37982458 | 0.05973068 | 1.78577007 | 0.1720019 | 0.02015401 | 1.01256668 | 0.25484724 | 0.06939465 | 0.31622787 | 0.07100054 | 0.02202204 |
| Fatbody | 2 | 0.36462519 | 0.03545316 | 1.52287551 | 0.13813673 | 0.01471057 | 0.86834908 | 0.26382372 | 0.06408197 | 0.29766432 | 0.08303307 | 0.02613096 |
| Rest | 2 | 0.0195376 | 0 | 0.05282844 | 0 | 0 | 0.03941621 | 0 | 0 | 0 | 0 | 0 |
| Rest | 2 | 0.01237938 | 0 | 0.05360085 | 0 | 0 | 0.03566531 | 0 | 0 | 0 | 0 | 0 |
| Rest | 2 | 0.01838375 | 0 | 0.05774271 | 0 | 0 | 0.0333968 | 0 | 0 | 0 | 0 | 0 |
| Rest | 2 | 0.01531466 | 0 | 0.0450067 | 0 | 0 | 0.02179658 | 0 | 0 | 0 | 0 | 0 |
| Fat body | 3 | 0.23654369 | 0.04097661 | 0.39591596 | 0.05988706 | 0.00504637 | 0.07808938 | 0.03691485 | 0.00486038 | 0.07857671 | 0.01696709 | ND |
| Fat body | 3 | 0.37734329 | 0.07609335 | 0.58817415 | 0.09632843 | 0.00314464 | 0.10734566 | 0.05608573 | 0.00541608 | 0.10260384 | 0.02155425 | ND |
| Fat body | 3 | 0.05692369 | 0.01273939 | 0.10704878 | 0.01750591 | 0.0022504 | 0.02519872 | 0.01552124 | 0.00348861 | 0.03033019 | 0.01042394 | ND |
| Fat body | 3 | 0.39211275 | 0.07025868 | 0.90451867 | 0.13453662 | 0.00653474 | 0.1326194 | 0.06726955 | 0.00613137 | 0.1537773 | 0.03321025 | ND |
| Rest | 3 | 0.01293644 | 0 | 0.02940652 | 0 | 0 | 0 | 0 | 0 | 0 | 0 | ND |
| Rest | 3 | 0.01399194 | 0 | 0.01250798 | 0 | 0 | 0 | 0 | 0 | 0 | 0 | ND |
| Rest | 3 | 0.0201328 | 0 | 0.03105576 | 0 | 0 | 0 | 0 | 0 | 0 | 0 | ND |
| Rest | 3 | 0.06001079 | 0 | 0.04785787 | 0 | 0 | 0 | 0 | 0 | 0 | 0 | ND |
| Carcass | 3 | 0 | 0 | 0.05491658 | 0 | 0 | 0 | 0 | 0 | 0 | 0 | ND |
| Carcass | 3 | 0 | 0 | 0 | 0 | 0 | 0 | 0 | 0 | 0 | 0 | ND |
| Carcass | 3 | 0 | 0 | 0 | 0 | 0 | 0 | 0 | 0 | 0 | 0 | ND |
| Carcass | 3 | 0 | 0 | 0 | 0 | 0 | 0 | 0 | 0 | 0 | 0 | ND |
