## Supplemental Table S4 for "Identification of a specialized lipid barrier for *Drosophila* metamorphosis"

| Figure | Comparison | Statistical test | p-value |
|---|---|---|---|
| Figure 2E | me-C28:0: w - Cyp4g1 | linear mixed effect model; Bonferroni correction | 0.0269 |
| Figure 2E | me-C28:0: GFP [i] - Cyp4g1 | linear mixed effect model; Bonferroni correction | 0.008 |
| Figure 2E | C31:1: w - Cyp4g1 | linear mixed effect model; Bonferroni correction | 0.0032 |
| Figure 2E | C31:1: GFP [i] - Cyp4g1 | linear mixed effect model; Bonferroni correction | 0.0034 |
| Figure 2E | C33:2: w - Cyp4g1 | linear mixed effect model; Bonferroni correction | 0.0006 |
| Figure 2E | C33:2: GFP [i] - Cyp4g1 | linear mixed effect model; Bonferroni correction | 0.0001 |
| Figure 2F | L:29C: Cyp4g1 [i] | one sample t-test; mean = 100 | 2.20E-16 |
| Figure 2F | L:18C: Cyp4g1 [i] | one sample t-test; mean = 100 | 0.0653 |
| Figure 3C | CG6660 ; female: wL3 - adult | Anova; Tukey post hoc test | 0.0002366 |
| Figure 3C | CG6660 ; male: wL3 - adult | Anova; Tukey post hoc test | 0.0029949 |
| Figure 3C | CG8435 ; female: wL3 - adult | Anova; Tukey post hoc test | 0.0237918 |
| Figure 3C | CG8435 ; male: wL3 - adult | Anova; Tukey post hoc test | 0.0778432 |
| Figure 3C | EloF ; female: wL3 - adult | Anova; Tukey post hoc test | 7.36E-05 |
| Figure 3C | EloF ; male: wL3 - adult | Anova; Tukey post hoc test | 0.46 |
| Figure 3D | w | one sample t-test; mean = 100 | 0.103 |
| Figure 3D | GFP[i] | one sample t-test; mean = 100 | 0.02231 |
| Figure 3D | Cyp4g1 [i] | one sample t-test; mean = 100 | 0.1632 |
| Figure 3D | CG8534 [i] | one sample t-test; mean = 100 | 0.1563 |
| Figure 3D | CG6660 [i] | one sample t-test; mean = 100 | < 2.2e-16 |
| Figure 4D | 7-C23:1 :EloF [i] vs.GFP [i] | linear model; Bonferroni correction | 2.98E-06 |
| Figure 4D | 7-C25:1 :EloF [i] vs.GFP [i] | linear model; Bonferroni correction | 1.94E-04 |
| Figure 4D | 7-C27:1 :EloF [i] vs.GFP [i] | linear model; Bonferroni correction | 0.10397788 |
| Figure 4D | 9-C25:1 :EloF [i] vs.GFP [i] | linear model; Bonferroni correction | 0.094720873 |
| Figure 4D | 9-C27:1 :EloF [i] vs.GFP [i] | linear model; Bonferroni correction | 2.21E-07 |
| Figure 4D | C23:0 :EloF [i] vs.GFP [i] | linear model; Bonferroni correction | 1.25E-15 |
| Figure 4D | C25:0 :EloF [i] vs.GFP [i] | linear model; Bonferroni correction | 0.336277911 |
| Figure 4D | C25:2 :EloF [i] vs.GFP [i] | linear model; Bonferroni correction | 6.16E-13 |
| Figure 4D | C27:0 :EloF [i] vs.GFP [i] | linear model; Bonferroni correction | 0.201145364 |
| Figure 4D | C27:2 :EloF [i] vs.GFP [i] | linear model; Bonferroni correction | 0.063056345 |
| Figure 4D | C29:2 :EloF [i] vs.GFP [i] | linear model; Bonferroni correction | 6.77E-04 |
| Figure 4D | me-C22:0 :EloF [i] vs.GFP [i] | linear model; Bonferroni correction | 0.304632129 |
| Figure 4D | me-C24:0 :EloF [i] vs.GFP [i] | linear model; Bonferroni correction | < 2.2e-16 |
| Figure 4D | me-C26:0 :EloF [i] vs.GFP [i] | linear model; Bonferroni correction | 0.065717913 |
| Figure 4D | me-C28:0 :EloF [i] vs.GFP [i] | linear model; Bonferroni correction | < 2.2e-16 |
| Figure 4D | 7-C23:1 :Df/+ vs. EloHL/Df | linear model; Bonferroni correction | 2.32E-04 |
| Figure 4D | 7-C23:1 :Df/+ vs. EloHL/+ | linear model; Bonferroni correction | 0.146115315 |
| Figure 4D | 7-C23:1 :EloHL/+ vs. EloHL/Df | linear model; Bonferroni correction | 1.64E-08 |
| Figure 4D | 7-C25:1 :Df/+ vs. EloHL/Df | linear model; Bonferroni correction | 1 |
| Figure 4D | 7-C25:1 :Df/+ vs. EloHL/+ | linear model; Bonferroni correction | 0.103395972 |
| Figure 4D | 7-C25:1 :EloHL/+ vs. EloHL/Df | linear model; Bonferroni correction | 0.14574403 |
| Figure 4D | 7-C27:1 :Df/+ vs. EloHL/Df | linear model; Bonferroni correction | 1 |
| Figure 4D | 7-C27:1 :Df/+ vs. EloHL/+ | linear model; Bonferroni correction | 1 |
| Figure 4D | 7-C27:1 :EloHL/+ vs. EloHL/Df | linear model; Bonferroni correction | 0.813235567 |
| Figure 4D | 9-C25:1 :Df/+ vs. EloHL/Df | linear model; Bonferroni correction | 0.595820247 |
| Figure 4D | 9-C25:1 :Df/+ vs. EloHL/+ | linear model; Bonferroni correction | 0.157580705 |
| Figure 4D | 9-C25:1 :EloHL/+ vs. EloHL/Df | linear model; Bonferroni correction | 0.004012754 |
| Figure 4D | 9-C27:1 :Df/+ vs. EloHL/Df | linear model; Bonferroni correction | 0.27949015 |
| Figure 4D | 9-C27:1 :Df/+ vs. EloHL/+ | linear model; Bonferroni correction | 1 |
| Figure 4D | 9-C27:1 :EloHL/+ vs. EloHL/Df | linear model; Bonferroni correction | 0.075103845 |
| Figure 4D | C23:0 :Df/+ vs. EloHL/Df | linear model; Bonferroni correction | 1 |
| Figure 4D | C23:0 :Df/+ vs. EloHL/+ | linear model; Bonferroni correction | 0.056328057 |
| Figure 4D | C23:0 :EloHL/+ vs. EloHL/Df | linear model; Bonferroni correction | 0.125810445 |
| Figure 4D | C25:0 :Df/+ vs. EloHL/Df | linear model; Bonferroni correction | 0.110536899 |
| Figure 4D | C25:0 :Df/+ vs. EloHL/+ | linear model; Bonferroni correction | 0.484826998 |
| Figure 4D | C25:0 :EloHL/+ vs. EloHL/Df | linear model; Bonferroni correction | 1 |
| Figure 4D | C25:2 :Df/+ vs. EloHL/Df | linear model; Bonferroni correction | < 2.2e-16 |
| Figure 4D | C25:2 :Df/+ vs. EloHL/+ | linear model; Bonferroni correction | 1 |
| Figure 4D | C25:2 :EloHL/+ vs. EloHL/Df | linear model; Bonferroni correction | 1.21E-35 |
| Figure 4D | C27:0 :Df/+ vs. EloHL/Df | linear model; Bonferroni correction | 0.869902492 |
| Figure 4D | C27:0 :Df/+ vs. EloHL/+ | linear model; Bonferroni correction | 1 |
| Figure 4D | C27:0 :EloHL/+ vs. EloHL/Df | linear model; Bonferroni correction | 1 |

| | | | |
|---|---|---|---|
| Figure 4D | C27:2 :Df/+ vs. EloHL/Df | linear model; Bonferroni correction | 1.76E-05 |
| Figure 4D | C27:2 :Df/+ vs. EloHL/+ | linear model; Bonferroni correction | 0.018138761 |
| Figure 4D | C27:2 :EloHL/+ vs. EloHL/Df | linear model; Bonferroni correction | 3.74E-12 |
| Figure 4D | C29:2 :Df/+ vs. EloHL/Df | linear model; Bonferroni correction | 0.007963987 |
| Figure 4D | C29:2 :Df/+ vs. EloHL/+ | linear model; Bonferroni correction | 0.019132954 |
| Figure 4D | C29:2 :EloHL/+ vs. EloHL/Df | linear model; Bonferroni correction | 5.13E-08 |
| Figure 4D | me-C22:0 :Df/+ vs. EloHL/Df | linear model; Bonferroni correction | 1 |
| Figure 4D | me-C22:0 :Df/+ vs. EloHL/+ | linear model; Bonferroni correction | 0.070822226 |
| Figure 4D | me-C22:0 :EloHL/+ vs. EloHL/Df | linear model; Bonferroni correction | 0.007096245 |
| Figure 4D | me-C24:0 :Df/+ vs. EloHL/Df | linear model; Bonferroni correction | < 2.2e-16 |
| Figure 4D | me-C24:0 :Df/+ vs. EloHL/+ | linear model; Bonferroni correction | 0.766792529 |
| Figure 4D | me-C24:0 :EloHL/+ vs. EloHL/Df | linear model; Bonferroni correction | < 2.2e-16 |
| Figure 4D | me-C26:0 :Df/+ vs. EloHL/Df | linear model; Bonferroni correction | 1 |
| Figure 4D | me-C26:0 :Df/+ vs. EloHL/+ | linear model; Bonferroni correction | 1 |
| Figure 4D | me-C26:0 :EloHL/+ vs. EloHL/Df | linear model; Bonferroni correction | 1 |
| Figure 4D | me-C28:0 :Df/+ vs. EloHL/Df | linear model; Bonferroni correction | 0.589909476 |
| Figure 4D | me-C28:0 :Df/+ vs. EloHL/+ | linear model; Bonferroni correction | 1 |
| Figure 4D | me-C28:0 :EloHL/+ vs. EloHL/Df | linear model; Bonferroni correction | 0.2165722 |
| Figure 5B | Controls - Mutant: 0h | Anova; Tukey post hoc test | 0.2301865 |
| Figure 5B | Controls - Mutant: 24h | Anova; Tukey post hoc test | 0.001293 |
| Figure 5B | Controls - Mutant: 48h | Anova; Tukey post hoc test | 0.0000024 |
| Figure 5B | Controls - Mutant: 72h | Anova; Tukey post hoc test | 0.0031812 |
| Figure 5C | Controls - Cyp4g1 [i]: 0h | Anova; Tukey post hoc test | 0.9876937 |
| Figure 5C | Controls - Cyp4g1 [i]: 24h | Anova; Tukey post hoc test | < 2.2e-16 |
| Figure 5C | Controls - Cyp4g1 [i]: 48h | Anova; Tukey post hoc test | < 2.2e-16 |
| Figure 5C | Controls - Cyp4g1 [i]: 72h | Anova; Tukey post hoc test | < 2.2e-16 |
| Figure 5F | internal pupal vs external pupal cuticle | Anova; Tukey post hoc test | 0.0002679 |
