## Supplemental Table S5 for "Identification of a specialized lipid barrier for *Drosophila* metamorphosis"

| Models | $\omega$ | lnL | np | Models compared | $2\Delta(\ln L)$ | P-value |
|---|---|---|---|---|---|---|
| A: All branches have the same $\omega$ | $\omega = 0.16605$ | -8157.765611 | 52 | | | |
| B: All branches in EloF clade have $\omega 1$ and other branches have $\omega 0$ | $\omega 0 = 0.14904$, $\omega 1 = 0.24708$ | -8153.820019 | 53 | B vs. A | 7.891184 | 0.004967633 |
| C: All branches in EloHL clade have $\omega 1$ and other branches have $\omega 0$ | $\omega 0 = 0.15862$, $\omega 1 = 0.21707$ | -8156.564574 | 53 | C vs. A | 2.402074 | 0.121174504 |
| D: All branches in EloHL clade have $\omega 1$ and those in eloF clade have $\omega 2$; other branches have $\omega 0$ | $\omega 0 = 0.13807$, $\omega 1 = 0.20799$, $\omega 2 = 0.24122$ | -8151.824032 | 54 | D vs. A | 11.883158 | 0.002627877 |
| | | | | D vs. B | 3.991974 | 0.045717474 |
| | | | | D vs. C | 9.481084 | 0.002076013 |
