## Supplemental Table S6 for "Identification of a specialized lipid barrier for *Drosophila* metamorphosis"

| Gene name | Primer name | Primer sequence | Assay |
|---|---|---|---|
| CG31522 | CG31522_fw | AGGCGTTCGTCTGGTGGATT | Gene expression |
| CG31523 | CG31523_fw | AGAGGGTGACGCCCTGATTG | Gene expression |
| CG2781 | CG2781_fw | ACGCGCATTTAGTACCTGTTGAA | Gene expression |
| CG33110 | CG33110_fw | ACAACAGCAGCTCGCGATTC | Gene expression |
| CG32072 | CG32072_fw | ATGCCAACGGGTTCGACCTA | Gene expression |
| CG6921 | CG6921_fw | TGAGCTCACCTATGCCCGTG | Gene expression |
| CG6660 | CG6660_fw | TGGATCTGCTCGACACGGTAT | Gene expression |
| CG8534 | CG8534_fw | CGCTGACCATCGTTTCGCTC | Gene expression |
| CG18609 | CG18609_fw | CGCATACCTCAAGCGGATCCTA | Gene expression |
| CG16904 | CG16904_fw | AAGCTGGGCAGGAAGCTGAT | Gene expression |
| CG9458 | CG9458_fw | GCGGGTCCAAAGATCATGCG | Gene expression |
| CG9459 | CG9459_fw | GGTTTATGCAGAATCAAAAGCCC | Gene expression |
| CG16905 | CG16905_fw | GCACATTGATTGGCTATCTGCT | Gene expression |
| CG30008 | CG30008_fw | ATGGAGGCGTCAGCAAGTAT | Gene expression |
| CG3971 | CG3971_fw | TCTGCGTGCCCAGCTACATC | Gene expression |
| CG31522 | CG31522_rv | CGTCATGGGCGTTGATGCTC | Gene expression |
| CG31523 | CG31523_rv | GCCCTCCTTGTAGGCACCAT | Gene expression |
| CG2781 | CG2781_rv | CGTGTTCGCGGATCCGATTT | Gene expression |
| CG33110 | CG33110_rv | TAGACGGCCACCATGCCAAA | Gene expression |
| CG32072 | CG32072_rv | GAGGACCCAGCACACTCAGG | Gene expression |
| CG6921 | CG6921_rv | TCTCTTGGATGGACGTGCGG | Gene expression |
| CG6660 | CG6660_rv | TGCAGTGTGATCCACCCAGG | Gene expression |
| CG8534 | CG8534_rv | CGAACAAAAAGTGAAGACCCGCT | Gene expression |
| CG18609 | CG18609_rv | TGCAGATTGCAAATGCCCACG | Gene expression |
| CG16904 | CG16904_rv | AACAGATGGATGCCCCACACA | Gene expression |
| CG9458 | CG9458_rv | CCAGCATGAAGTGGACAGCAAAA | Gene expression |
| CG9459 | CG9459_rv | CCAGGACCAAGCATGAAGTG | Gene expression |
| CG16905 | CG16905_rv | GATTTGGTAGGCTTTCAGGACA | Gene expression |
| CG30008 | CG30008_rv | GAGCTCGTAAGGTTTGCGCC | Gene expression |
| CG3971 | CG3971_rv | CTGCTTGCGGAGCACAATGA | Gene expression |
| CG8534 | Homology_1_fw | AGCTCAGCTAGCCCACTCGAGGTAATGTTTCACTG | Mutant generation |
| CG8534 | Homology_1_rv | TATCGAGCGGCCGCCAGTGAAACATTACCTCGAGTGGCGCATCTGATGCACCTGTCCGC | Mutant generation |
| CG8534 | Homology_2_fw | AATCAGACTAGTCCAAGTTACTCGCTAGTCTTTTA | Mutant generation |
| CG8534 | Homology_2_rv | GTGGAGCAGAATAATGGCGTAAAAGACTAGCGAGTAACTTGGACCGGTTAGCCG | Mutant generation |
| CG8534 | CFD4_T1_CG8534 | TATATATAGGAAAGATATCCGGGTGAACTTCGAGTGAAACATTACCTCGAGGTTTTAGAGCTAGAAATAGCAAG | Mutant generation |
| CG8534 | CFD4_T2_CG8534 | GACCTATTTTCAATTTAACGTCGAAAAGACTAGCGAGTAACTGTTTTAGAGCTAGAAATAGCAAGTTAAAAT | Mutant generation |
| outside CG8534 (5') | Primer 1 | GCCACACATTTGTGCAAGACAAAGG | Mutant integration check |
| mCherry | Primer 2 | CTCCATGTGCACCTTGAAGCGC | Mutant integration check |
| CG8534 | Primer 3 | CGCCTTGCCACTCATCGGAAG | Gene check |
| outside CG8534 (3') | Primer 4 | CATGGCGAATCTAATCGAGTTGTGCG | Gene check |
